# The dormancy-specific regulator, SutA, is intrinsically disordered and modulates transcription initiation in *Pseudomonas aeruginosa*

**DOI:** 10.1101/423384

**Authors:** Megan Bergkessel, Brett M. Babin, David G. VanderVelde, Michael J. Sweredoski, Annie Moradian, Roxana Eggleston-Rangel, Sonja Hess, David A. Tirrell, Irina Artsimovitch, Dianne K. Newman

## Abstract

Though bacteria in nature are often nutritionally limited and growing slowly, most of our understanding of core cellular processes such as transcription comes from studies in a handful of model organisms doubling rapidly under nutrient-replete conditions. We previously identified a small protein of unknown function, called SutA, in a global screen of proteins synthesized in *Pseudomonas aeruginosa* under growth arrest (Babin BM*, et al.* (2016) SutA is a bacterial transcription factor expressed during slow growth in *Pseudomonas aeruginosa*. *PNAS* 113(5):E597-605). SutA binds RNA polymerase (RNAP), causing widespread changes in gene expression, including upregulation of the ribosomal RNA (rRNA) genes. Here, using biochemical and structural methods, we examine how SutA interacts with RNAP and the functional consequences of these interactions. We show that SutA consists of a central α-helix with unstructured N- and C-terminal tails, and binds to the β1 domain of RNAP. It activates transcription from the *P. aeruginosa rrn* promoter by both the housekeeping sigma factor holoenzyme (Eσ^70^) and the general stress response sigma factor holoenzyme (Eσ^S^) *in vitro,* and its N-terminal tail is required for activation in both holoenzyme contexts. However, we find that the interaction between SutA and each holoenzyme is distinct, with the SutA C-terminal tail and an acidic loop unique to σ^70^ playing the determining roles in these differences. Our results add SutA to a growing list of transcription regulators that use their intrinsically disordered regions to remodel transcription complexes.

**SIGNIFICANCE:** Little is known about how bacteria regulate their activities during periods of dormancy, yet growth arrest dominates bacterial existence in most environments and is directly relevant to the problem of physiological antibiotic tolerance. Though much is known about transcription in the model organism, *Escherichia coli*, even there, our understanding of gene expression during dormancy is incomplete. Here we explore how transcription under growth arrest is modulated in *Pseudomonas aeruginosa* by the small acidic protein, SutA. We show that SutA binds to RNA polymerase and controls transcription by a mechanism that is distinct from other known regulators. Our work underscores the potential for fundamental, mechanistic discovery in this important and understudied realm of bacterial physiology.

## INTRODUCTION

Despite the fact that most natural environments do not allow bacteria to double every 20-30 minutes, our understanding of essential cellular processes—such as DNA replication, transcription, and translation—has been shaped by studies of a few model organisms growing exponentially at these rates, or responding to a rapid shift from exponential to slow growth. We do not know how the molecular machines responsible for transcription and translation (processes that are necessary to maintain homeostasis even when cell division is not occurring) adapt to long periods of reduced activity and low or uneven substrate availability (2). *P. aeruginosa* and many other members of the Pseudomonadales order are notable opportunists, capable of using diverse substrates for rapid growth but also able to persist in dormancy for long periods (3), making them attractive model systems for addressing such questions. A better understanding of slow-growing or dormant states in *P. aeruginosa* also has clinical importance, as these states are thought to contribute to this organism’s antibiotic tolerance in chronic infections (4–6).

Accordingly, in previous work, we used a proteomics-based screen to identify *P. aeruginosa* regulators that are preferentially expressed during hypoxia-induced growth arrest. We identified an RNAP-binding protein, SutA, that had broad impacts on gene expression and affected that affected the ability of *P. aeruginosa* to form biofilms and produce virulence factors. Notably, SutA expression led to upregulation of the rRNA genes under slow-growth conditions, and ChIP data showed both that SutA localized to rRNA promoters and that higher levels of RNAP localized to rRNA promoters when SutA was present (1). In this study, we investigate whether, as these results suggest, SutA directly impacts transcription initiation at the *rrn* promoters, and how these effects are carried out.

The regulation the *rrn* promoters in *E. coli* is one of the best-studied examples of growth-rate-responsive control of bacterial gene expression. While they can drive extremely high levels of expression (up to about 70% of all transcription) during exponential growth, they are rapidly and strongly repressed entry into stationary phase (7). This behavior depends on an extremely unstable open complex (OC) formed at *rrn* P1, which facilitates high levels of expression by enabling rapid promoter clearance by RNAP but also sensitizes initiation to conditions encountered during nutrient downshifts, such as decreased concentrations of the initiating nucleotides ([iNTPs]) (8). A second *rrn* promoter that drives low levels of expression and is relatively insensitive to regulatory inputs, P2, has been proposed as the mechanism by which some rRNA transcription can be maintained during stationary phase (9, 10), but this would imply that expression levels in *E. coli* are not actively modulated during protracted dormancy.

Expression of *rrn* is further modulated by diverse regulators acting at different stages of transcription initiation in different organisms. In many cases, the unstable OC is the target of additional regulation. For example, in *E. coli*, the signaling molecule (p)ppGpp and its co-regulator DksA bind to RNAP during early stationary phase and further destabilize the final *rrn* OC (11); the identities of the iNTPs (adenosine or guanosine) allow for direct coordination with the diminished energy stores available to drive translation (7). Also, in many clades outside the Gammaproteobacteria, homologs of the global regulator CarD can enhance rRNA expression by directly stabilizing the OC (12). By contrast, some factors that activate *rrn* P1 during rapid exponential growth in *E. coli* (*e.g.,* Fis and DNA supercoiling) exert their effects before the final OC has formed, by helping to recruit RNAP or facilitating the initial opening of the double-stranded DNA (7, 13).

SutA lacks sequence or structural homology to any known transcription factor, raising the possibility that its mode of action is unique. Here, we report that SutA binds to a site on RNAP that is distinct from the binding sites of other regulators, that its activation of *rrn* transcription depends on its intrinsically disordered N- and C-terminal tails, and that its activity is modulated by the identity of the σ factor. Though our work focuses on a specific transcription factor and promoter in *P. aeruginosa*, the topic it tackles and the questions it raises are broadly relevant to understanding how bacteria survive periods of slow growth or dormancy in diverse environments.

## RESULTS

### SutA consists of a conserved alpha helix flanked by flexible N- and C-terminal tails

Because SutA is a small protein (105 amino acids) with no similarity to any known domains, we first explored its structural characteristics. We began by looking at structure predictions (using the Jpred4 algorithm for secondary structure and DISOPRED3 for intrinsic disorder) and sequence conservation (14–16). SutA homologs are found in most organisms in the “Pseudomonadales-Oceanospirallales” clade of Gammaproteobacteria (17). Residues 56-76 are predicted to form an α-helix, followed by a β-strand comprising residues 81-84, but the rest of the protein has no predicted secondary structural elements, and residues 1-50 and 101-105 are predicted to be intrinsically disordered (Figure 1A). While the central, potentially structured region is reasonably well conserved, some homologs completely lack the last 15-18 residues, and others lack most or all of the first 40 residues (Figure S1). This suggests that the N-and C-terminal tails (N-tail and C-tail) might function independently and that their removal might not affect folding/function of other regions.

**Figure 1.**
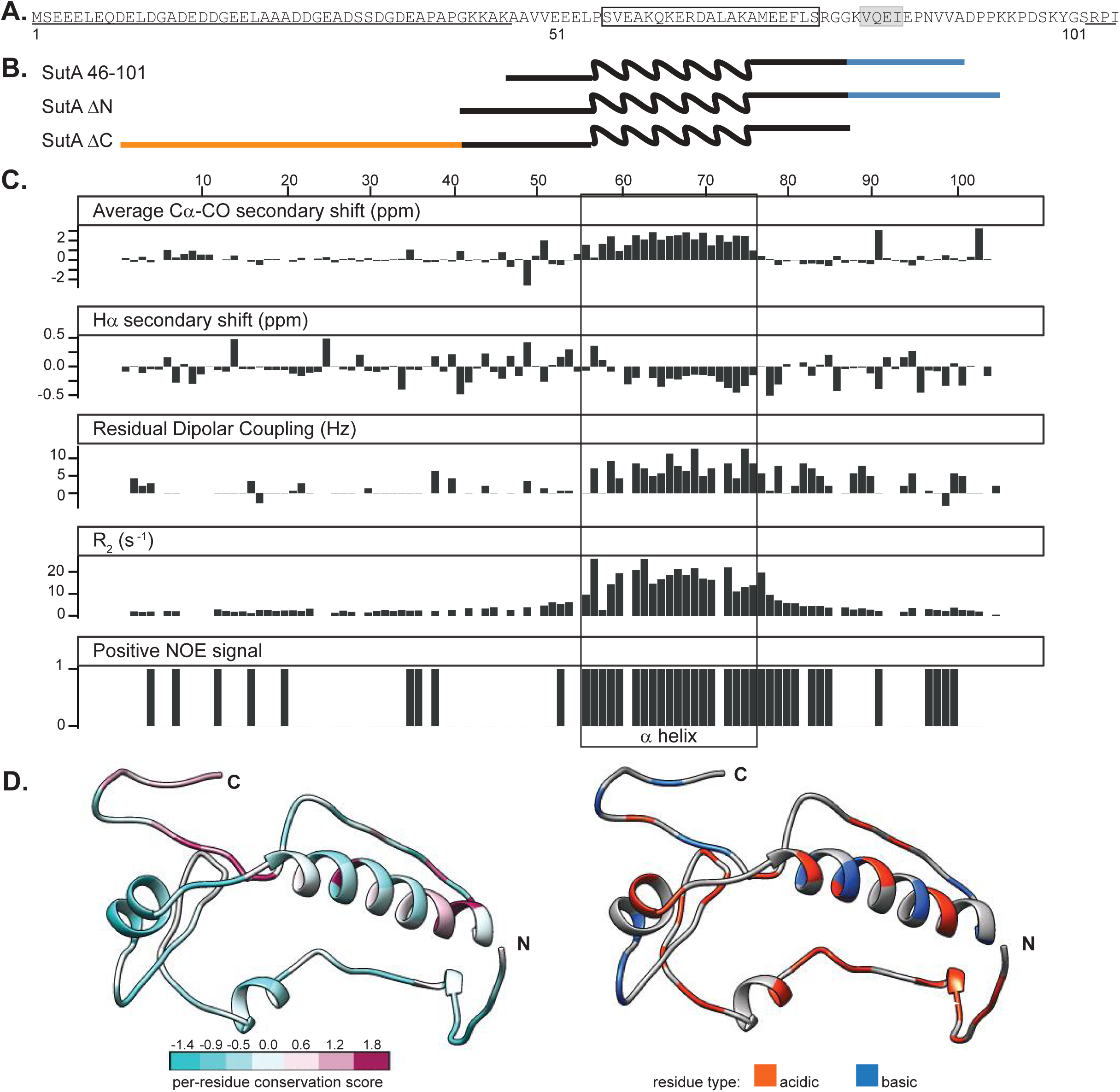
NMR data confirm presence of alpha helix from aa 56-76 and flexible N- and C-terminal tails. **A.** Primary amino acid sequence for SutA, with computational predictions indicated: underlining = intrinsic disorder; boxing = α-helix; gray shading = β-strand. **B.** Schematic of constructs used; wavy line = α-helix region; blue = C-tail; orange = N-tail. Schematics are aligned with residue numbers and NMR data in (C). **C.** Secondary chemical shift indices, residual dipolar coupling values, transverse relaxation rates, and peaks present in the positive amide NOE spectra following assignment of most backbone resonances for the full-length SutA. Secondary shifts were calculated using TALOS as part of the PINE automated assignment server. RDCs were measured by manual comparison of inphase-anti-phase spectra between stretched gel and aqueous solution conditions. R_2_ values were calculated by fitting single exponential decay curves to peak integrals from spectra with increasing T_2_ delays. Positive NOE signal indicates that a peak was detected in the positive (^1^H-^15^N) NOE. The box indicates the location of the α-helix. **D.** One of many possible SutA structures modelled using the Robetta fragment server to incorporate chemical shift and RDC data, and PyRosetta. On the left, residues are colored by per-residue conservation score following alignment of 25 representative homologs (see Extended Materials and Methods for details). On the right, residues are colored by charge.

For structural characterization by NMR, we purified ^15^N- and ^13^C-labeled full-length SutA, as well as a ^15^N- and ^13^C-labeled construct that lacked most of the predicted disordered residues: SutA 46-101. We also constructed two deletion mutants (Figure 1B): SutA ∆N, retaining residues 41-105, and SutA ∆C, retaining residues 1-87.

We were able to assign resonances and determine backbone chemical shifts for about 85% of the residues of the full-length protein (Table 1). Low sequence complexity and large regions of disorder caused a high degree of overlap in the spectra and made assignment difficult; spectra from the 46-101 variant were easier to assign, and served as a starting point for assignment of the full-length SutA. We focused on measuring secondary-structure chemical shift index values, R_2_ relaxation rates, and ^1^H-^15^N NOE magnitude and sign to determine secondary-structure elements and degree of disorder for each residue that we could assign. We also embedded the protein in a stretched polyacrylamide gel to achieve weak alignment, and calculated residual dipolar couplings (RDCs) by measuring differences in in-phase–antiphase spectra between the isotropic solution sample and the anisotropic stretched-gel sample (Figure 1C). The results of these analyses lend credence to the bioinformatics predictions. Residues 56-76 show the positive Cα and CO and negative Hα secondary chemical shifts associated with an α-helix structure (18), and also show fast R_2_ relaxation rates and positive (^1^H-^15^N)NOE, suggesting that they are not disordered (19). RDCs for the helix region are also positive, as has been observed for α-helical regions of a partially denatured protein (20). The short β-strand is less strongly supported, but secondary shifts for those residues are mostly of the appropriate sign for a β-strand (albeit of small magnitudes). In the N-tail, a small number of residues have a positive NOE signal or secondary shifts that are not near zero, but in general, the residues of this region have the low R_2_, secondary shift, and RDC values that are characteristic of disorder. The C-tail has several residues that show somewhat higher R_2_ values and non-zero RDCs, suggestive of some degree of structure, but classic secondary structure elements are not apparent. We also compared ^15^N HSQC spectra for ^15^N-labeled ∆N and ∆C mutants to the full-length SutA (Figures S2 and S3). Deletion of either tail had minimal impact, affecting only the 2-4 residues adjacent to the newly created terminus.

The difficulty of making unambiguous assignments for all residues and the high likelihood that much of the protein is intrinsically disordered precluded building a full NMR-based structural model of SutA. To model some of the conformations that might be adopted by SutA, we used the Robetta Server and PyRosetta to perform low-resolution Monte Carlo–based modeling, using the chemical shifts and RDC values from our NMR analysis to guide fragment library construction (21–23). Figure 1D shows a resulting model. The most highly conserved residues are found in the α-helix, and the C-tail is also highly conserved among homologs that have it. The N-tail is less conserved and varies in length, but is generally quite acidic. Figure S4 shows additional models (see Extended Materials and Methods for modeling details).

### SutA binds to the β1 domain of RNAP

To investigate the binding interaction between SutA and RNAP, we used cross-linking and protein footprinting to map the region of RNAP with which SutA interacts. First we used the homobifunctional reagent bis(sulfosuccinimidyl)suberate (BS^3^), which cross-links primary amines that are within about 25 Å of each other (24). We added BS^3^ to complexes formed with purified core RNAP and SutA (Figure S2A), used the peptidase Glu-C to digest cross-linked complexes, and subjected the resulting fragments to LC-MS/MS. Analysis with the software package Protein Prospector (25) identified species that comprised one peptide from SutA and one peptide from RNAP (see Extended Materials and Methods and Figures S5 and S7), which allowed mapping of cross-link sites. We also used the photoreactive non-canonical amino acid p-benzoyl-L-phenylalanine (BPA), which, when irradiated with UV light, can form covalent bonds with a variety of moieties within 10 Å (26, 27). We introduced BPA at 9 different positions of SutA (residue 6, 11, 22, 54, 61, 74, 84, 89, or 100); we then formed complexes with purified core RNAP and each of the BPA-modified SutA proteins, irradiated them with UV light, and visualized cross-linked species following SDS-PAGE (Figure S6). For the most efficient cross-linkers (BPA at positions 54 and 84), we determined the sites of the cross-links on RNAP by identifying cross-linked peptides via StavroX (28) analysis of LC-MS/MS data after tryptic digest of the complexes (Figure S8).

Both cross-linking approaches identified interactions between the central region of SutA and the β1 domain or nearby regions of the β subunit of RNAP (Figure 2A and B, green and orange). All SutA residues participating in the cross-links were within the α-helix region (BS^3^) or just outside it (BPA). BPA cross-linking is sensitive to the orientations of the residues, so BPA residues within the helix that did not cross-link efficiently may not have been oriented optimally.

**Figure 2.**
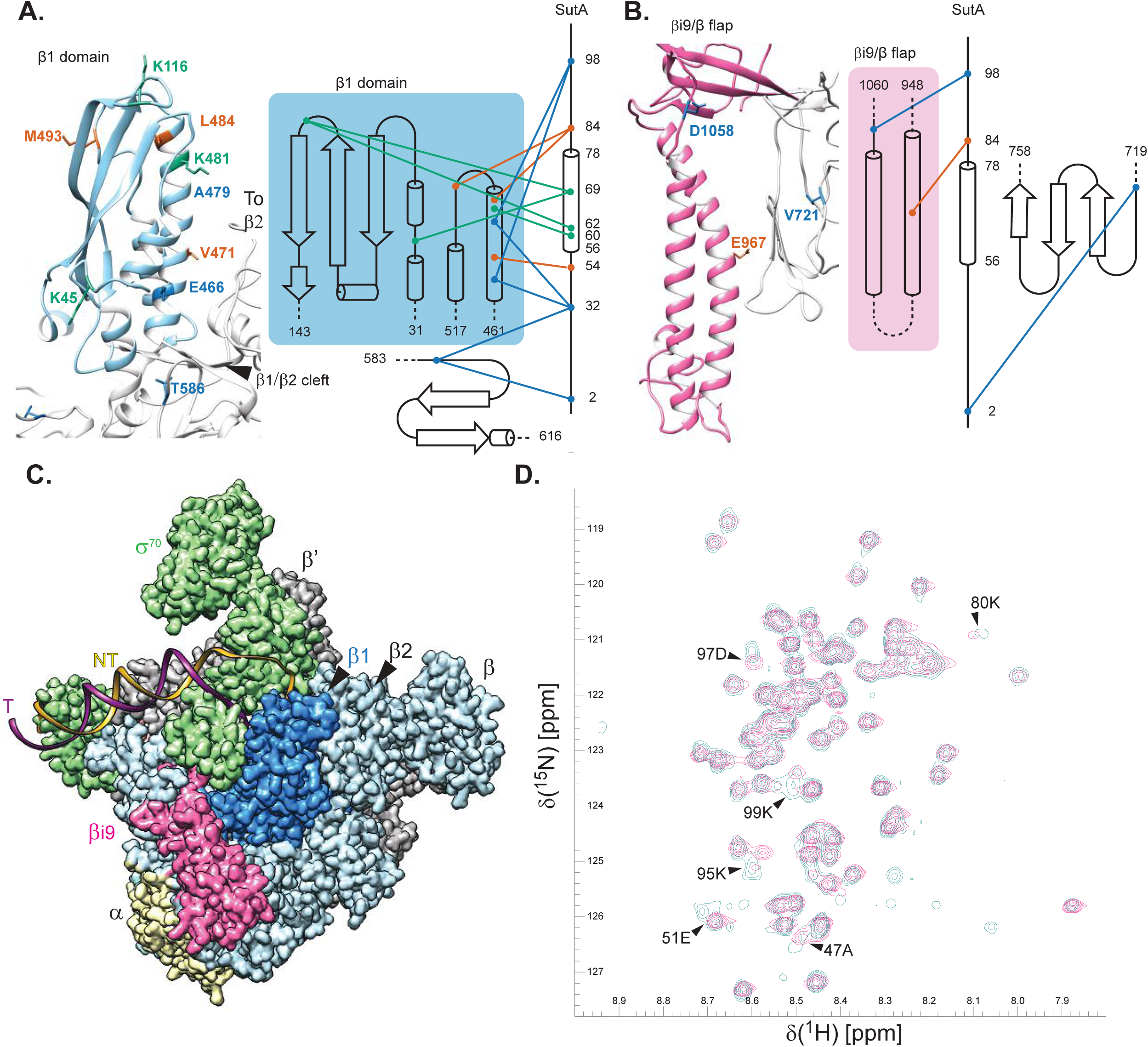
SutA interacts with the β subunit of RNAP. **A.** Contacts with the β1 domain. *P. aeruginosa* sequence was threaded onto a structural model of the *E. coli* β (PDB ID: 5UAG) for interpretation of cross-linking results. A topology diagram of the contacts inferred by cross-linking (BS^3^, green lines; BPA, orange lines) and FeBABE-mediated cleavage (blue lines). The contact residues on β1 are colored accordingly. Cross-linked residues were identified by LC-MS/MS, cleavage sites were determined by SDS-PAGE and Western blotting of the cleaved complexes, using a large-format gel system and Abcam antibody EPR18704, against the extreme C-terminus of the *E. coli* β. See text, SI, and Extended Materials and Methods for further details. **B.** Contacts around the βi9 and β flap domains. **C.** Cryo-EM structure of *E. coli* Eσ^70^ (PDB: 6CA0), indicating relative positions of β1 (darker blue, same as region shown in (A), and fragment purified for (D)) and βi9/β flap regions (pink, same as region shown in (B)). **D.** ^1^H-^15^N HSQC spectra showing that chemical shifts for a handful of residues are perturbed when ^15^N-labeled SutA is mixed with unlabeled β1 domain (pink) vs unlabeled σ^S^ (turquoise). A small number of extra peaks show up only in the σ^S^ mixture (turquoise, lower right quadrant); these are most likely due to very low levels of protein cleavage in the C-terminal disordered tail of SutA caused by a minor protease contaminant present in the σ^S^ protein preparation.

To identify the positions of the N- and C-tails, we designed variants of SutA for affinity cleavage experiments. We introduced cysteine residues at SutA position 2, 32, or 98 and conjugated the chelated iron reagent, iron-(S)-1-[*p*-(bromoacetamido)benzyl]EDTA (FeBABE), to these cysteines. FeBABE catalyzes localized (estimated to occur within 12 Å of the FeBABE moiety) hydroxyl radical cleavage (29). We assembled complexes with the FeBABE-modified SutA variants and core RNAP, initiated the cleavage reactions, and analyzed the cleavage products by SDS-PAGE followed by Western blotting with a monoclonal antibody against a peptide near the C-terminus of β. To map the cleavage sites, products were compared to C-terminal β fragments with known N-terminal endpoints (Figure S9). While the primary cleavage product of the N-terminal FeBABE (at residue 2; N-Fe) was in the cleft between the β1 and β2 (a.k.a. β lobe) domains, the strongest cleavage product of the C-terminal FeBABE (at residue 98; C-Fe) was in the long α-helix on the inside surface of β1 (designated α6 (30)), amongst BS^3^ and BPA cross-linking sites (Figure 2A, blue). The FeBABE at residue 32 induced cleavage at both β positions, suggesting that the N-tail remains mobile to some degree even when bound to RNAP.

We also detected possible interactions with βi9, which is an insertion in the β flap domain (31): BPA at residue 84 crosslinked to β967, and weak cleavage products were detected at β721 for the N-Fe variant and β1058 for the C-Fe variant (Figure 2B). β967 and β484/493 residues that were strongly cross-linked to BPA84 are too far apart to be reached from a single, stably bound position of SutA 84. However, we did not detect more than one shifted band after cross-linking with the 54 or 84 BPA variants (Figure S2B), suggesting that two separate sites on β are not likely to be occupied by two SutA molecules at the same time. Instead, it may be that SutA’s inherent flexibility, combined with a binding interaction with a surface on the outside of the β1 domain that allows some rotation or translation of SutA, could allow for all of the observed cross-links and cleavages.

To corroborate SutA-β interaction without cross-linking or cleavage and assess which residues of SutA might directly participate, we conducted an NMR experiment. We were able to purify only a small amount of soluble β1 domain (colored darker blue in Figure 2C), which we mixed with an equimolar amount of ^15^N-labeled full-length SutA. As a control to rule out non-specific interactions, we mixed SutA with an equimolar amount of σ^S^, which does not appear to bind SutA. Several SutA residues showed chemical shift perturbations in the β1 mixture, compared with the σ^S^ mixture (Figure 2D). Three of these residues, K95, D97 and K99, would be on the same side of an extended peptide chain in the C-tail, suggesting that this tail could directly interact with β1 in an extended conformation. However, the C-Fe SutA variant induced weaker cleavage than the N-Fe variant, suggesting that this interaction is probably not the only binding determinant. The other perturbed residues flank the α-helix, suggesting that the regions at the junctions with the flexible tails may change conformation upon binding to β.

### SutA activates the *rrn* promoter *in vitro*

Next we investigated SutA effects on transcription. We focused on the rRNA promoter because our ChIP data suggested that SutA directly affects *rrn* initiation (1). We first asked whether SutA affects transcription by the closely related *E. coli* RNAP, for which extensive *in vitro* tools are available. Overexpressing SutA in *E. coli* did not lead to *rrn* upregulation *in vivo* as it did in *P. aeruginosa* (Figure S10), necessitating the use of a cognate *P. aeruginosa in vitro* transcription system. We purified the core RNAP natively from a ∆*sutA* strain using a protocol originally designed for *E. coli* RNAP and previously used to purify RNAP from *P. aeruginosa* (32–34). The *P. aeruginosa* homologs of σ^S^, σ^70^, and DksA (as well as SutA) were heterologously expressed in *E. coli* with cleavable N-terminal 6xHis tags and purified by metal affinity and size-exclusion chromatography.

Unlike its well-studied *E. coli* counterpart, the *P. aeruginosa rrn* initiation region has not been characterized. We mapped the dominant *rrn* transcription start site using 5’-RACE to a cytidine 8 bp downstream of a -10 consensus sequence (Figure S11). We produced linear templates of 120-170 bp containing *the rrn* promoters and 42 bp of transcribed sequence for use in single-turnover initiation experiments (see supplement for details).

Transcription initiation proceeds via a multi-step pathway consisting of: 1)formation of a closed complex between the double-stranded DNA and the RNAP holoenzyme; 2)initial DNA strand separation, followed by isomerization through several open intermediates into a final open complex (OC) in which the +1 position of the template DNA strand is loaded into the active site and the downstream DNA duplex is stably held by RNAP; 3)initial abortive rounds of nucleotide addition; and 4)promoter clearance and transition into the elongation phase. Any of these steps could theoretically be affected by a regulator, and the details of this pathway differ at different promoters (35). Because much of the control of the *E. coli* P1 depends on the inherent instability of its OC (7), we sought to determine 1) whether the *P. aeruginosa rrn* has similar structure and response to regulatory inputs, and 2) whether SutA affects the *rrn* OC stability. The *P. aeruginosa rrn* promoter shares some of the features known to contribute to *E. coli rrn* P1 OC instability (Figure 3A): suboptimal spacing (16 nt vs the optimal 17-18 nt) between near-consensus -35 and -10 hexamers, a GC-rich discriminator region, and a C residue 2 nt downstream of the -10 hexamer that cannot make productive contacts to σ^70^ (36). However, the *P. aeruginosa* discriminator is 7 nt, which is longer than the optimal 6 nt, but still one base shorter than that of *E. coli*. Also, the initiating nucleotide is a cytidine rather than the adenosine or guanine iNTPs found in *E. coli,* indicating potentially different regulatory connections to cellular energy levels.

**Figure 3.**
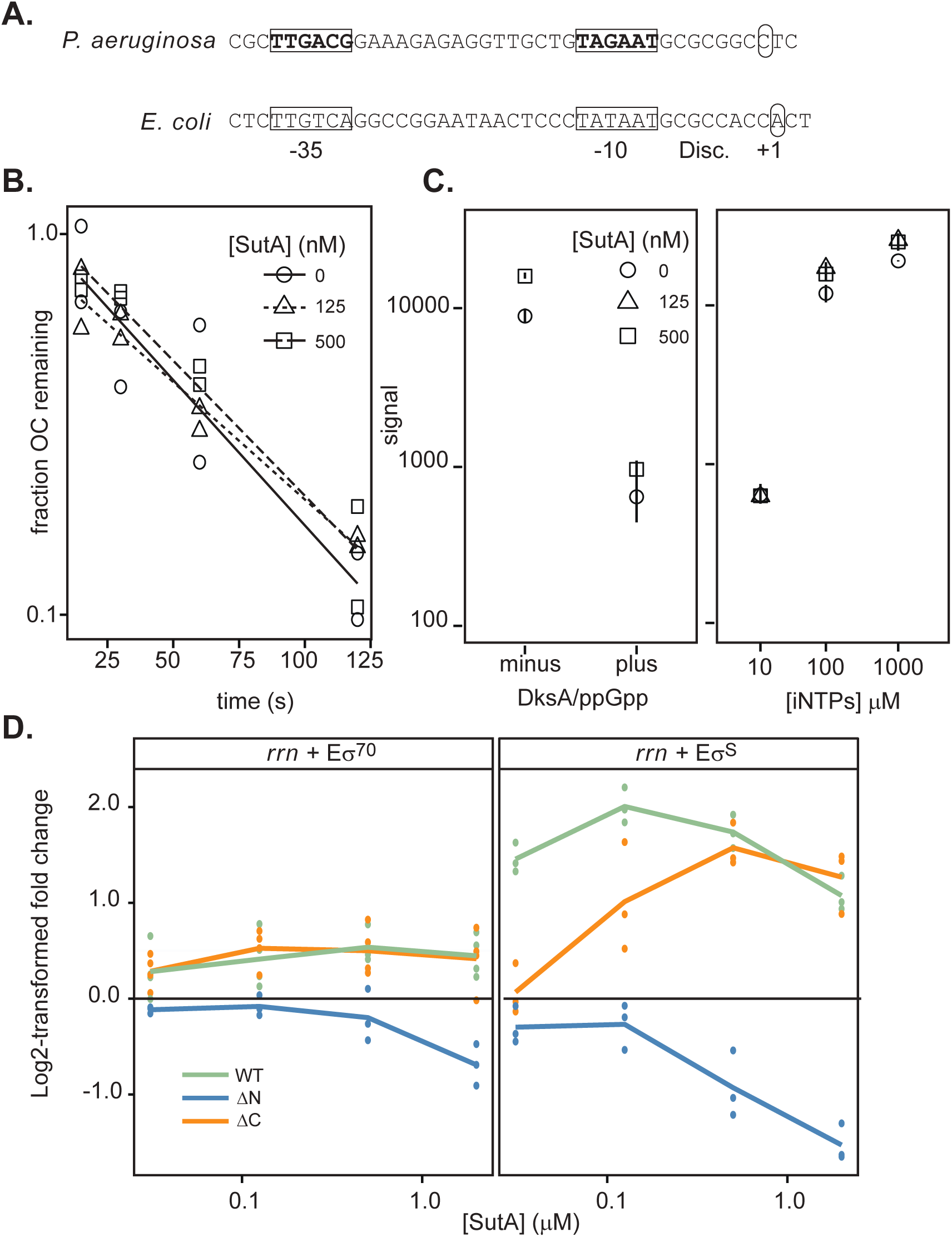
Effects of SutA variants on transcription initiation. **A.** *rrn* promoter sequences of *P. aeruginosa* and *E. coli* (P1). -10 and -35 motifs are indicated in bold and boxed, transcription start sites are indicated by circles, and the discriminator region is noted (Disc.). **B.** The heparin-resistant *P. aeruginosa rrn* OC is short-lived and its lifetime is not affected by SutA. The OC was formed with 20 nM Eσ^70^ (black) or Eσ^S^ (red) and 15 nM promoter DNA and challenged with heparin. NTPs were added at the indicated times and transcription was allowed to proceed for 8 minutes before quenching the reaction and running on a 20% gel. Reactions were performed at least in duplicate. **C.** DksA and ppGpp, or low [iNTPs] repress initiation from the *rrn* promoter, and SutA does not overcome these effects. Single turnover initiation reactions were performed using 20 nM Eσ^70^, 15 nM promoter DNA, 50 µM NTPs for the bases not labeled in the experiment, 20 µM cold NTP for the base carrying the ^32^P label, and 0.75 µCi α^32^P labeled GTP or CTP per 5 µl reaction, and 20 µg/ml heparin, in TGA buffer. In the left panel, 75 µM initiating dinucleotide was used and 500 nM SutA and/or 250 nM DksA and 2.5 µM ppGpp were added as indicated. On the right, varying concentrations of SutA and different concentrations of CTP and UTP, the first two nucleotides of the *rrn* transcript were used as indicated. RNAs were run on a 20% denaturing polyacrylamide gel and visualized by phosphorimaging. Symbols indicate the average value for the three replicates and lines represent the range of values observed in replicate experiments (n=3) (normalized such that the average signal for the 0 nM SutA condition for a given [iNTP] was the same across different gels). **D.** Amount of transcript produced in the presence of varying concentrations of SutA or SutA variant protein, compared to the amount produced in the absence of SutA, expressed as a log2-transformed ratio. Reactions were as described above. Individual replicate values are plotted (n≥3), and lines connect the average of all replicates at each concentration.

First, we directly measured the half-life of the heparin-resistant Eσ^70^ OC in a transcription-based assay. In contrast to what has been seen in *E. coli* (37), we detected some OC at standard salt concentrations and on a linear template, but its half-life was quite short, at about 45 seconds. Addition of SutA at 125 or 500 nM had no significant effect (Figures 3B and S12). Next, we measured effects of ppGpp/DksA and increasing [iNTPs], which repress and activate transcription from the *E. coli rrn* P1, respectively, in the presence or absence of SutA (Figures 3C, S13, and S14). As observed in *E. coli*, *rrn* transcription was strongly repressed at low [iNTPs] and by DksA/ppGpp, but SutA did not significantly counter these effects. Taken together, these results suggest that while the *P. aeruginosa rrn* promoter forms an inherently unstable OC, which is sensitive to regulatory inputs that utilize its instability, SutA does not alter the OC stability.

To directly measure the effects of SutA on transcription initiation, we performed single turnover initiation assays using the wild type (WT) SutA and the ∆N- and ∆C-tail variants described in Figure 1. Because Eσ^S^ binds the *rrn* locus *in vivo* during stationary phase in *E. coli* (38), we wanted to investigate whether SutA effects on *rrn* transcription *in vivo* could be mediated through Eσ^S^ or Eσ^70^, or both. We found that WT SutA increased transcription by both holoenzymes *in vitro*, but the magnitude of the effect was much larger for Eσ^S^ (up to 400% increase) than for Eσ^70^ (up to 70% increase) (Figures 3D, S15). In both cases, the effect saturated at concentrations of SutA between 125 and 500 nM. The acidic N-tail is strictly required for activation, as the ∆N mutant inhibited transcription in a dose-dependent manner. The ∆C mutant was still able to enhance transcription, albeit with a small shift in the concentration dependence evident with Eσ^S^. This shift may reflect C-tail interactions with Eσ^S^: we observed that the chemical shifts of three residues in the C-tail were perturbed upon mixing with the β1 domain.

### A disordered acidic loop in σ^70^ modulates SutA binding

We wondered what difference between σ^70^ and σ^S^ could explain the difference in SutA’s impact on *rrn* initiation by Eσ^70^ compared to Eσ^S^. Domains 2, 3, and 4 are highly similar, and both σ^70^ and σ^S^ have unstructured acidic regions, referred to as 1.1, near their N-termini (39). However, σ^70^ contains a large (~245 amino acids) insertion, termed the “non-conserved region” or NCR, which is not present in σ^S^ (Figure 4A). Crystal and cryoEM structures show that most of the NCR is situated relatively far from the β1 binding site of SutA, contacting the β’ subunit on the opposite side of the main channel of RNAP, but an acidic stretch of ~40 residues within the NCR is too flexible to be resolved in these structures (herein AL for <u>A</u>cidic <u>L</u>oop) (40, 41).

**Figure 4.**
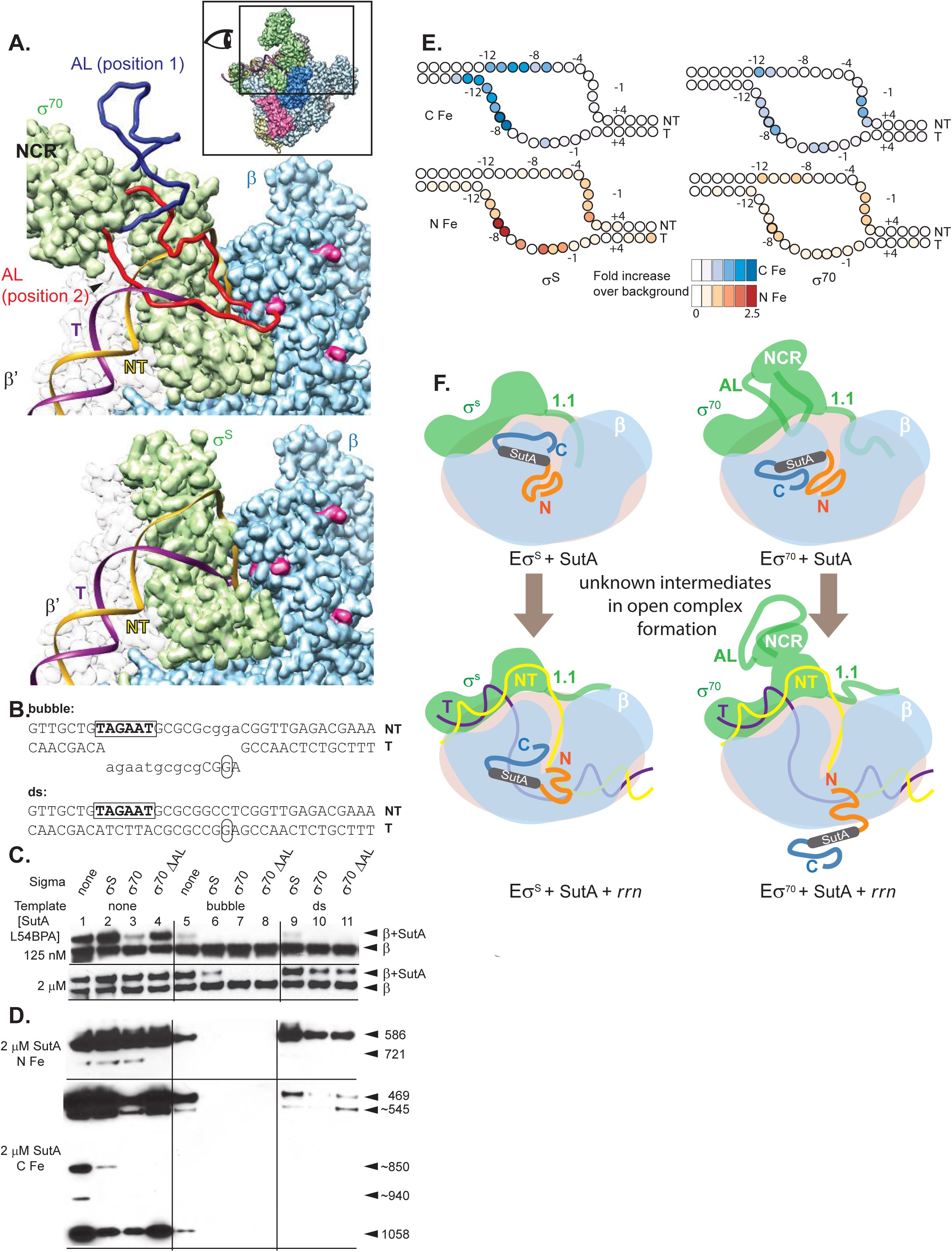
Both σ factor and DNA compete with SutA for binding to the core RNAP. **A.** Models based on *E. coli* σ^70^ and σ^S^ holoenzyme structures show potential interactions between σ factors and SutA. The inset shows the perspective and extent of this view relative to the holoenzyme structure shown in (2C). The *P. aeruginosa* β sequence was threaded onto an *E. coli* crystal structure (PBD: 5UAG), and then the β subunit from this was docked into the Eσ^70^ cryoEM structure (top) (PDB:6CA0) or the Eσ^S^ crystal structure (bottom) (PDB:5IPN). Residues showing cross-link or cleavage reactivity with SutA (Fig. 2) are colored magenta. Residues 168-212 of σ^70^, which are not visualized in the cryoEM structure, were modelled in as a flexible loop. Two different possible positions are shown (red and dark blue), one of which (red) could clearly clash with both the DNA and SutA positions (top). In contrast, σ^S^ does not appear likely to directly contact SutA (bottom). B. Sequence and structure of template DNA surrounding transcription start site. **C.** Western blot showing cross-linking of L54BPA SutA to β, in the context of different σ factors and promoter DNA. Reactions contained 100 nM RNAP, 100 nM DNA, 100 mM NaCl, and the indicated amounts of L54BPA SutA in TGA buffer. “∆AL” refers to a mutant of σ^70^ lacking amino acids 171-214 (*P. aeruginosa* sequence). **D.** Western blots showing β cleavage mediated by N-Fe or C-Fe SutA FeBABE conjugates. Reactions components besides SutA variant were the same as in (B). Sizes of cleavage products were estimated by comparison to β fragments of known sizes analyzed on large non-gradient gels (see supplements to figures 2 and 4); for some products (~), only approximate sizes can be determined. The blot for C-Fe was exposed for longer (4 minutes) than the blot for N-Fe (30 seconds). **E.** Cleavage of the DNA in the *rrn* OCs formed by Eσ^70^ or Eσ^S^ in the presence of N-Fe or C-Fe SutA, revealed by primer extension. Average log2-transformed enrichment in signal between the FeBABE reaction and a negative control reaction containing unmodified SutA, from triplicate measurements, is represented by color intensity for each base(C-Fe patterns are shown in blue and N-Fe patterns in orange). **F.** Model for SutA interaction with Eσ^S^ and Eσ^70^ holoenzymes, in the absence of DNA or with the *rrn* promoter. In the absence of DNA (top panels), patterns of crosslinking and FeBABE cleavage on RNAP are similar with Eσ^S^ to what is observed with core RNAP alone, suggesting that the N-tail is located in or near the cleft between β1 and β2, and the C-tail is located near the top of β1, where it may contribute to binding. With Eσ^70^, the AL that forms part of the NCR of σ^70^ appears to interfere with the C-tail of SutA, resulting in changes in the position and/or stability of the SutA interaction. When holoenzyme forms an OC with the *rrn* promoter (bottom panels), interaction with SutA is inhibited, suggesting that SutA may be displaced from its main interactions with the β1/β2 cleft. Low levels of crosslinking and cleavage are still detectable, especially in the Eσ^S^ context, but the loss of interaction suggests that SutA may exert its activating influence primarily on early intermediates in initiation, which are not structurally well-characterized.

To investigate possible interactions between the AL of σ^70^ and SutA, we threaded the *P. aeruginosa* sequence onto the β subunit of an *E. coli* RNAP crystal structure (42), docked that model into the recent cryoEM structure of the *E. coli* Eσ^70^ OC (41), and modeled the missing AL (using the *E. coli* sequence for both the structured and flexible regions of σ^70^) using the MODELLER software suite (43). The highly flexible AL could occupy a wide range of positions (e.g., Figure 4A, top), some of which would stay well above the DNA in the main channel (position 1) and some of which would clash with the DNA (position 2), and it could reach the β1 residues that participate in SutA cross-links, especially in the absence of DNA. In contrast, σ^S^ has no corresponding flexible region and remains far from the SutA cross-links (Figure 4A, bottom).

To determine whether the AL might contribute to the observed differences between Eσ^70^ and Eσ^S^ activation by SutA, we constructed and purified a *P. aeruginosa* σ^70^ mutant lacking residues 171-214 (∆AL), which correspond to the region missing in the *E. coli* structure, and repeated our cross-linking and cleavage assays using Eσ^70^, Eσ^S^, or Eσ^70^∆AL holoenzymes instead of just the core enzyme. In the absence of DNA, SutA L54BPA cross-linked to Eσ^70^ less efficiently than to E or Eσ^S^. Interestingly, Eσ^70^∆AL largely restored the cross-linking to the levels seen with E or Eσ^S^ (Figure 4C, lanes 1-4), suggesting that the σ^70^ AL modulates the SutA interaction with Eσ^70^. The difference in cross-linking efficiency between Eσ^70^ and Eσ^70^∆AL decreased at higher SutA concentrations, as might be expected if SutA and AL are competing to occupy a similar space.

### SutA competes with DNA in the final open complex

Our cross-linking and cleavage results suggested that SutA’s position on RNAP might allow it to compete with the promoter DNA. To explore this possibility, we added to our crosslinking assay either a double-stranded (ds) *rrn* promoter DNA or a bubble template in which the region of the DNA that forms the transcription bubble in the OC was non-complementary (Figure 4B). The dsDNA requires σ to melt the DNA strands, and will support a native population of promoter complex intermediates. By contrast, the bubble template obviates the need for σ and would be expected to stabilize an OC formed with the holoenzyme, but this complex may not represent the dominant native complex, as the *E. coli rrn* P1 does not form a stable final OC (37). The addition of the bubble DNA had a large negative effect on SutA binding that was synergistic with the presence of σ (Figure 4C, lanes 5-8). Cross-linking could still be readily detected in the absence of σ, and to a lesser extent when σ^S^ was present, but not with either σ^70^ or σ^70^∆AL; longer exposures revealed that cross-linking did occur at low efficiency (Figure S16). Addition of dsDNA allowed more SutA binding, but still less than in the absence of DNA (Figure 4C, lanes 9-11).

### The flexible SutA tails approach the transcription bubble

The flexible, acidic σ1.1 region competes with promoter DNA for binding to the main channel of RNAP, but still enhances initiation from some promoters (44). We wondered whether SutA might likewise use its acidic N-tail to compete with promoter DNA, but enhance initiation at the *rrn* promoter. The BPA crosslinking reports on the interactions established by the central region of SutA, but gives no information on the position of its flexible tails, and a decrease in crosslinking could be due to a loss of binding or a change of SutA conformation. We used FeBABE cleavage assays with the N-Fe and C-Fe SutA variants (Figures 4D, S17) to address these questions. We found that the addition of σ^70^ had a much larger negative effect on cleavage induced by C-Fe than on cleavage induced by N-Fe. The σ^70^∆AL mutant partially restored C-Fe cleavage levels to those observed with the core enzyme or Eσ^S^, but caused a decrease compared to σ^70^ in the N-Fe cleavage at residue 721. This suggests that the σ^70^AL does not fully displace SutA, but instead interferes with a binding interaction of the C-tail (Figure 4F; compare top right to top left panel), decreasing the crosslinking efficiency of the 54BPA variant, consistent with our observation that the ∆C mutant required higher concentrations for maximal activity on Eσ^S^ but not on Eσ^70^ (Figure 3D).

By contrast, the effects of template DNA were similar for both C-Fe and N-Fe cleavage reactions, as well as for BPA crosslinking. This suggested that DNA might induce SutA dissociation, rather than its subtle repositioning (Figure 4F; compare top to bottom panels), prompting us to investigate whether SutA and DNA could form a ternary complex with RNAP holoenzyme. We measured FeBABE SutA-dependent cleavage of the template and non-template DNA strands using primer extension. We saw stronger cleavage with Eσ^S^ than with Eσ^70^, but in both cases the signal was relatively weak, as might be expected for a factor that does not directly bind DNA (Figures 4E, S19). In the Eσ^S^ complex, C-Fe induced cleavage of both strands between residues -8 and -12, suggesting that it remains near the upstream fork junction of the transcription bubble. N-Fe cleaved the template strand near the upstream junction but also cleaved both strands further downstream. For Eσ^70^, the cleavage was weaker overall and showed a different pattern; for C-Fe in particular, more cleavage took place on the downstream region of the non-template strand. This difference could reflect the AL-mediated repositioning of the C-tail.

Our results argue that SutA may not stably bind the final OC. However, SutA-induced DNA cleavage demonstrates that SutA does bind to some promoter complexes, in turn suggesting that an earlier intermediate in the initiation pathway may be the main target of SutA activity (Figure 4F).

## DISCUSSION

As part of their response to fluctuating environmental conditions, bacterial cells produce regulators that directly bind RNAP and modify its behavior, eliciting global changes in gene expression patterns, in addition to producing different DNA-binding transcription factors that help recruit RNAP to specific genes (reviewed in (45)). We previously identified SutA as a global regulator that binds RNAP and contributes to a broad response to protracted growth arrest, enhancing ongoing, low-level expression of housekeeping genes (1). SutA directly affects initiation at the *rrn* promoter, prompting comparison to other regulators that affect rRNA expression by directly binding RNAP, such as DksA, ppGpp, and CarD. DksA and ppGpp, which appear to operate similarly in *P. aeruginosa* and *E. coli*, broadly destabilize OCs, leading to repression of *rrn* P1 and activation of amino acid biosynthesis genes in response to nutrient downshifts (46). In contrast, CarD, constitutively expressed in many non-Gammaproteobacteria, broadly stabilizes OCs and modestly enhances rRNA transcription (approximately 3-fold *in vitro*) (12). SutA is distinct from both of these examples: though it acts on a *P. aeruginosa rrn* promoter that also forms an unstable OC with Eσ^70^, its activity does not affect the stability of this complex. This difference is perhaps unsurprising, as both its structure and interactions with RNAP are also distinct, as discussed below.

### A model for SutA interactions with promoter complexes

Unlike DksA and CarD, which have well-defined structures (12, 47), SutA is largely intrinsically disordered, with its flexible tails playing key functional roles. SutA binds to the β1 domain of RNAP; its crosslinking and cleavage interactions suggest a binding site that is close to but distinct from that of CarD (12), and far from the sites occupied by DksA and ppGpp (11, 48). Although the extreme flexibility of SutA and the relatively large distances over which our cross-linking and cleavage reagents could act (10-25 Å) preclude precise docking of SutA onto RNAP, a binding site on the outside of the β1 domain is consistent with our data. SutA failed to activate *rrn* transcription in *E. coli in vivo* (Figure S10) and failed to bind the *E. coli* Eσ^70^ *in vitro* (Figures S16 and S17), suggesting that its binding site is in a region that is different between the two polymerases. Most of the β residues are identical (72%) or similar (87%) between *E. coli* and *P. aeruginosa*, but two β1 loops that contain residues involved in BS^3^ cross-linking (K45 and K116) are among a small number of amino acid sequences with reduced similarity (Figure S20). From such a binding site for the SutA helix, its flexible tails could reach into the main channel of RNAP through the cleft between β1 and β2. The fact that FeBABE-modified SutA variants could catalyze cleavage of the *rrn* DNA suggests that the tails approach the DNA in some promoter complexes.

The β1/β2 cleft has been shown to accommodate the non-template strand during the early rounds of nucleotide addition, when it must scrunch to allow additional bases of the downstream DNA to enter the enzyme before the upstream contacts are released (49); on *E. coli rrnB* P1, scrunching occurs even before initiation (50). Interestingly, the β1/β2 cleft is also the site of several point mutations that suppress the auxotrophy phenotype of a *∆dksA* mutant in *E. coli*, consistent with a model where modulating its interaction with the DNA could be functionally important in growth-phase-dependent gene expression regulation (51). Our results show that the fully melted promoter DNA inhibits SutA binding, suggesting that DNA and SutA may compete for similar contacts with RNAP on β1 or near the β1/β2 cleft. SutA (and especially its N-tail) could positively influence the formation of the *rrn* OC through interaction with early promoter complex intermediates, and then be displaced as the final OC forms, potentially leading to its dissociation (Figure 4F). This is analogous to the regulatory mechanism of σ^70^ 1.1, an acidic flexible region that binds in the main RNAP channel in early promoter complexes and must be ejected to accommodate the promoter DNA that binds to the same site in the final OC (44, 52–55). Like SutA, σ^70^ 1.1 stimulates initiation at some promoters but does not affect the stability of their OCs (44).

### The roles of acidic disordered regions in SutA regulation and beyond

In addition to the critical role of SutA’s unstructured acidic N-tail in mediating its enhancement of *rrn* transcription, we also found that an unstructured, acidic region of σ^70^, the AL, directly modulates SutA’s activity, possibly by interfering with the ability of the C-tail to bind β1. The AL is part of the NCR region that is unique to σ^70^ (56), so this interaction may contribute to the difference in the SutA’s effects on initiation by Eσ^70^ versus Eσ^S^. The NCR makes contacts with the upstream DNA duplex (41), and our modeling suggests that the AL could be positioned near the upstream junction of the transcription bubble. Like region 1.1 and the SutA N-tail, the σ^70^ AL is a highly dynamic element that could modulate the DNA trajectory in early intermediates in open complex formation, before the bubble is locked in place in the final OC, and we found that in addition to affecting the RNAP-SutA interaction, σ^70^ ∆AL also has mild defects in initiation at the *rrn* promoter on its own (Figure S18). Since the OC formation pathways and the relative occupancies of the intermediates vary among promoters (37), the effects of these unstructured elements are expected to be distinct for different promoters.

Dynamic interactions of intrinsically disordered (and often highly acidic) modules play key roles in eukaryotic transcriptional regulators, bacteriophage proteins, and σ factors. Unstructured regions can gain access to and remodel dynamic regions of transcription complexes, as in the case of the phage proteins Gp2 and Nun, leading to inhibition of RNA synthesis (57, 58). They also can bind or mimic flexible nucleic acid sequences, as in the case of the λN protein (59) or σ1.1 (55), and activate transcription. In the case of eukaryotic transcriptional activators such as Gcn4 or Ino2, they can serve as flexible protein-protein interaction domains, capable of mediating interactions whose structural constraints vary depending on nuances of nucleic acid sequences, chromatin states, and other aspects of the intracellular environment (60–62). While their disordered nature has made many of these domains difficult to study using traditional structural and biochemical approaches, it is becoming increasingly clear that they play critical roles in many aspects of transcriptional regulation in all domains of life, and SutA adds to this growing body of evidence.

### Implications for the *in vivo* role of SutA during slow growth

Our finding that SutA can differentially impact transcription in a σ-dependent manner could have important implications *in vivo* during growth arrest. σ^70^ and σ^S^ are closely related σ factors with partially overlapping promoter specificities (40, 56, 63). Our previous RNA-Seq results imply that SutA interactions with both holoenzymes are likely to be functionally relevant, as some affected genes are *bona fide* σ^S^ regulon members (63), but overall the affected genes are biased toward classic targets of σ^70^ (1). In *E. coli*, the activities and relative abundances of Eσ^S^ and Eσ^70^ change throughout different growth phases, with σ^S^ upregulated at the transition to stationary phase (64, 65), and much of Eσ^70^ sequestered by the 6S RNA late in stationary phase (66). In addition, Eσ^S^ and Eσ^70^ appear to be differentially sensitive to changes in cellular conditions that occur during stationary phase, such as an overall decrease in the negative supercoiling of the chromosome, shifts in patterns of nucleoid associated proteins, and different concentrations of solutes. Moreover, in specific cases that have been examined *in vitro*, Eσ^S^ initiation efficiency increases under the stationary phase-associated condition, while Eσ^70^ initiation efficiency decreases (67, 68). These characteristics of Eσ^S^ may in part explain why σ^S^ ChIP signal at the *rrn* promoters increases in stationary phase in *E. coli* and σ^70^ ChIP signal decreases (38). The ability of SutA to enhance initiation by Eσ^S^ and Eσ^70^ differentially could allow greater flexibility during different stages of growth arrest. For example, SutA may enable baseline levels of housekeeping gene expression regardless of which holoenzyme ismost available and active, or could allow for combinatorial control whereby promoter activity during dormancy would be synergistically affected by σ preference and SutA interaction. More work will be required to fully understand how SutA contributes to the regulatory architecture that allows *P. aeruginosa* to thrive during dormancy, but this study represents important mechanistic insight into the function of this global regulator.

## Supporting information

## ACKNOWLEDGMENTS

We thank Ben Ramirez (University of Illinois at Chicago) for helping us with preliminary NMR studies of SutA, Jacqueline Barton (Caltech) for giving us access to her lab to perform experiments involving radioactivity, Nate Glasser for help with HPLC measurements to quantify SutA, Hsiau-Wei (Jack) Lee and Aimee Marceau (University of California, Santa Cruz) for help with the NMR binding experiment using the Bruker AVIII HD 800 MHz NMR, Weidong Hu (City of Hope) for help with NMR experiments using the Bruker AV III 700 MHz spectrometer, and Julia Kardon and Niels Bradshaw (Brandeis University) and members of the Newman lab for feedback on the project at different stages. MB was supported by a post-doctoral fellowship from the Cystic Fibrosis Foundation. Grants from the NIH (GM067153) to IA and grants from the HHMI and NIH (5R01HL117328-03 and 1R01AI127850-01A1) to DKN supported this work. The Proteome Exploration Laboratory is supported by the Beckman Institute and NIH 1S10OD02001301. This work was also supported by the Institute for Collaborative Biotechnologies through grant W911NF-09-0001 from the U.S. Army Research Office. The content of the information does not necessarily reflect the position or the policy of the Government, and no official endorsement should be inferred.

## MATERIALS AND METHODS

See Extended Materials and Methods in SI for additional details about all experiments, for strain construction details, and for tables of strains and primers used.

### Protein purifications

*P. aeruginosa* core RNAP was purified as previously described(32–34). N-terminal 6xHis-tagged SutA, SutA variants, DksA, σ^S^, σ^70^, σ^70^∆AL, and β1 were heterologously expressed in *E. coli* and purified by standard metal affinitiy chromatography followed by cleavage of the 6xHis tag with TEV protease and size exclusion chromatography. For NMR experiments, cells were grown in minimal media prepared with ^15^NH_4_Cl or ^13^C glucose or both. For BPA crosslinking, amber stop codons were introduced at positions of interest and BPA was incorporated via amber suppression following co-transformation of the SutA plasmid with pEVOL-pBpF as previously described (26). For preparation of FeBABE variants, cysteine residues were introduced at positions of interest (SutA has no natural cysteines) and following purification of the protein, the FeBABE moeity was conjugated to the cysteine as previously described (29). Conjugation efficiencies were estimated to be 57%, 38%, and 76% respectively for the residue 2, 32, and 98 variants.

### NMR experiments

Data were collected from SutA proteins at concentrations of 300 µM in a buffer containing 20 mM sodium phosphate, pH 7.0, 100 mM sodium chloride, and 10% D_2_O. For the 46-101 variant, the following spectra were acquired on a Varian Inova 600 MHz NMR with a triple resonance inverse probe running VnmrJ 4.2A: ^15^N HSQC, ^13^C HSQC, HNCO, HNCA, HNCACB, CBCACONH, HNCOCA, HNCACO, CCONH, and ^15^N HSQC experiments modified for measurement of T_2_ and of ^15^N-^1^H NOE. For the full-length protein, ^15^N HSQC, ^13^C HSQC, HNCACB, and CBCACONH spectra were acquired at 7 °C on a Bruker AV III 700 MHz spectrometer with a TCI cryoprobe running Topspin 3.2, but ^15^N HSQC experiments modified for measurement of T_2_ and of ^15^N-^1^H NOE were collected on the Varian Inova 600 MHz NMR, as were ^15^N HSQC spectra for the SutA ∆N and SutA ∆C SutA proteins. ^15^N^13^C-labeled full-lengthSutA was embedded in a stretched polyacrylamide gel for measurement of residual dipolar couplings as previously described (20), using the Varian Inova 600 MHz NMR. To assess SutA binding to β1 by NMR, ^15^N-labeled SutA and β1 fragment were buffer exchanged into 20 mM sodium phosphate, pH 7.0, 100 mM sodium chloride and the resulting complex was isolated by size exclusion chromatography, resulting in a final concentration of complex of approximately 25 µM. In addition, ^15^N-labeled SutA was mixed with σ^S^ at 50 µM each. ^15^N HSQC spectra were acquired on a Bruker 800 MHZ AV III HD spectrometer with a TCI cryoprobe at 25 °C. Peak assignments and analysis were done using the PINE Server, CcpNmr Analysis Suite, and MestreNova software.

### Crosslinking experiments and data analysis

BS^3^ crosslinking of core RNAP and SutA was carried out as previously described (24) with modifications. Crosslinked complexes were subjected to in-solution digestion by the Glu-C peptidase, and the resulting fragments were analyzed by LC-MS/MS on an Orbitrap Elite Hybrid Ion Trap MS. Crosslinked pepties were identified as previously described, with modifications (25). BPA crosslinking was achieved by irradiating RNAP core or holoenzyme complexes with UV light from an Omnicure S2000 lamp, complexes were digested in solution with trypsin, and analyzed by LC-MS/MS on a Q Exactive HF Orbitrap MS. Crosslinked peptides were identified using the StavroX software package (28).

### FeBABE experiments and analysis

Cleavage reactions of holoenzyme complexes assembled in TGA buffer were initiated by the addition of ascorbate and hydrogen peroxide to final concentrations of 5 mM each, as previously described (29). For measuring protein cleavage, reactions were quenched by the addition of SDS loading buffer and were evaluated by SDS-PAGE followed by Western blotting, using a monoclonal antibody raised against a peptide from the extreme C-terminus of *E. coli* β (EPR18704 from Abcam). To generate standards for size comparison, several different C-terminal fragments of RpoB with endpoints ranging from aa 355 to aa 1062 were overexpressed in *E. coli* and crude lysates from these strains were subjected to SDS-PAGE and Western blotting alongside the FeBABE cleavage products. For measuring DNA cleavage, reactions were quenched with thiourea and treated with proteinase K. The DNA was precipitated, and subjected to primer extension using Cy3- or Cy5-labeled primers against the non-template or template strand respectively.. Products were separated on denaturing 12% polyacrylamide gels and imaged by laser scanner.

### In vitro transcription experiments

Experiments were carried out as previously described with minor modifications (46, 69).

## REFERENCES

1. Babin BM, et al. (2016) SutA is a bacterial transcription factor expressed during slow growth in Pseudomonas aeruginosa. Proceedings of the National Academy of Sciences of the United States of America 113(5):E597–605.

2. Bergkessel M, Basta DW, & Newman DK (2016) The physiology of growth arrest: uniting molecular and environmental microbiology. Nature Reviews. Microbiology 14(9):549–562.

3. Udikovic-Kolic N, Wichmann F, Broderick NA, & Handelsman J (2014) Bloom of resident antibiotic-resistant bacteria in soil following manure fertilization. Proceedings of the National Academy of Sciences of the United States of America 111(42):15202–15207.

4. Olivares J, et al. (2013) The intrinsic resistome of bacterial pathogens. Frontiers in Microbiology 4:103.

5. Babin BM, et al. (2017) Selective Proteomic Analysis of Antibiotic-Tolerant Cellular Subpopulations in Pseudomonas aeruginosa Biofilm. mBio 8(5):16.

6. Ciofu O, Tolker-Nielsen T, Jensen PO, Wang H, & Hoiby N (2015) Antimicrobial resistance, respiratory tract infections and role of biofilms in lung infections in cystic fibrosis patients. Adv Drug Deliv Rev 85:7–23.

7. Paul BJ, Ross W, Gaal T, & Gourse RL (2004) rRNA transcription in Escherichia coli. Annu Rev Genet 38:749–770.

8. Murray HD, Schneider DA, & Gourse RL (2003) Control of rRNA Expression by Small Molecules Is Dynamic and Nonredundant. Molecular Cell 12(1):125–134.

9. Murray HD, Appleman JA, & Gourse RL (2003) Regulation of the Escherichia coli rrnB P2 Promoter. Journal of Bacteriology 185(1):28–34.

10. Murray HD & Gourse RL (2004) Unique roles of the rrn P2 rRNA promoters in Escherichia coli. Molecular Microbiology 52(5):1375–1387.

11. Ross W, et al. (2016) ppGpp Binding to a Site at the RNAP-DksA Interface Accounts for Its Dramatic Effects on Transcription Initiation during the Stringent Response. Molecular Cell 62(6):811–823.

12. Bae B, et al. (2015) CarD uses a minor groove wedge mechanism to stabilize the RNA polymerase open promoter complex. eLife 4.

13. Leirmo S & Gourse RL (1991) Factor-independent activation of Escherichia coli rRNA transcription. I. Kinetic analysis of the roles of the upstream activator region and supercoiling on transcription of the rrnB P1 promoter in vitro. J Mol Biol 220(3):555–568.

14. Buchan DW, Minneci F, Nugent TC, Bryson K, & Jones DT (2013) Scalable web services for the PSIPRED Protein Analysis Workbench. Nucleic Acids Research 41(Web Server issue):W349–357.

15. Drozdetskiy A, Cole C, Procter J, & Barton GJ (2015) JPred4: a protein secondary structure prediction server. Nucleic Acids Research 43(W1):W389–394.

16. Jones DT & Cozzetto D (2015) DISOPRED3: precise disordered region predictions with annotated protein-binding activity. Bioinformatics 31(6):857–863.

17. Williams KP, et al. (2010) Phylogeny of Gammaproteobacteria. Journal of Bacteriology 192(9):2305–2314.

18. Wishart DS, Sykes BD, & Richards FM (1991) Relationship between nuclear magnetic resonance chemical shift and protein secondary structure. J Mol Biol 222(2):311–333.

19. Reddy T & Rainey JK (2010) Interpretation of biomolecular NMR spin relaxation parameters. Biochemistry and cell biology = Biochimie et biologie cellulaire 88(2):131–142.

20. Mohana-Borges R, Goto NK, Kroon GJ, Dyson HJ, & Wright PE (2004) Structural characterization of unfolded states of apomyoglobin using residual dipolar couplings. J Mol Biol 340(5):1131–1142.

21. Bowers PM, Strauss CE, & Baker D (2000) De novo protein structure determination using sparse NMR data. J Biomol NMR 18(4):311–318.

22. Rohl CA & Baker D (2002) De novo determination of protein backbone structure from residual dipolar couplings using Rosetta. J Am Chem Soc 124(11):2723–2729.

23. Kim DE, Chivian D, & Baker D (2004) Protein structure prediction and analysis using the Robetta server. Nucleic Acids Research 32(Web Server issue):W526–531.

24. Rappsilber J (2011) The beginning of a beautiful friendship: cross-linking/mass spectrometry and modelling of proteins and multi-protein complexes. Journal of Structural Biology 173(3):530–540.

25. Trnka MJ, Baker PR, Robinson PJ, Burlingame AL, & Chalkley RJ (2014) Matching cross-linked peptide spectra: only as good as the worse identification. Mol Cell Proteomics 13(2):420–434.

26. Chin JW, Martin AB, King DS, Wang L, & Schultz PG (2002) Addition of a photocrosslinking amino acid to the genetic code of Escherichia coli. Proceedings of the National Academy of Sciences of the United States of America 99(17):11020–11024.

27. Kauer JC, Erickson-Viitanen S, Wolfe HR, & DeGrado WF (1986) p-Benzoyl-L-phenylalanine, A New Photoreactive Amino Acid. The Journal of Biological Chemistry 261(23):6.

28. Götze M, et al. (2012) StavroX—A Software for Analyzing Crosslinked Products in Protein Interaction Studies. J. Am. Soc. Mass Spectrom. 23:12.

29. Meares CF, Datwyler SA, Schmidt BD, Owens J, & Ishihama A (2003) Principles and Methods of Affinity Cleavage in Studying Transcription. Methods in Enzymology 371:25.

30. Lane WJ & Darst SA (2010) Molecular evolution of multisubunit RNA polymerases: sequence analysis. J Mol Biol 395(4):671–685.

31. Opalka N, et al. (2010) Complete structural model of Escherichia coli RNA polymerase from a hybrid approach. PLoS Biology 8(9).

32. Kuznedelov K, et al. (2011) The antibacterial threaded-lasso peptide capistruin inhibits bacterial RNA polymerase. J Mol Biol 412(5):842–848.

33. Hager DA, Jin DJ, & Burgess RR (1990) Use of Mono Q high-resolution ion-exchange chromatography to obtain highly pure and active Escherichia coli RNA polymerase. Biochemistry 29(34):7890–7894.

34. Burgess RR & Jendrisak JJ (1975) A procedure for the rapid, large-scall purification of Escherichia coli DNA-dependent RNA polymerase involving Polymin P precipitation and DNA-cellulose chromatography. Biochemistry 14(21):4634–4638.

35. Ruff EF, Record MT, Jr., & Artsimovitch I (2015) Initial events in bacterial transcription initiation. Biomolecules 5(2):1035–1062.

36. Haugen SP, et al. (2006) rRNA promoter regulation by nonoptimal binding of sigma region 1.2: an additional recognition element for RNA polymerase. Cell 125(6):1069–1082.

37. Ruff EF, et al. (2015) E. coli RNA Polymerase Determinants of Open Complex Lifetime and Structure. J Mol Biol 427(15):2435–2450.

38. Raffaelle M, Kanin EI, Vogt J, Burgess RR, & Ansari AZ (2005) Holoenzyme switching and stochastic release of sigma factors from RNA polymerase in vivo. Molecular Cell 20(3):357–366.

39. Gowrishankar J, Yamamoto K, Subbarayan PR, & Ishihama A (2003) In vitro Properties of RpoS (σ^S^) Mutants of Escherichia coli with Postulated N-Terminal Subregion 1.1 or C-Terminal Region 4 Deleted. Journal of Bacteriology 185(8):2673–2679.

40. Liu B, Zuo Y, & Steitz TA (2016) Structures of E. coli sigmaS-transcription initiation complexes provide new insights into polymerase mechanism. Proceedings of the National Academy of Sciences of the United States of America 113(15):4051–4056.

41. Narayanan A, et al. (2018) Cryo-EM structure of Escherichia coli sigma(70) RNA polymerase and promoter DNA complex revealed a role of sigma non-conserved region during the open complex formation. The Journal of Biological Chemistry 293(19):7367–7375.

42. Molodtsov V, Scharf NT, Stefan MA, Garcia GA, & Murakami KS (2017) Structural basis for rifamycin resistance of bacterial RNA polymerase by the three most clinically important RpoB mutations found in Mycobacterium tuberculosis. Molecular Microbiology 103(6):1034–1045.

43. Yang Z, et al. (2012) UCSF Chimera, MODELLER, and IMP: an integrated modeling system. Journal of Structural Biology 179(3):269–278.

44. Vuthoori S, Bowers CW, McCracken A, Dombroski AJ, & Hinton DM (2001) Domain 1.1 of the sigma(70) subunit of Escherichia coli RNA polymerase modulates the formation of stable polymerase/promoter complexes. J Mol Biol 309(3):561–572.

45. Browning DF & Busby SJ (2016) Local and global regulation of transcription initiation in bacteria. Nature Reviews. Microbiology 14(10):638–650.

46. Paul BJ, et al. (2004) DksA: a critical component of the transcription initiation machinery that potentiates the regulation of rRNA promoters by ppGpp and the initiating NTP. Cell 118(3):311–322.

47. Perederina A, et al. (2004) Regulation through the secondary channel--structural framework for ppGpp-DksA synergism during transcription. Cell 118(3):297–309.

48. Ross W, Vrentas CE, Sanchez-Vazquez P, Gaal T, & Gourse RL (2013) The magic spot: a ppGpp binding site on E. coli RNA polymerase responsible for regulation of transcription initiation. Molecular Cell 50(3):420–429.

49. Winkelman JT, et al. (2015) Crosslink Mapping at Amino Acid-Base Resolution Reveals the Path of Scrunched DNA in Initial Transcribing Complexes. Molecular Cell 59(5):768–780.

50. Winkelman JT, Chandrangsu P, Ross W, & Gourse RL (2016) Open complex scrunching before nucleotide addition accounts for the unusual transcription start site of E. coli ribosomal RNA promoters. Proceedings of the National Academy of Sciences of the United States of America 113(13):E1787–1795.

51. Rutherford ST, Villers CL, Lee JH, Ross W, & Gourse RL (2009) Allosteric control of Escherichia coli rRNA promoter complexes by DksA. Genes & Development 23(2):236–248.

52. Wilson C & Dombroski AJ (1997) Region 1 of sigma70 is required for efficient isomerization and initiation of transcription by Escherichia coli RNA polymerase. J Mol Biol 267(1):60–74.

53. Mekler V, et al. (2002) Structural Organization of Bacterial RNA Polymerase Holoenzyme and the RNA Polymerase-Promoter Open Complex. Cell 108:16.

54. Hook-Barnard IG & Hinton DM (2009) The promoter spacer influences transcription initiation via sigma70 region 1.1 of Escherichia coli RNA polymerase. Proceedings of the National Academy of Sciences of the United States of America 106(3):737–742.

55. Murakami KS (2013) X-ray crystal structure of Escherichia coli RNA polymerase sigma70 holoenzyme. The Journal of Biological Chemistry 288(13):9126–9134.

56. Feklistov A, Sharon BD, Darst SA, & Gross CA (2014) Bacterial sigma factors: a historical, structural, and genomic perspective. Annu Rev Microbiol 68:357–376.

57. Bae B, et al. (2013) Phage T7 Gp2 inhibition of Escherichia coli RNA polymerase involves misappropriation of σ70 domain 1.1. Proceedings of the National Academy of Sciences of the United States of America 110(49):6.

58. Kang JY, et al. (2017) Structural basis of transcription arrest by coliphage HK022 Nun in an Escherichia coli RNA polymerase elongation complex. eLife 6.

59. Said N, et al. (2017) Structural basis for lambdaN-dependent processive transcription antitermination. Nature Microbiology 2:17062.

60. Staller MV, et al. (2018) A High-Throughput Mutational Scan of an Intrinsically Disordered Acidic Transcriptional Activation Domain. Cell Syst 6(4):444–455 e446.

61. Tuttle LM, et al. (2018) Gcn4-Mediator Specificity Is Mediated by a Large and Dynamic Fuzzy Protein-Protein Complex. Cell Reports 22(12):3251–3264.

62. Pacheco D, et al. (2018) Transcription Activation Domains of the Yeast Factors Met4 and Ino2: Tandem Activation Domains with Properties Similar to the Yeast Gcn4 Activator. Molecular and Cellular Biology 38(10).

63. Schulz S, et al. (2015) Elucidation of sigma factor-associated networks in Pseudomonas aeruginosa reveals a modular architecture with limited and function-specific crosstalk. PLoS pathogens 11(3):e1004744.

64. Battesti A, Majdalani N, & Gottesman S (2015) Stress sigma factor RpoS degradation and translation are sensitive to the state of central metabolism. Proceedings of the National Academy of Sciences of the United States of America 112(16):5159–5164.

65. Schuster M, Hawkins AC, Harwood CS, & Greenberg EP (2004) The Pseudomonas aeruginosa RpoS regulon and its relationship to quorum sensing. Molecular Microbiology 51(4):973–985.

66. Wassarman KM & Storz G (2000) 6S RNA Regulates E. coli RNA Polymerase Activity. Cell 101:11.

67. Bordes P, et al. (2003) DNA supercoiling contributes to disconnect σ^S^ accumulation from σ^S^- dependent transcription in Escherichia coli. Molecular Microbiology 48(2):11.

68. Meyer AS & Grainger DC (2013) The Escherichia coli Nucleoid in Stationary Phase. Adv Appl Microbiol 83:69–86.

69. Artsimovitch I & Henkin TM (2009) In vitro approaches to analysis of transcription termination. Methods 47(1):37–43.

