## Supplementary material for "The dormancy-specific regulator, SutA, is intrinsically disordered and modulates transcription initiation in *Pseudomonas aeruginosa*"

**Supplementary Information Text**

**EXTENDED MATERIALS AND METHODS**

**Media and Growth Conditions.** All cultures were grown at 37 °C with shaking unless otherwise noted. Liquid media were LB (5 g yeast extract, 10 g tryptone, 10 g NaCl per liter), Terrific Broth (TB) (24 g of yeast extract, 20 g of tryptone, and 4 mL of glycerol per liter, buffered to pH 7.0 with 18.9 mM potassium phosphate ), or M9 minimal salts medium for isotopically labeled protein expression (6 g/L sodium phosphate dibasic, 3 g/L potassium phosphate monobasic, 0.5 g/L NaCl, 1 g/L $^{15}NH_4Cl$ (Cambridge Isotope Laboratories, Cambridge MA), 2.5 g/L $^{13}C$ glucose(Cambridge Isotope Laboratories)) supplemented with 100 µg/ml carbenicillin (Gold Bio), 1 mM $MgSO_4$, 300 µM $CaCl_2$ and trace metals (7.5 µM $FeCl_2 \cdot 4H_2O$, 0.8 µM $CoCl_2 \cdot 6H_2O$, 0.5 µM $MnCl_2 \cdot 4H_2O$, 0.5 µM $ZnCl_2$, 0.2 µM $Na_2MoO_4 \cdot 2H_2O$, 0.1 µM $NiCl_2 \cdot 6H_2O$, 0.1 µM $H_3BO_3$, 0.01 µM $CuCl_2 \cdot 2H_2O$)

**Strain and plasmid construction**. See Table 2 (strains and plasmids) and Table 3 (primers) for relevant details. In general, standard methods were used for plasmid and strain construction. For comparing the effects of SutA overexpression in *E. coli* to its effects in *P. aeruginosa*, either the expression plasmid (from strain DKN1640) or the empty vector (from strain DKN548 (1)) was transformed by electroporation into either the *P. aeruginosa* Δ*sutA* strain (DKN1625) or *E. coli* MG1655 (DKN81). For overexpression and Ni-NTA purification of SutA, a plasmid in which the an HA-tagged *sutA* gene had been amplified and cloned into the multiple cloning site of pQE-80L (Qiagen) between the BamHI and HindIII restriction sites (DKN1643) was amplified using outward-directed primers flanking the sequence for the HA tag (not amplifying it) and encoding the TEV cleavage site. The PCR product was phosphorylated and subjected to a blunt end ligation to generate the plasmid encoding a 6His-TEV-SutA construct, and this was transformed into BL31 DE3 cells to generate strain DKN 1697. This construct was subjected to site directed mutagenesis using outward-facing primers encoding the desired changes to generate all of the SutA variant constructs used in this study (DKN1879-DKN1892), except SutA 46-101 (DKN1878). Sequences of SutA46-101, DksA (DKN1893) and *rpoB* β1 (DKN1895) were cloned out of genomic DNA from *P. aeruginosa* UCBPP-PA14 and into the pQE-80L plasmid from strain DKN1697, replacing the SutA sequence but retaining the TEV cleavage site, using Gibson assembly (2). The *rpoD* and *rpoS* sequences were cloned from *P. aeruginosa* gDNA and into the pET15b vector (DKN1901 and DKN1894, respectively), as expressing *rpoD*from pQE-80L proved somewhat toxic to *E. coli*. The $\sigma^{70}\Delta$171-214 construct (DKN1902) was generated using outward facing primers and blunt-end ligation of the plasmid from strain DKN1901. Fragments of β to use as standards in the affinity cleavage experiment were cloned from *P. aeruginosa* gDNA into pQE-80L, removing the sequence for the 6xHis affinity tag and TEV cleavage site, by Gibson assembly (DKN1896-DKN1900). *Rrn* template sequence was cloned from *P. aeruginosa* gDNA into the pUC18 vector (DKN1903).

**Protein purification.**

**RNAP:** RNAP was purified from the *P. aeruginosa ΔsutA* strain essentially as previously described ((3) and references therein). Briefly, cells were grown in 6 L of TB to an $OD_{600}$ of approximately 1.0. Cells were washed with TBS and pellets were frozen at -80 °C. Cell pellets were resuspended in 90 mL RNAP lysis buffer (50 mM Tris pH 8.0, 100 mM

NaCl, 1 mM EDTA, and cOmplete complete Ultra EDTA-free protease inhibitor tablets (Roche)) containing 40 Kunitz units DNAseI and cells were lysed by passage through an EmulsiFlex-C3 (Avestin). Lysates were clarified by centrifugation at 12,000 xg, and nucleic acids and acidic proteins were precipitated by addition of a 10% polyethyleneimine (polymin P; Sigma-Aldrich) solution at pH 7.9 to a final concentration of 0.5%. Precipitated protein was pelleted, washed with TGEB (10 mM Tris pH 8.0, 5% glycerol, 0.1 mM EDTA, 10 mM β-mercaptoethanol) plus 0.3 M NaCl, and the RNAP fraction was eluted with TGEB plus 1 M NaCl. Residual polymin P was removed by ammonium sulfate precipitation (2M). The ammonium sulfate pellet was resuspended in TGEB and loaded onto a 50 mL Heparin Sepharose 6 Fast Flow column (GE Healthcare). The column was washed with 2 column volumes of TGEB plus 0.3 M NaCl, and RNAP was eluted with a step to TGEB plus 0.6 M NaCl. The elution fraction was precipitated with 2 M ammonium sulfate, and resuspended into approximately 1 mL of TGEB plus 0.5 M NaCl. Low molecular weight contaminants were removed via size exclusion chromatography on a HiPrep 16/60 Sephacryl S-300 HR column (GE Healthcare). Fractions containing RNAP were diluted in TGEB to a final NaCl concentration of 0.3 M and loaded onto a HiTrap Q HP 5 mL column (GE Healthcare). RNAP was eluted into TGEB with a gradient between 0.3 M and 0.5 M NaCl over 20 column volumes. RNAP was dialyzed into RNAP storage buffer (20 mM Tris-HCl pH 8.0, 0.1 mM EDTA, 10 mM β-mercaptoethanol, 100 mM NaCl, 20% glycerol), concentrated to 1.4 mg/mL and frozen at -80 °C. The total yield was approximately 2.9 mg of high purity core enzyme.

**6xHis-tagged proteins:** For all tagged proteins, the following central steps were in common, and initial protein expression and lysis steps, plus additional purification steps specific to each protein are detailed below: Soluble protein was mixed with His-Pur Ni-NTA beads (Thermo Scientific or Clontech) in batch and binding was allowed to occur for 1h at 4 °C. Beads were washed three times with lysis buffer containing 20 mM imidazole and eluted three times with lysis buffer containing 250-500 mM imidazole. Eluents were combined, loaded onto an Amicon 3 or 10 kDa centrifugal filter (EMD Millipore), and buffer exchanged to TEV-digestion buffer (50 mM Tris pH 8.0, 0.5 mM EDTA, and 1 mM DTT). The 6xHis-tag was cleaved by addition of His-tagged TEV protease in a 1:50 mass ratio and incubation overnight at 4 °C. The digested sample was reapplied to His-Pur Ni-NTA, and washed with lysis buffer containing 20 mM imidazole; the protein of interest remained unbound or was eluted in this wash step, while the cleaved peptide tag and His-tagged TEV protease remained bound to the resin. The cleaved protein product includes the native protein sequence with an additional N-terminal serine (or glycine for $\sigma^{70}$purified from the pET15b vector).

**SutA (unlabeled):** Strain DKN1697 was grown with 200 µg/ml ampicillin. A 20 mL culture grown overnight in LB was distributed between two flasks, each containing one liter of 2xYT and grown at 37 °C to $OD_{600}$=0.6. Protein expression was induced by addition of 1 mM isopropyl β-D-1-thiogalactopyranoside (IPTG) and expression was allowed to continue for 4 hr. Cells were pelleted and frozen at -80 °C. Pellets were resuspended in lysis buffer (40 mM $NaH_2PO_4$, 300 mM NaCl, pH 8) containing 5 mM imidazole, 1 mg/mL lysozyme, and cOmplete mini protease inhibitor, EDTA free and lysed by probe sonication. The lysate was treated with Benzonase Nuclease on ice for 30 min and centrifuged. Following TEV cleavage of the SutA protein, the protein was concentrated on an Amicon Ultra-15 centrifugal filter, applied to a Superdex 75 10/300 column, buffer exchanged to SutA storage buffer (25 mM Tris pH 8, 100 mM NaCl, 20% glycerol, and 2 mM β-mercaptoethanol), and stored at -80 °C.

**SutA 46-101 ($^{15}N^{13}C$):** Strain DKN1878 was grown overnight in 10 ml LB and then split between two baffled flasks containing M9 minimal salts medium. Cultures were grown at 37 °C until they reached mid-exponential phase (8 hrs) and then protein expression was induced by adding IPTG to a final concentration of 1 mM. Cells were harvested after 5 hr of induction and frozen at -80 °C. Pellets were resuspended in lysis buffer (50 mM sodium phosphate, pH 8.0, 300 mM NaCl, 5 mM imidazole) plus 20 Kunitz units DNaseI and EDTA-free cOmplete mini protease inhibitor tablets (Roche) and lysed by passage through an EmulsiFlex-C3. Following TEV cleavage, the protein was concentrated and loaded onto a Hi-Load 16/600 Superdex75 pg size exclusion column, buffer exchanging into the NMR buffer containing 20 mM sodium phosphate pH 7.0 and 100 mM sodium chloride.

**SutA WT ($^{15}N^{13}C$):** Strain DKN1697 was grown, protein expression induced, and cells lysed as described for the SutA 46-101 ($^{15}N^{13}C$) protein. As an additional purification step following TEV cleavage, the protein was concentrated and buffer exchanged into a buffer containing 20 mM N-methylpiperazine, pH 5.0, and 100 mM NaCl and loaded onto a 5 ml HiTrap Q Sepharose fast flow anion exchange column (GE Healthcare Life Sciences). The protein was eluted with a 20 column-volume gradient to 600 mM NaCl and then was concentrated to 1 ml before size exclusion chromatography as described for the SutA 46-101 ($^{15}N^{13}C$) protein.

**SutA WT ($^{15}N$):** Protein was produced and purified as described for the SutA WT ($^{15}N^{13}C$) protein, except glucose with the natural carbon isotope ratios was used at 4 g/L.

**SutAΔN unlabeled:** Strain DKN1879 was grown overnight in 5 ml LB then diluted 1:200 into TB plus 100 µg/ml carbenicillin and grown at 37 °C. Expression was induced when the culture reached mid-exponential phase with 1 mM IPTG and cells were harvested after 4 hrs of induction. Lysis and purification steps were the same as described for the SutA 46-101 ($^{15}N^{13}C$) protein, and the final protein storage buffer was 25 mM Tris pH 8, 100 mM NaCl, 20% glycerol, and 2 mM β-mercaptoethanol.

**SutAΔC unlabeled:** Strain DKN1880 was used, and all expression and purification steps were the same as for the unlabeled SutAΔN protein.

**SutAΔN ($^{15}N$):** Strain DKN1879 was used, and all expression and purification steps were the same as for the SutA 46-101 ($^{15}N^{13}C$) protein, except glucose with the natural carbon isotope ratios was used at 4 g/L.

**SutAΔC ($^{15}N$):** Strain DKN1880 was used, and all expression and purification steps were the same as for the SutA 46-101 ($^{15}N^{13}C$) protein, except glucose with the natural carbon isotope ratios was used at 4 g/L.

**SutA BPA variants:** *E. coli* BL21 DE3 was co-transformed with pEVOL-pBpF (4) and the plasmids from strains DKN1881-DKN1889 (pQE80L-6xHis-TEV-SutA amber mutants). Approximately 20 colonies were scraped from the agar plate and grown at 33 °C in LB to OD600 = 0.6. Cultures were treated with 1 mM BPA (Iris-Biotech, Marktredwitz, Germany) and 1 mM IPTG and incubated in the dark for 20 h. Cells were pelleted and frozen at -80 °C. Pellets were resuspended in lysis buffer (40 mM NaH2PO4, 300 mM NaCl, pH 8) containing 5 mM imidazole, 1 mg/mL lysozyme, and cOmplete mini protease inhibitor, EDTA free and lysed by probe sonication. The lysate was treated with Benzonase Nuclease on ice for 30 min and centrifuged. Following TEV cleavage, SutA fractions were pooled and loaded onto an Amicon 10 kDa centrifugal filter, and buffer exchanged to SutA storage buffer (25 mM Tris pH 8, 100 mM NaCl, 20% glycerol), and stored at -80°C.

**SutA FeBABE variants:** Strains DKN1890-DKN1892 were used. Expression and purification steps were the same as for the SutAΔN unlabeled protein.

**β1:** Strain DKN1895 was used. An overnight culture was grown in LB plus 100 µg/ml carbenicillin and 10 µg/ml gentamicin at 37 °C. The culture was diluted 1:100 into TB and grown for 3 hrs without antibiotics at 30 °C. The culture was cooled to 13 °C and expression was induced for 24 hrs with 400 µg/ml IPTG. Cell pellets were collected and frozen at -80 °C. Pellets were resuspended in a modified RNAP purification buffer (20 mM Tris pH 7.6, 5% glycerol, 3 mM 2-mercaptoethanol, 200 mM NaCl, 10 mM imidazole) plus 20 Kunitz units DNaseI and EDTA-free cOmplete mini protease inhibitor tablets (Roche) and lysed by passage through an EmulsiFlex-C3 (Avestin). Much of the expressed protein was not soluble, but the soluble fraction was bound in batch to Ni-NTA beads, washed, eluted, and its TEV tag cleaved as described above, except TEV was used at a mass ratio of 1:25. Following TEV cleavage, the protein was concentrated to 1 ml in SEC buffer (30 mM Tris pH 7.6, 120 mM NaCl, 0.1 mM EDTA, 5% glycerol, 2.1 mM 2-mercaptoethanol) and passed over a HiLoad 16/600 Superdex 200 pg column. Fractions containing the protein of interest were collected and concentrated, and the glycerol concentration was brought to 20% before storage at -80 °C.

**$\sigma^{70}$:** Strain DKN1901 was grown overnight in LB containing 100 µg/ml carbenicillin, then diluted 1:1000 into TB. After 4 hr growth, the culture was cooled to 16 °C and expression was induced with 400 µg/ml IPTG for 18 hr. Cell pellets were collected, and lysis and purification was carried out as described for the RpoB B1 protein.

**$\sigma^S$:** Strain DKN1894 was used. Expression and purification were carried out as described for the unlabeled WT SutA, except TEV cleavage was not performed and an addition size exclusion step using a Superdex 200 column was added. Final protein storage buffer included 25 mM Tris pH 8, 100 mM NaCl, 20% glycerol, and 2 mM β-mercaptoethanol.

**DksA:** Strain DKN1893 was used. Expression and purification were carried out as described for the unlabeled WT SutA.

**$\sigma^{70}$ Δ171-214:** Strain DKN1902 was used. Expression and purification steps were carried out as for the full-length$\sigma^{70}$.

**FeBABE conjugation.** FeBABE was conjugated to the purified SutA S2C, S32C, and S98C proteins as described (5). Briefly, the purified proteins were de-metallated and fully reduced by incubating in a buffer containing 20 mM sodium phosphate pH 7.0, 100 mM NaCl, 20 mM EDTA, and 1 mM DTT overnight. They were then buffer exchanged into conjugation buffer (20 mM MOPS pH 8.0, 100 mM NaCl, 2 mM EDTA, 5% glycerol) using Amicon 3 kDa centrifugal filters, with care taken to reduce DTT concentrations to sub-micromolar levels. The concentration of free cysteines was measured using Ellman's reagent (see below) and this measurement was used as the SutA concentration for the FeBABE variant proteins. SutA concentrations in the labeling reactions were 25-30 µM. The FeBABE reagent (Dojindo Molecular Technologies, Rockville MD) was dissolved in DMSO to 20 mM and added to a final concentration of 300 µM in a reaction volume of 1 ml. The reaction was incubated for 1 hr at 37°C and then quenched by dilution of the FeBABE reagent via dialysis into protein storage buffer (20 mM Tris, pH 7.6, 100 mM

NaCl, 20% glycerol, 0.1 mM EDTA). The concentration of free cysteines was measured again using Ellman's reagent to determine the efficiency of FeBABE conjugation, which was as follows: S2C variant (N-Fe): 57.4% labeled; S32C variant: 37.9%; S98C variant (C-Fe): 76.3%.

**Protein quantification.** As we characterized SutA, it became clear that standard methods for protein quantification were very inaccurate for this protein, and that the degree and direction of the inaccuracy was different for the N- and C-terminal SutA mutants. This is likely due to the unusual amino acid composition of SutA compared to the bovine serum albumin (BSA) standard that is usually used for calibration in Bradford and BCA assays. We found that the Bradford assay (and coomassie staining of gels) greatly underestimated SutA concentration (likely due to a lack of aromatic amino acids and overabundance of acidic amino acids), and that the ΔC mutation exacerbated this problem by removing one of the two aromatic amino acids. The BCA assay slightly overestimated SutA concentration, likely due to the high accessibility of protein backbone, and this was also exacerbated in the ΔC protein, perhaps because a higher percentage of the remaining protein was the completely unstructured N-tail. Accordingly, we quantified the concentrations of our unmodified SutA proteins using total acid hydrolysis, derivatization of the resulting free amino acids, and HPLC as described below (6). The FeBABE SutA variants were quantified using Ellman's reagent to measure their free cysteines (one per protein) before FeBABE conjugation as described below. The BPA SutA variants were quantified using the BCA assay (Thermo Fisher) according to the manufacturer's instructions, which was reasonably accurate for the full-length protein. All other proteins

(RNAP core enzyme and β1 fragment, σ factors, and DksA) were quantified using the Quick Start Bradford Protein Assay (Bio-Rad) with BSA as a standard.

**Ellman's reagent assay:** Ellman's reagent (5,5-dithio-bis-(2-nitrobenzoic acid)) (Thermo Fisher) was dissolved in FeBABE conjugation buffer at 4 mg/ml. This stock was further diluted 1:50 into the buffer containing the protein to be assayed and distributed to the wells of a 96-well plate at 200 µl per well. 20 µl protein sample or cysteine hydrochloride monohydrate calibration standard was added, and absorbance at 412 nm was measured on a plate reader after incubation for 15 min at room temperature.

**Amino acid hydrolysis and HPLC:** SutA proteins prepared for NMR, which were stored in 20 mM sodium phosphate, 100 mM NaCl buffer without glycerol, were used for quantification by amino acid hydrolysis. Subsequently, the concentrations of the glycerol stocks of the corresponding unlabeled proteins were determined by quantifying the intensity of Coomassie staining on an SDS-PAGE gel of the quantified NMR protein stocks and the glycerol stocks, run side by side. Vacuum hydrolysis of the SutA protein stocks was carried out by continuous boiling for 24 hr at 105 °C in 6 N HCl in a Thermo Scientific Pierce 1 ml vacuum hydrolysis tube (Thermo Fisher), according to the manufacturer's instructions. After hydrolysis, the protein was dried *in vacuo* and resuspended in 100 µl 150 mM $NaHCO_3$ pH 9.0. 100 µl 15 mM dabsyl chloride (Sigma) in acetonitrile was added and the samples were incubated at 70 °C for 15 min. The reaction was quenched by the addition of 800 µl of a 1:1 mixture of ethanol and water. Debris were removed by centrifugation at top speed in a microfuge and the sample was transferred to an HPLC vial. 5 µl of the sample was injected onto a Waters Alliance HPLC system, composed of an e2695 separation module, 2998 PDA detector, and

Acquity QDa detector, and fitted with a 3x100 mm XBridge BEH C18 reversed-phase chromatography column, 2.5 µm particle size. Buffer A contained 0.04% $NH_4OH$ in water, and Buffer B contained 0.04% $NH_4OH$ in acetonitrile. Each sample was loaded onto the column in a mixture of 8% buffer B and 92% buffer A, and a gradient from 8-30% buffer B was run over 40 min, followed by a gradient from 30-90% buffer B over 10 min. The column was then cleaned for 2 min with 90% buffer B, and returned to 8% buffer B over 8 min. A 2.5 mM amino acid standard mix in 0.1 N HCl (Sigma) was subjected to the same hydrolysis and derivatization protocol and used to calibrate amino acid peak areas. The identity of each peak was confirmed by mass spectrometry. Quantifications of alanine, glycine, isoleucine, leucine, lysine, methionine, phenylalanine, proline, and serine were averaged for each sample to estimate the concentration of the SutA variant.

**NMR experiments**

Proteins were purified as described above. Except where noted, protein concentrations were 300 µM and the buffer contained 20 mM sodium phosphate, pH 7.0, 100 mM sodium chloride, and 10% $D_2O$.

**46-101:** 2D and 3D NMR spectra were collected on a Varian Inova 600 MHz NMR with a triple resonance inverse probe running VnmrJ 4.2A. The optimal temperature for minimizing the linewidth of $^{15}N$ HSQC peaks was found to be 7 °C. Although SutA was stable in solution at fairly high concentration at a range of temperatures, the peaks showed concentration-dependent broadening that was only alleviated by decreasing the concentration and acquiring the spectra below ambient temperature. The following spectra were acquired: $^{15}N$ HSQC, $^{13}C$ HSQC, HNCO, HNCA, HNCACB, CBCACONH,

HNCOCA, HNCACO, CCONH, and $^{15}N$ HSQC experiments modified for measurement of $T_2$ and of $^{15}N$-$^{1}H$ NOE. These experiments were all done with standard Varian/Agilent pulse programs included in the Biopack extension of VnmrJ. The processed spectra were imported into the CcpNmr Analysis program (7), and Assign-derived peak lists from the spectra were submitted to the PINE web server assignment program maintained by NMRFAM at the University of Wisconsin, pine.nmrfam.wisc.edu (8). Assignments proposed by the PINE output were validated or corrected in the Analysis software.

**Full-Length SutA:** Spectra were acquired at 7 °C on a Bruker AV III 700 MHz spectrometer with a TCI cryoprobe running Topspin 3.2. The spectra ($^{15}N$ HSQC, $^{13}C$ HSQC, HNCACB, and CBCACONH) were all acquired with standard Bruker pulse programs. $^{15}N$ HSQC experiments modified for measurement of $T_2$ and of $^{15}N$-$^{1}H$ NOE were performed on a Varian Inova 600 MHz NMR with a triple resonance inverse probe running VnmrJ 4.2A, at 7 °C, with standard Varian/Agilent pulse programs included in the Biopack extension of VnmrJ. Standard $^{15}N$ HSQC spectra were also collected at 7 °C, 16 °C and 25 °C. The spectra were imported into CcpNmr Analysis and partially assigned via the PINE web server as described previously.

**Additional $^{15}N$ HSQC experiments:** $^{15}N$ HSQC spectra for the SutA ΔN and SutA ΔC SutA proteins were collected on a Varian Inova 600 MHz NMR with a triple resonance inverse probe running VnmrJ 4.2A, with standard Varian/Agilent pulse programs included in the Biopack extension of VnmrJ, to test whether the truncations influenced the overall structure of the protein.

**Stretched gel preparation for residual dipolar coupling measurements:** $^{15}N^{13}C$-labeled SutA was embedded in a stretched polyacrylamide gel using the “Gel NMR

Starter Kit" (cat. #NE-373-B-5.4/4.2, New Era, Vineland NJ), according to the manufacturer's instructions. Briefly, a cylindrical 8% polyacrylamide gel of about 300 µl, with a diameter of 5.4 mm (29:1 acrylamide:bisacrylamide ratio) was prepared. After polymerization, the gel was dialyzed 3 times against nanopure water, then dried overnight at 37 °C, and then returned to the cylindrical chamber in which it was cast. 300 µl $^{15}N^{13}C$-labeled SutA at a concentration of 300 µM in a buffer containing 20 mM sodium phosphate, pH 7.0, 100 mM sodium chloride, and 10% $D_2O$ was added to the dried gel and allowed to soak into it overnight at room temperature. The SutA-impregnated gel was then pushed into an NMR tube with a diameter of 4.2 mm, resulting in its stretching. Spectra were collected on a Varian Inova 600 MHz NMR with a triple resonance inverse probe running VnmrJ 4.2A. To extract $^{1}J(^{15}N, ^{1}H)$ coupling constants, the pulse sequence gNhsqc_IPAP was used to acquire the in-phase and antiphase spectra alternately. The sum and difference spectra were generated in VnmrJ with appropriate 2D transform coefficients and imported into CcpNmr Analysis for overlay with the conventional $^{15}N$ HSQC spectrum.

**NMR binding experiment:** $^{15}N$-labeled SutA and β1 fragment purified as described above were buffer exchanged into 20 mM sodium phosphate, pH 7.0, 100 mM sodium chloride, at an approximate concentration of 30 µM each. To increase the chances that most of the SutA would be bound to β1 fragment, the mixture was run over a HiLoad Superdex 200 pg size exclusion column (GE Healthcare Life Sciences, Marlborough MA) and fractions representing the complex were retained and concentrated to 270 µl before adding $D_2O$ to 10%. The final concentration of the complex was approximately 25 µM. In addition, $^{15}N$-labeled SutA was mixed with $\sigma^S$ at 50 µM each and buffer exchanged into 20 mM sodium phosphate, pH 7.0, 100 mM sodium chloride, and 10% $D_2O$. $^{15}N$ HSQC spectra were acquired on a Bruker 800 MHZ AV III HD spectrometer with a TCI cryoprobe at 25 °C using the standard Bruker pulse sequence hsqcetfpf3gpsi.

**Data analysis:** Secondary shifts were calculated by the TALOS software package as part of the PINE output. RDC values were evaluated manually by comparing the overlaid sum and difference spectra in the CcpNmr Analysis Suite, and the presence or absence of a peak in the positive ($^{1}H$-$^{15}N$) NOE was also evaluated manually for each assigned residue in the CcpNmr Analysis Suite. $R_2$ values were calculated by fitting a single exponential to the series of peak integral values collected with different $T_2$ relaxation times for each assigned residue. To generate structural models based on the chemical shift and RDC values we collected, these values were uploaded to the Robetta Fragment Server (9), and 3- and 9-residue fragment libraries were picked. Each library contained 200 fragments per SutA amino acid position. Using these fragment libraries, 16,000 decoy structures were generated using the PyRosetta suite (10), following a folding protocol based on the PyRosetta folding tutorial published by the Gray lab (11). Briefly, the SutA protein sequence was set to a linear structure, then 1000-1500 cycles of fragment insertion and energy minimization were performed to generate each decoy. Each cycle consisted of 3 short fragment (3 residues) and 1 long fragment (9 residues) insertions, followed by a low-resolution Monte Carlo scoring. As is perhaps unsurprising for a protein that has large regions of intrinsic disorder, the decoys did not converge to a single family of lowest-energy structures. We calculated the RMSD for each decoy compared to an ab initio structural prediction for SutA that was produced by the Robetta Server (9). In general, decoys with lower RMSDs compared to this ab initio prediction also contained some version of the α helix that is supported by our NMR data; some other decoys (and some with the lowest energy scores) did not have the α helix. We arbitrarily chose several decoys to show a range of conformations that SutA might adopt; the strongest predictions of our NMR data are that residues 56-76 adopt an α-helix secondary structure and that the N- and C-tails are disordered, and all of the chosen models conform to those predictions. To color the model shown in Figure 1D according conservation, the alignment shown in Figure 1 Supplement 1 (generated using the MEGA6 software suite (12), with gaps in the *P. aeruginosa* UBCPP-PA14 sequence removed, and the alignment visualized using the Jalview2 applet(13)) was opened in Chimera (14), and the "Render by conservation" function was used.

***In vitro* transcription experiments**

Experiments were carried out broadly as described in (15). In general, RNAP holoenzyme was prepared by mixing core enzyme with a 3-fold ($\sigma^{70}$) or 5-fold ($\sigma^{s}$) excess of σ factor and incubating for 15 min at 37 °C. dsDNA templates were prepared by PCR from a plasmid carrying the relevant promoter sequences, using the Kappa high-fidelity hot-start 2x master mix according to the manufacturer's instructions (see strain and primer tables for plasmid and primer details). PCR products were checked by electrophoresis on 2% agarose gels to ensure that they consisted of a single product, purified from primers and residual dNTPs using the DNA Clean and Concentrator kit (Zymo Research, Irving CA), and quantified by NanoDrop (Thermo Fisher). The *rrn* bubble template was prepared by annealing the template strand and non-template strand oligos as follows: 80-mer oligos (Integrated DNA Technologies) were resuspended at a concentration of 100 µM in 0.1x TE and mixed together in 10X annealing buffer to give final concentrations of 45 µM duplex, 10 mM Tris-Cl ,100 mM NaCl, and 1 mM EDTA, then heated to 95 °C for 5 min and allowed to cool from 95 °C to 70 °C at a rate of 0.1 °C/ second, incubated at 70 °C for 20 min, then allowed to cool to 22 °C at a rate of 0.1 °C/second. All pre-incubations and reaction incubations took place at 37 °C, and all reactions used TGA buffer (20 mM Tris-acetate pH 8.0, 2 mM Na-acetate, 2 mM Mg-acetate, 4% glycerol, 0.1 mM DTT, 0.1 mM EDTA). Water used in reaction and running buffer preparation was treated with diethyl pyrocarbonate (DEPC). Reactions were quenched with an equal volume of urea stop buffer (8 M urea, 10 mM EDTA, 0.8x TBE, 2 mg/ml bromophenol blue, 2mg/ml xylene cyanol FF, 2 mg/ml amaranth), and heated to 95 °C for 2 min immediately before gel loading. 20% acrylamide denaturing Urea-TBE gels were prepared using the Sequa-gel system (National Diagnostics) according to the manufacturer's instructions except TBE was added to 0.5x instead of 1x. A 60-well comb was used and gels were run using the Owl S3 vertical sequencing gel system (Thermo Fisher). 2 µl sample was loaded per lane. After electrophoresis, one glass plate was removed and the gel was covered with plastic wrap and exposed directly to the phosphorimager screen (Molecular dynamics) for 12-48 hr.

**Single turnover initiation experiments:** For SutA titrations, reactions were assembled as follows: RNAP holoenzyme (20 nM final concentration), DNA template (15 nM final concentration), TGA buffer, and water were mixed in a volume of 3 µl and added to 1 µl SutA (at 5x the final concentration) or storage buffer on ice. These 4 µl reactions were incubated for 6 min to allow open complex to form. 1 µl NTP mix (375 µM initiating dinucleotide, 250 µM each NTP not carrying $^{32}P$ label (ATP, UTP, and either CTP or GTP), 100 µM cold NTP of the same type as that carrying the label (either CTP or GTP),

0.75 µCi $\alpha^{32}$P GTP or CTP (3000 Ci/mmol, 10 mCi/ml, Perkin Elmer, Waltham MA), and 100 µg/ml heparin) was added and transcription was allowed to continue for 8 minutes before reactions were quenched. Initiating nucleotides were CpU for the *rrn* promoter and ApC for the *bcn* promoter (IBA Lifesciences, Göttingen, Germany). The final NaCl concentration in these reactions (due to NaCl in protein storage buffers) was 26 mM for the *rrn*. For the iNTP titration experiments, no dinucleotides were included, and instead the NTP mix contained 50, 500, or 5000 µM CTP and UTP (for the 10, 100, and 1000 µM iNTP conditions), 250 µM ATP, 100 µM GTP, and 0.75 µCi $\alpha^{32}$P GTP per 1 µl NTP mix. For the DksA/ppGpp experiments, 0.5 µl 5 µM SutA (or 0.5 µl storage buffer) and 0.5 µl of a mixture containing 2.5 µM DksA and 25 µM ppGpp (Sigma) in storage buffer (or 0.5 µl storage buffer) were distributed to tubes. The remainder of the experimental set-up was the same as for the SutA titration experiments.

**Open complex stability assays:** A 7x reaction master mix containing RNAP holoenzyme (for 20 nM final concentration in transcription reactions) template (15 nM final), SutA at the indicated concentration, or storage buffer and water in a volume of 27.5 µl was mixed on ice. 1 µl NTP mix (at the same concentrations as described for single turnover reactions, but without heparin) was distributed to each of 6 reaction tubes. The reaction master mix was incubated for 6 min to allow open complex to form, and then 0.5 µl of heparin at 1.5 mg/ml was added. Immediately, 4 µl of the master mix was removed and added to 1 µl NTP mix for the time 0 point. At the indicated time points after the addition of heparin, additional 4 µl aliquots were removed and added to tubes containing 1 µl NTP mix. Each reaction was quenched 8 min after mixing the reaction mix with the NTP mix.

**Gel image acquisition and analysis:** Phosphorimager screens were scanned on a Typhoon FLA 9000 gel imaging system (GE Healthcare Life Sciences), at the maximum PMT setting and with each pixel representing 200 µm. Images were analyzed using the gel lane analysis tool of the FIJI open-source image analysis suite (16). First, images were rotated, background-subtracted, and contrast-adjusted (ensuring that no pixels were saturated), then pixel densities in the relevant regions of each lane were plotted, and the areas under each peak quantified. For the SutA titration experiments, 2-3 bands spanning a length range of about 4-8 nucleotides represented the run-off transcripts (RNAP terminates inefficiently on a linear transcript, sometimes producing multiple bands), and all of the major bands in this range were quantified (the ratios of each band to the total were not generally affected by SutA). The most prominent higher band likely represents the product of transcription initiating at one end of the linear transcript and running to the other end, and was ignored. For the iNTP titrations, the major products were the same as those seen for initiation with the CpU dinucleotide, but at the highest [iNTP], additional bands within the 8 nucleotide range appeared, and all bands in this range were quantified. For initiation on the bubble template, abortive products were much more prominent, and their relative abundances were affected by the σ used. In order to calculate a value representing the number of initiation events, all major bands in each lane were quantified and the signal intensity for each band was divided by the number of G bases in the sequence corresponding to that band to obtain a value proportional to the number of transcripts represented by the band. The sum of the values for each band in the lane was used as a measure of the total number of transcripts initiated. Whenever possible, values were normalized and compared within the same gel. Where comparisons across gels were necessary, values from each gel were normalized to the values obtained for reactions containing 0 nM SutA and E$\sigma^{70}$ on that gel.

**Transcription start site mapping:** *P. aeruginosa* UCBPP-PA14 culture was collected in mid-exponential, early stationary, and late stationary phases, cells were pelleted, and pellets were frozen in liquid nitrogen. RNA was extracted from the pellets using the RNeasy kit (Qiagen, Hilden, Germany) according to the manufacturer's instructions. Genomic DNA was depleted using the Turbo DNA-free kit (Ambion/Invitrogen, Carlsbad CA) according to instructions. cDNA corresponding to the 5' ends of nascent rRNA transcripts was generated by reverse transcription using 10 µg total RNA, 4 pmol rRNA-specific primer, 500 µM dNTPs, 5µM DTT, 1x reverse transcriptase buffer, and 300 units SuperScript reverse transcriptase in a 40 µl reaction. Primer binding was allowed to occur for 5 min at 65 °C, then the reverse transcriptase was added and the reaction allowed to proceed for 45 min at 55°C, then the reaction was stopped by incubation for 15 min at 70 °C. 2 units RNaseH were added and the reactions were incubated at 37 °C for 20 min to degrade RNA-DNA hybrids, and the cDNA was cleaned up using the Qiaquick PCR clean-up kit (Qiagen). Poly-T tails were added to the 3' ends of the cDNA using terminal transferase (Promega) according to instructions. The resulting T-tailed cDNA was then used as a template in a first round PCR reaction with a primer against the rRNA transcript and one against the poly-T tail that adds an additional specific sequence. This PCR product was then used as the template in a second round PCR reaction with primers against the rRNA transcript and the newly added specific sequence that was part of the primer in the first round PCR. Two different DNA polymerases were tried (GoTaq, Promega; or Q5, NEB), according to instructions, and gave similar results (Figure 2, – figure supplement 2). The resulting PCR products from the stationary phase time points were cloned into the pUC18 plasmid using Gibson assembly and approximately 40 individual clones were sequenced. Many of the products turned out to represent the site that is cleaved by RNAse III in the first step of 16S rRNA maturation, which is very similar in sequence and distance upstream of the mature 16S rRNA start (17) to the *E. coli rrn* RNAse III cleavage site (these correspond to the strong, lowest band in Figure 2 – figure supplement 2). However, a number of the products corresponded to the proximal putative transcription initiation site (second lowest band) and most of these initiated at the cytidine 8 bp downstream of the -10 motif, although a few also initiated at a cytidine 7 bp downstream of the -10 motif. Although we detected some fainter bands potentially corresponding to start sites further upstream, we were unable to recover any sequences corresponding to these start sites, even after the higher faint bands were gel-purified before cloning into pUC18. We also tested a promoter corresponding to  the next putative start site upstream of this start site *in vitro* and found that it drove initiation more weakly than the proximal start site (data not shown). Together, these data suggest that this proximal start site is the dominant one in *P. aeruginosa*, at least under the conditions we investigated.

**Cross-linking and Affinity Cleavage**

**$BS^3$ cross-linking:** Bis(sulfosuccinimidyl) suberate ($BS^3$) $d_0$ and $d_4$ isotopologs were purchased from Thermo Scientific. RNAP and SutA were mixed in a 1:10 molar ratio (0.5 µM RNAP, 5.0 µM SutA) in 10 mM HEPES pH 8, 100 mM potassium acetate and incubated on ice for 1.5 hr. Cross-linking was initiated by addition of 5 mM of a 4:1 molar ratio of $BS^3$ $d_0$:$d_4$ and the reaction was incubated on ice for 2 hr. Cross-linking was quenched by addition of ammonium bicarbonate to a final concentration of 50 mM. Proteins were digested in solution by incubation with 500 ng GluC overnight at 37 °C. Digestion was quenched by addition of 5% formic acid. Digested peptides were desalted by HPLC using a C8 microtrap (Optimize Technologies, Oregon City OR), using a gradient of buffer A: 0.2% formic acid in $H_2O$ and buffer B: 0.2% formic acid in acetonitrile) and concentrated *in vacuo*. Samples were resuspended in 0.2% formic acid and analyzed on the Orbitrap Elite Hybrid Ion Trap MS equipped with an Easy 1000 nanoUHPLC (Thermo Scientific). Solvent A consisted of 97.8% $H_2O$, 2% ACN, and 0.2% formic acid and solvent B consisted of 19.8% $H_2O$, 80% ACN, and 0.2% formic acid. Digested peptides were directly loaded at a flow rate of 500 nL/min onto a 16-cm analytical HPLC column (75 µm ID) packed in-house with ReproSil-Pur $C_{18}$AQ 3 µm resin (120 Å pore size, Dr. Maisch, Ammerbuch, Germany). The column was enclosed in a column heater operating at 45 °C. After 30 min of loading time, the peptides were separated with a 50 min gradient at a flow rate of 350 nL/min. The gradient was as follows: 2% B for five min, 2–40% B (60 min), and 100% B (10 min). The Orbitrap was operated in data-dependent acquisition mode to automatically alternate between a full scan ($m/z$=300–1600) in the Orbitrap and subsequent 5 HCD MS/MS scans in the Orbitrap. Normalized collision energy was 30% and activation time was 100 ms. Resolution on MS was set to 120,000 and MS/MS was 15,000. The experiment was performed with two replicates.

Raw files were first searched using MaxQuant to identify precursor mass pairs, differing by 4.02 Da, that represent cross-links made by both of the $BS^3$ linker isotopologs. Raw files were converted to peak lists with ProteoWizard (18) and subset for only those spectra that were identified as mass pairs. Subset peak lists were analyzed with Protein Prospector online, version 5.12.4, following reported protocols with modifications below (19). The protein database contained the sequences for purified SutA, RpoA, RpoB, RpoC, RpoD, and RpoZ. 80 peaks from each spectrum were searched using a tolerance of 10 ppm for precursor ions and 25 ppm for product ions. Enzyme specificity was GluC, and up to two missed cleavages per peptide were allowed. Carbamidomethylation of cysteines was specified as a constant modification, and protein N-terminal acetylation, oxidation of methionine, and dead-end modification with the cross-linker at lysine positions and protein N-termini were set as variable modifications. Additionally, incorrect monoisotopic peak assignments were considered as variable modifications. The analysis was run twice for each set of peak lists to search for both cross-linker isotopologs.

For cross-links detected between RNAP proteins, we used a reported structural model of the *E. coli* RNAP complex (PDB: 3LU0) to calculate the inter α-carbon distance between amino acids (20). We used this calculated distance as a metric to distinguish "quality" cross-links from all others. Based on the length of the linker, the maximum inter α-carbon distance between lysines cross-linked by BS3 is 24.6 Å, so we considered cross-links with distances near or below this value to be reasonable. Like the study by Trnka et al., we found Score Difference to be the best discriminant for making this distinction. A Score Difference cutoff of 8.0 (similar to the value of 8.5 found by Trnka et al.) separated high-distance and low-distance cross-links (Figure 3 supplement). The final criteria for assigning quality cross-links were: (i) found as a precursor mass pair and (ii) Score Difference greater than 8.0. These cross-links were aggregated to determine the number of spectra from each replicate and the maximum Score Difference for each amino acid linkage (Figure 3 supplement). To visualize cross-link spectra, peak lists subset for matched pairs were analyzed by StavroX (21) using the same settings described for Protein Prospector. The best spectra used to match the cross-links between SutA and RNAP are shown in Figure 3 supplement.

**BPA cross-linking for LC-MS/MS analysis:** 20 µl cross-linking reactions contained 500 nM core RNAP, 2 µM SutA (BPA54 or BPA84 variant), 100 mM NaCl, and TGA buffer (4% glycerol, 20 mM Tris-acetate pH 8.0, 2 mM sodium acetate, 2 mM magnesium acetate, 100 µM DTT, and 100 µM EDTA). Complexes were allowed to form for 6 min at 37 °C and were then UV-irradiated for 1 min at 1W/cm$^2$ using the Omnicure S2000 lamp (Excelitas, Waltham MA). Cross-linked complexes were dried *in vacuo*, resuspended in 40 µl 8 M urea and 100 mM Tris-HCl, reduced with 3 mM TCEP, alkylated with 10 mM iodoacetamide, digested with 100 ng lysyl endopeptidase for 4 hr, and then digested with 500 ng trypsin overnight in 2 M urea and 1 mM $CaCl_2$. Formic acid was added to 5% and then the sample was desalted by HPLC using a C8 microtrap (Optimize Technologies), with a gradient of buffer A: 0.2% formic acid in $H_2O$ and buffer B: 0.2% formic acid in acetonitrile), concentrated *in vacuo*, and resuspended in 0.2% formic acid. Samples were resuspended in 0.2% formic acid and run on the Q Exactive HF Orbitrap MS, equipped with an Easy 1200 nanoUHPLC (ThermoFisher Scientific). Solvent A consisted of 97.8% $H_2O$, 2% ACN, and 0.2% formic acid and solvent B consisted of 19.8% $H_2O$, 80% ACN, and 0.2% formic acid. Digested peptides were directly loaded at a flow rate of 220 nL/min onto a 20-cm analytical HPLC column (50 µm ID) packed in-house with ReproSil-Pur $C_{18}$AQ 1.9 µm resin (120 Å pore size, Dr.

Maisch, Ammerbuch, Germany). The column was enclosed in a column heater operating at 65 °C. After 45 min of loading time, the peptides were separated with a 60 min gradient at a flow rate of 220 nL/min. The gradient was as follows: 2–6% B (4 min), 6-25% B (41 min), 25-40% B (15 min), and 100% B (10 min). The Orbitrap was operated in data-dependent acquisition mode to automatically alternate between a full scan ($m/z$=300–1650) in the Orbitrap and subsequent 7 HCD MS/MS scans. Normalized collision energy was 28 and max injection time of 250 ms. Resolution on MS was set to 60,000 and MS/MS was 30,000. Raw files were converted to mzXML files by msConvert (22) and analyzed using StavroX (21) with a precursor and fragment ion tolerance of 5 ppm and a 1% FDR.

**FeBABE cleavage experiments:** FeBABE cleavage experiments were based on protocols described in (5).Our initial determination of SutA-FeBABE cleavage sites (as shown in Figure 3) utilized a large-format gel and Western blotting apparatuses (16x16 cm) to allow for higher resolution in calculating the cleavage site. 20 µl Reactions contained 250 nM RNAP (E, E$\sigma^S$, or E$\sigma^{70}$), 250 nM *rrn* template, 2 µM SutA (WT or FeBABE variant), 100 mM NaCl, in 1x TGA buffer (yielding a final glycerol concentration of 8% including enzyme storage buffers). Holoenzyme complexes were formed by mixing a 3-fold molar excess of $\sigma^S$ or $\sigma^{70}$ with E and incubating at 37 °C for 15 min. After assembling the rest of the reaction mixture, it was incubated at 37 °C for 10 min. to allow SutA and DNA-containing complexes to form, and then cleavage was initiated by the addition of 2.5 µl 50 mM sodium ascorbate, 10 mM EDTA then 2.5 µl 50 mM hydrogen peroxide (J.T. Baker Ultrex grade (Avantor, Radnor PA)), 10 mM EDTA.

Reactions were incubated for 7 min and then quenched by the addition of 8.3 µl 4x LDS loading buffer (Bio-Rad, Hercules CA).

**FeBABE protein cleavage reactions of open complexes:** Reactions containing different σ factors and promoter DNA were carried out on a smaller scale for SDS-PAGE and western blotting on mini gels, which allowed for more efficient transfer. 10 µl reactions contained 100 nM RNAP, 100 nM template, 2µM SutA, and 100 mM NaCl, in 1x TGA buffer, and sodium ascorbate, hydrogen peroxide, and loading buffer were added to the same final concentrations as described above.

**FeBABE DNA cleavage reactions:** Reactions were also 10 µl but contained 100 nM RNAP, 15 nM template DNA, and 2 µM SutA in 1x TGA buffer. The final NaCl concentration in these reactions (derived from the protein storage buffers) was 40 mM. The reactions were quenched by the addition of 37.5 µl 100 mM thiourea,and then 50 µl of a solution of 0.2% SDS and 2 mg/ml proteinase K was added and the reactions were incubated for 1 hr at 50 °C. 1 µl of linear acrylamide at a concentration of 10 mg/ml (as a carrier for nucleic acid precipitation), 10 µl of 3 M sodium acetate pH 5.2, and 275 µl of ethanol were added and DNA was precipitated overnight. Nucleic acid pellets were washed once with 70% ethanol, dried, and resuspended in 8 µl water. 12.5 µl primer extension reactions contained 10 mM Tris, 50 mM KCl, 1.5 mM $MgCl_2$, 5% DMSO, 2 M betaine, 250 µM dNTPs (TaKaRa, Kusatsu, Shiga Prefecture, Japan), 2.5 pmol Cy3 or Cy5 labeled primer (Integrated DNA Technologies, Coralville IA), 2.5 µl template, and 1 unit (0.2µl) Taq polymerase (NEB, Ipswich MA). After heating to 95 °C for 3 min, 15 cycles of 30 seconds at 95 °C, 30 seconds at 53 °C, and 30 seconds at 72 °C were carried out, followed by a final 3 min incubation at 72 °C. Reactions were mixed with an equal volume of formamide loading buffer (97% formamide, 10 mM Tris, 10 mM EDTA, 0.05% SDS), heated to 98 °C for 2 min, snap cooled on ice, and 8 µl were loaded onto a 12% Urea-TBE denaturing PAGE gel (Sequa-gel system, National Diagnostics, Atlanta GA) prepared with 0.5x TBE. Samples were run at 50 W (approx. 2500 V) with 0.5x TBE running buffer on a vertical sequencing gel apparatus (Ellard Instrumentation, Monroe WA). Sequencing ladders showing the positions of C or G bases in the template sequence were generated in 10 µl reactions containing 1x Thermopol reaction buffer (NEB), 1 µl Therminator polymerase (NEB), 250 µM dNTPs (TaKaRa), 25 µM ddGTP or ddCTP (TriLink Biotechnologies, San Diego CA), 100 nM template DNA (same as used in FeBABE cleavage assays), 1 µM Cy3 or Cy5 labeled primer (same as used for primer extension), and 2 M betaine. Reactions were incubated at 95 °C for 3 min, then 5 cycles of 95 °C for 30 seconds, 50 °C for 1 min, 72 °C for 1 min, followed by a final incubation at 72 °C for 3 min. Sequencing reactions were mixed with 30 µl formamide loading buffer and heated and cooled before loading as described for the samples. Sample lanes did not include loading dye, which is fluorescent in both Cy3 and Cy5 channels, but empty lanes were run with formamide loading buffer containing both Bromophenol Blue and Xylene cyanol FF.  Following electrophoresis, gels were scanned directly using the fluorescence mode of a Typhoon Trio variable mode imaging system (GE Healthcare Life Sciences), using a PMT setting of 600 and each pixel representing 200 µm. Image analysis was carried out using the FIJI analysis suite (16). Images were background subtracted and contrast-adjusted and all major bands of interest in each lane were quantified. For the FeBABE cleavage, the intensities of each band in the lanes containing

N-Fe or C-Fe SutA were normalized by dividing by the intensities of the corresponding bands in the negative control lanes containing WT SutA.

**SDS-PAGE and Western blotting**

For FeBABE initial large-format gels, markers for calibrating the observed cleavage positions were generated by cloning C-terminal fragments of β (aa 355-1357, 450-1357, 520-1357, 626-1357, and 1062-1357) into the pQE80L expression vector, and transforming into *E. coli* (see strain list). 5 ml cultures of these strains in LB were grown to late exponential phase and high levels of expression were induced by incubating with 1 mM IPTG for 4 hr. 100 µl aliquots of these cultures were pelleted by centrifugation and stored at -80 °C. Pellets were resuspended in 25 µl BugBuster (Novagen) and mixed together as follows: for 6% gels, 2 µl each of fragments 355, 450, and 520, plus 12 µl of fragment 626 were brought to a final volume of 200 µl 1x SDS loading buffer, and 10-15 µl were loaded; for 8% gel, 36 µl 1062 fragment was added to the mixture. 6% or 8% Tris-glycine-SDS gels were cast in the PROTEAN II xi Cell system using a 19:1 acrylamide: bisacrylamide mixture (Bio-Rad). Samples were denatured by heating in LDS sample buffer for 5 min at 80 °C and 1 mM DTT was added to the upper buffer to minimize protein oxidation during the 6-8 hr run time at 150 V. Following electrophoresis, gels were stained with Instant Blue colloidal Coomassie stain (Expedeon, San Diego CA) for 1 hr, briefly rinsed in water, and transferred to a nitrocellulose membrane using 1x Towbin transfer buffer containing 20% methanol and 0.03% SDS, for 4-6 hr at 250 mA using a Hoefer TE62 transfer apparatus (Hoefer, Holliston MA). Membranes were blocked for 1 hr in 2.5% non-fat dry milk in TBST, then incubated in primary antibody (EPR18704, Abcam, Cambridge MA) at a 1:1500 dilution for 8 hr, washed in TBST and incubated in the secondary antibody (goat anti-rabbit HRP, Sigma, St. Louis) at a dilution of 1:5000 for 1 hr before washing in TBST and developing with SuperSignal™ West Pico PLUS Chemiluminescent Substrate (ThermoFisher, Waltham MA) according to instructions. Blots were exposed to x-ray film for 5-15 min.

For the FeBABE reactions to analyze the effects of different σ factors and DNA templates, samples were run on 4-20% gradient Tris-glycine SDS mini-gels (Bio-Rad) for 1 hr. at 150 V, then stained with Coomassie Colloidal Blue and transferred to pre-cut nitrocellulose membranes (Bio-Rad) for 8 hr at 20 V in 1X Towbin transfer buffer without methanol or SDS added. The membranes were cut to separate region containing the uncleaved β subunit band from the region containing the cleavage products, which were of much lower abundance. Western blotting for the membrane region containing the cleavage products was the same as described above for the large-format gel, but the region containing the uncleaved band was incubated with primary antibody diluted 1:2000 and secondary antibody diluted 1:20,000. The two regions of the membrane were then placed next to each other for exposure to X-ray film. The cutting of the membrane occasionally resulted in the appearance of a second band immediately below the uncleaved band (especially in the outer lanes of the gel), which was just the edge of the uncleaved band. For analysis and Western blotting of BPA cross-linking in various holoenzyme/DNA contexts, reaction volumes were 10 µl and contained 100 nM RNAP core or holoenzyme, the concentrations of the BPA54 variant listed in the figures, 100 nM template DNA, 100 mM NaCl, and TGA buffer. Cross-linking was carried out as described above for LC-MS/MS analysis, and then samples were added to LDS loading buffer. 3-8% Tris-acetate gels and Tris-acetate-SDS running buffer (NuPAGE) were used to maximize separation of the cross-linked β+SutA band from the uncross-linked β only band. Subsequent steps of the Western blotting protocol were the same as for the FeBABE mini-gels, using the same antibody dilutions as for the uncleaved portion, described above.

**Data visualization**

Unless otherwise noted, molecular structures were visualized using the Chimera suite (14). Graphs were produced using the ggplot2 library in R (23). Gel images were background-subtracted and contrast adjusted using the FIJI suite (16). NMR spectra were visualized using the CcpNmr Analysis suite (7). LC-MS/MS spectra for cross-linked peptides were shown using StavroX software (21). Figures were assembled using Adobe Acrobat CC2018.

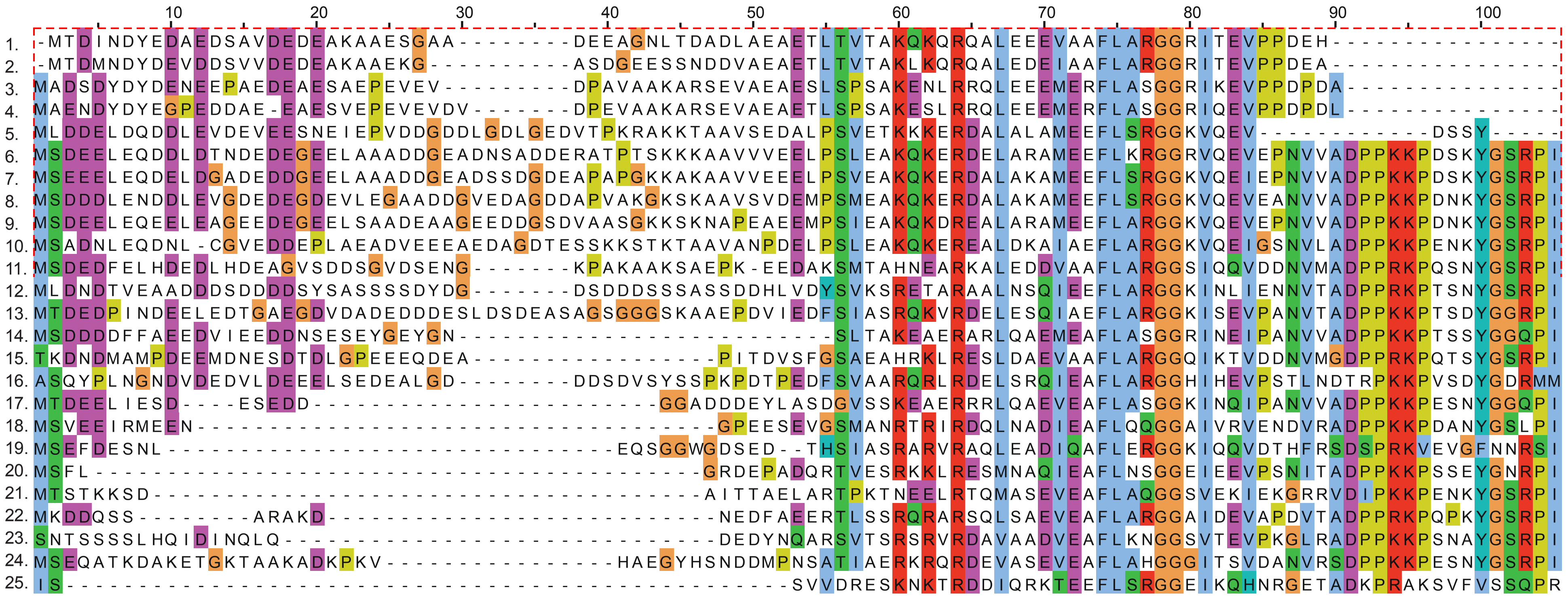

1. *Acinetobacter baumannii*
2. *Acinetobacter equi*
3. *Perlucidibaca piscinae*
4. *Perlucidibaca aquatica*
5. *Oblitimonas alkaliphila*
6. *Pseudomonas stutzeri*
7. *Pseudomonas aeruginosaUCBPP-PA14*
8. *Pseudomonas putida*
9. *Azotobacter vinelandiiDJ*
10. *Ventosimonas gracilis*
11. *Oceanobacter kriegii*
12. *Simiduia agarivorans*
13. *Cellvibrio japonicus*
14. *Saccharophagus degradans*
15. *Thalassolituus oleivorans*
16. *Marinimicrobium agarilyticum*
17. *Teredinibacter turnerae*
18. *Mangrovitalea sediminis*
19. *Marinobacter lutaoensis*
20. *Endozoicomonas numazuensis*
21. *Reinekea blandensis*
22. *Microbulbifer agarilyticus*
23. *Gynuella sunshinyii*
24. *Oleispira antarctica*
25. *Gamma proteobacterium HTCC2207*

**Figure S1.** P. aeruginosa UBCPP-PA14-centric alignment of representative SutA homologs. SutA homologs were detected by BLAST, and representatives were selected from multiple families in each of the four orders in which SutA could be found (Alteromonadales, Cellvibrionales, Oceanospirillales, and Pseudomonadales). After alignment using the MEGA6 software suite (12), gaps in the P. aeruginosa UBCPP-PA14 sequence were removed and the aligment was visualized using the Jalview2 applet(13).

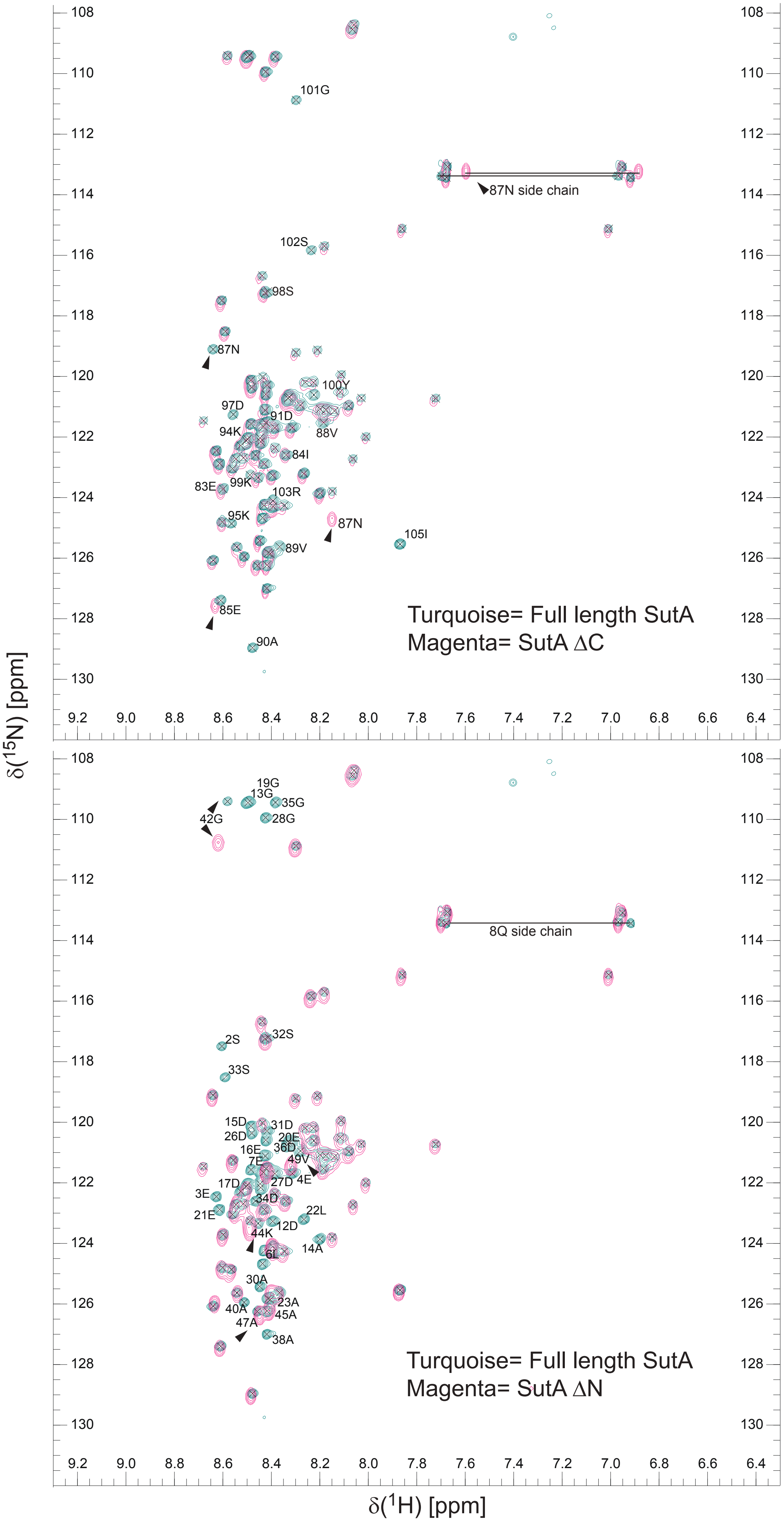


**Figure S2.** 15N HSQC spectra comparing the full-length WT SutA to ΔN and ΔC proteins. 15N HSQC spectra for SutAΔC (top) and SutAΔN (bottom) (both in magenta) were overlaid on the 15N HSQC for the full-length SutA (turquoise). Apart from the loss of the truncated residues, only a few peaks near the newly created C- or N-terminus are perturbed.

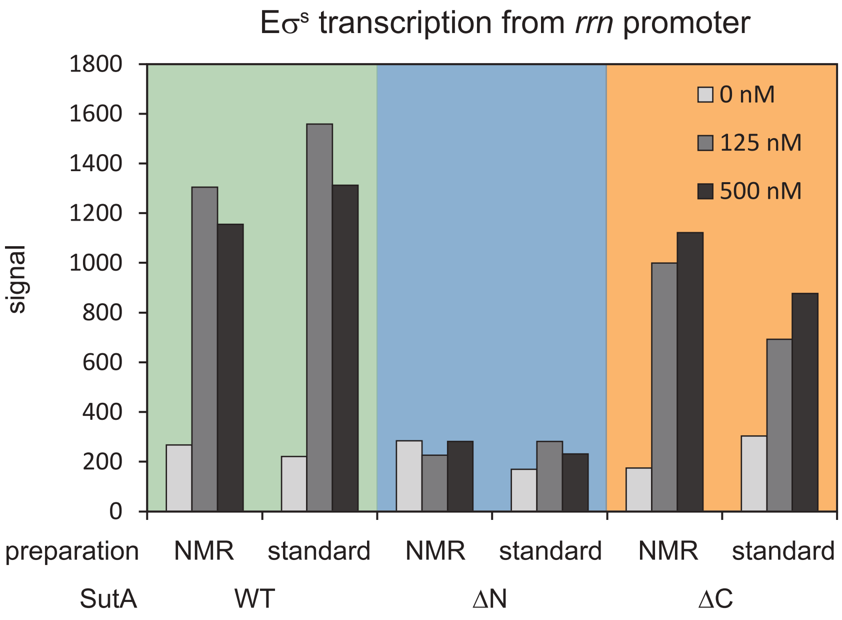


**Figure S3.** In vitro transcriptional activity of SutA proteins prepared for NMR, compared to the same proteins prepared using standard methods. Activity of proteins produced for NMR was tested using the single-turnover initiation assay with E$\sigma^S$ as described for Figure 3D.

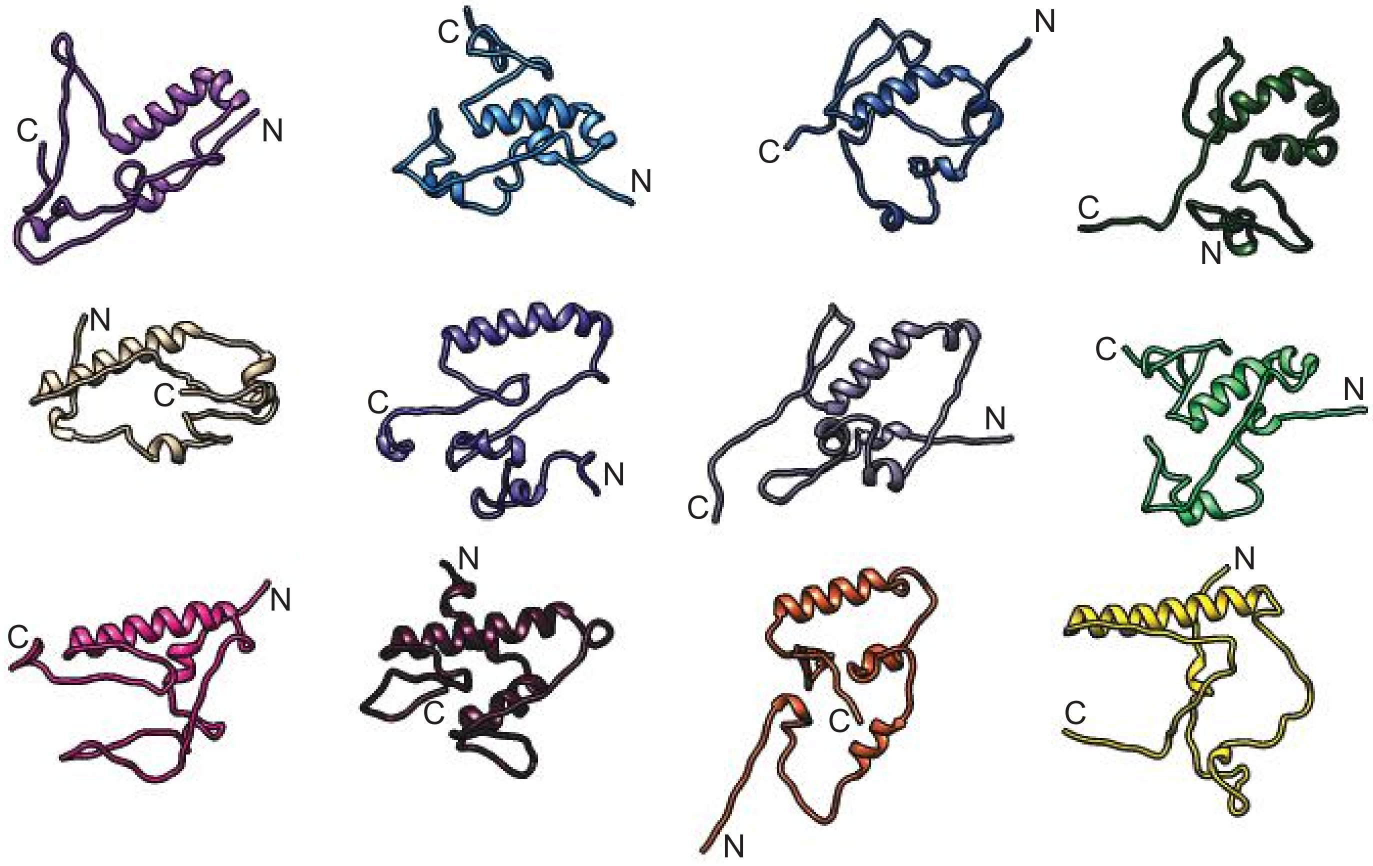

**Figure S4.** A selection of low-resolution SutA decoys generated by PyRosetta modeling utilizing NMR chemical shift and RDC data. The model used for Figure 1D is in the second row, first column. SutA is a very flexible protein, with its only secondary structural feature being an α-helix encompassing residues 56-76, and even that helix displays some predicted possible flexibility. We did not detect a peak for the Gln61 residue, the point in the helix that shows the most variation in these models.

BS$^3$ crosslinking:

Evaluation of crosslinks among RNAP subunits:

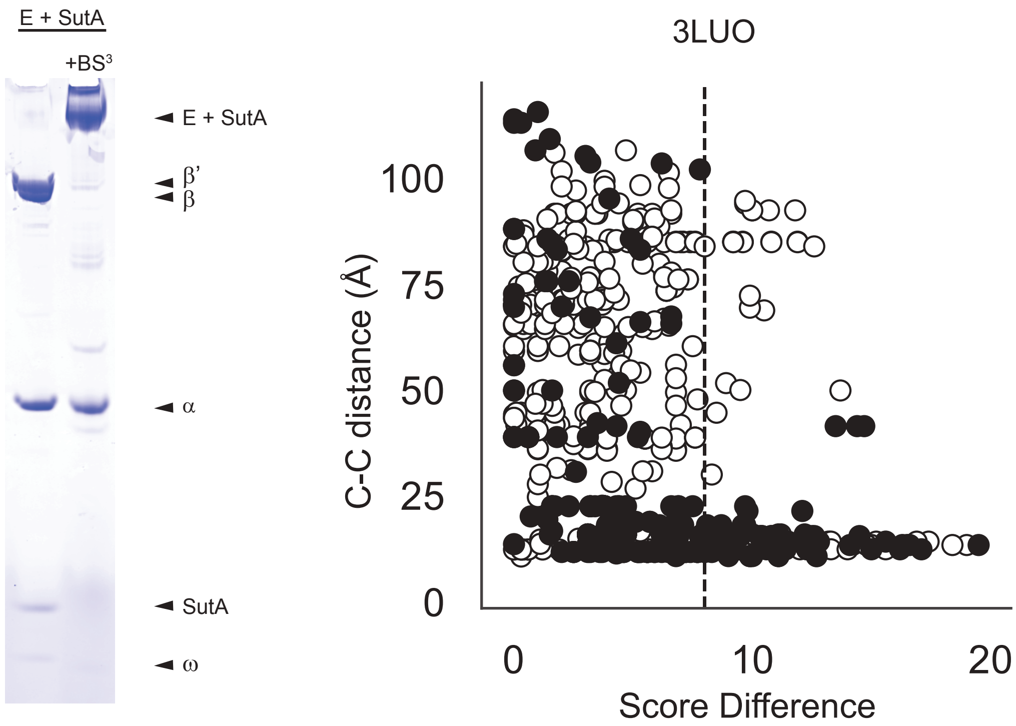


| crosslink | # spectra | score diff. |
|---|---|---|
| SutA 62-β 116 | 15 | 15.6 |
| SutA 69-β 116 | 11 | 18.8 |
| SutA 62-β 481 | 1 | 8.2 |
| SutA 69-β 45 | 1 | 11.4 |

**Figure S5.** BS$^3$ crosslinking visualization and analysis. A Coomassie-stained SDS-PAGE gel showing BS$^3$ cross-linking of RNAP-SutA complexes, a comparison of the score differences calculated for intra-RNAP cross-links versus the distances between the cross-links in a published RNAP structure that was used to determine an appropriate score difference cut-off for likely real cross-links, and a list of SutA-RNAP cross-links, the number of spectra in which they were detected, and the maximum score difference observed.

BPA crosslinking:

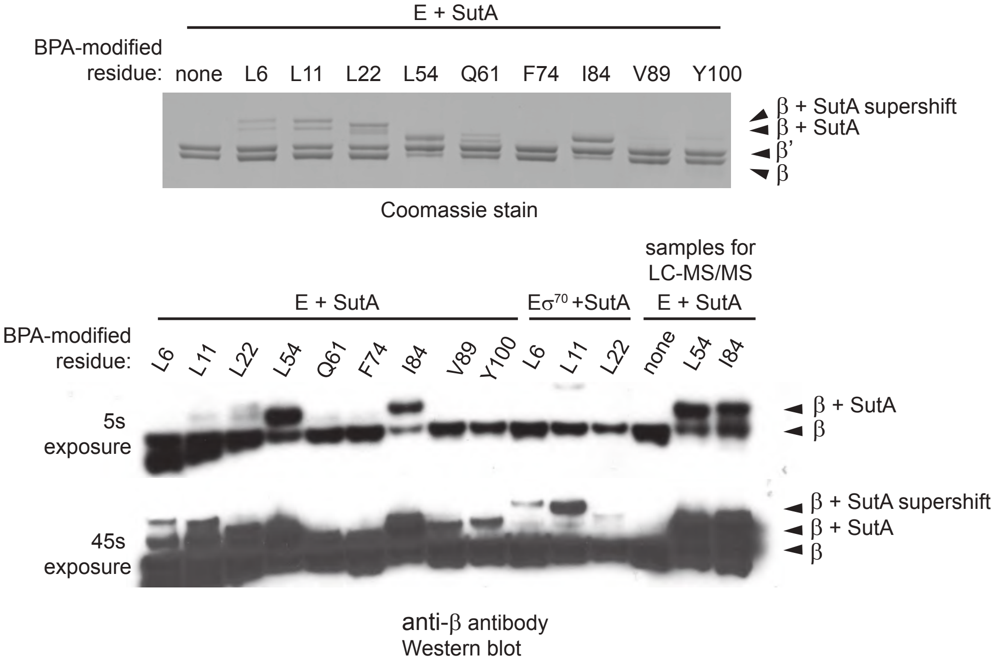


**Figure S6.** A Coomassie stained SDS-PAGE gel and a Western blot using an anti-β antibody showing the formation of shifted β bands following SutA BPA variant crosslinking to β.

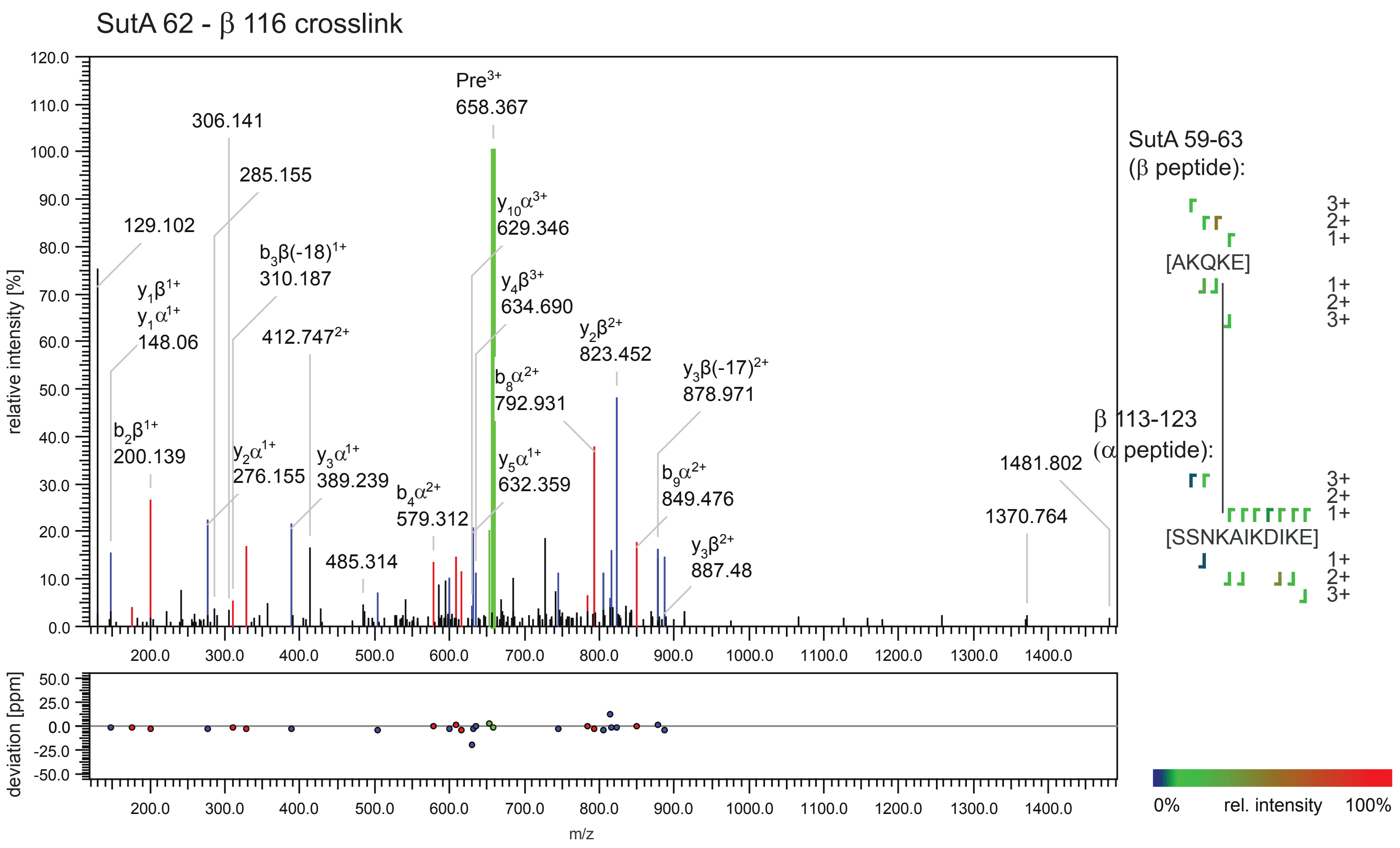


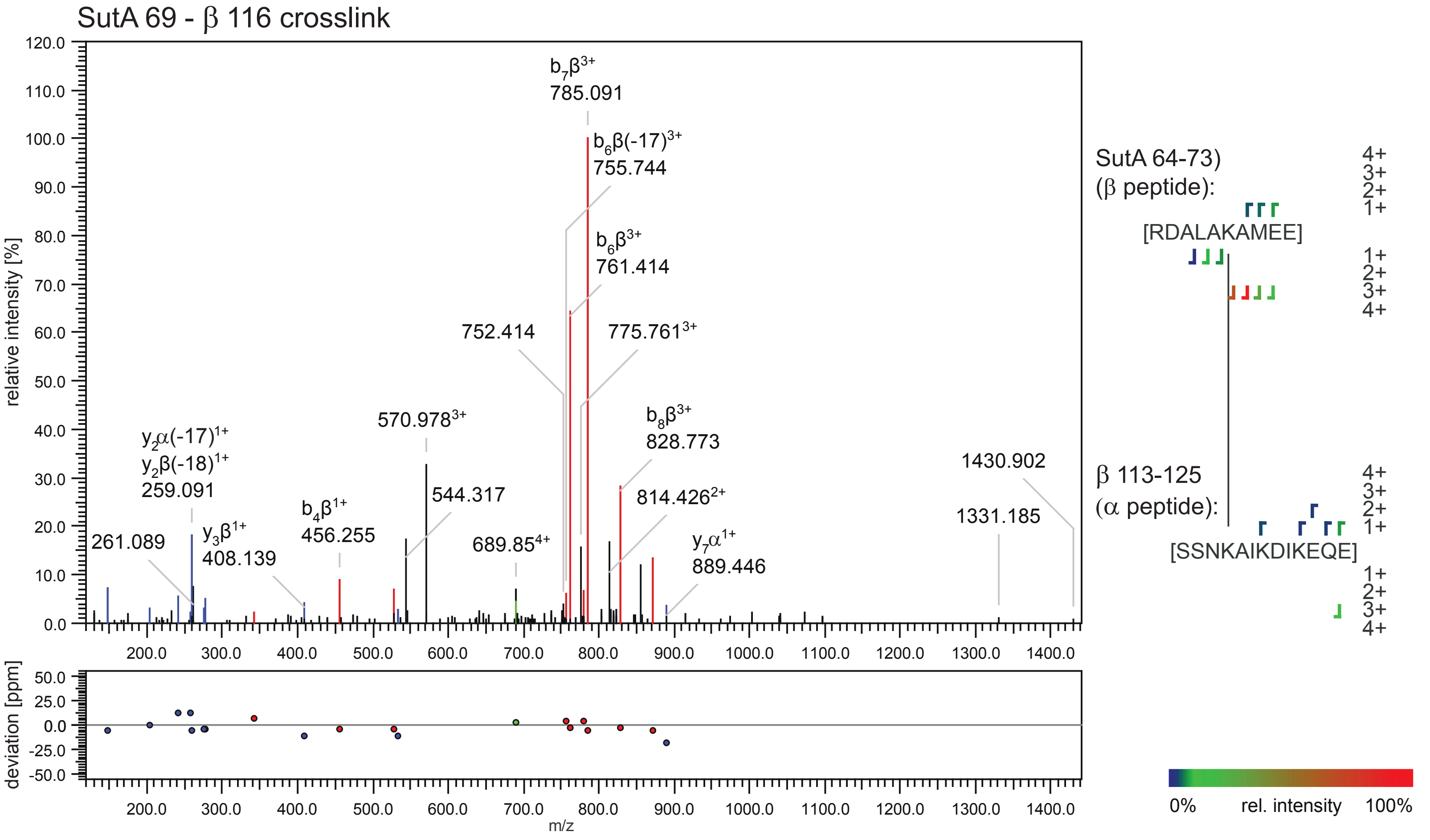


**Figure S7.** LC-MS/MS spectra from cross-linked peptides detected in the BS3 experiment. Output from StavroX analysis software shows multiple detected fragment ions from both component peptides, indicating high-quality identifications of cross-linked peptides (continues on next page).

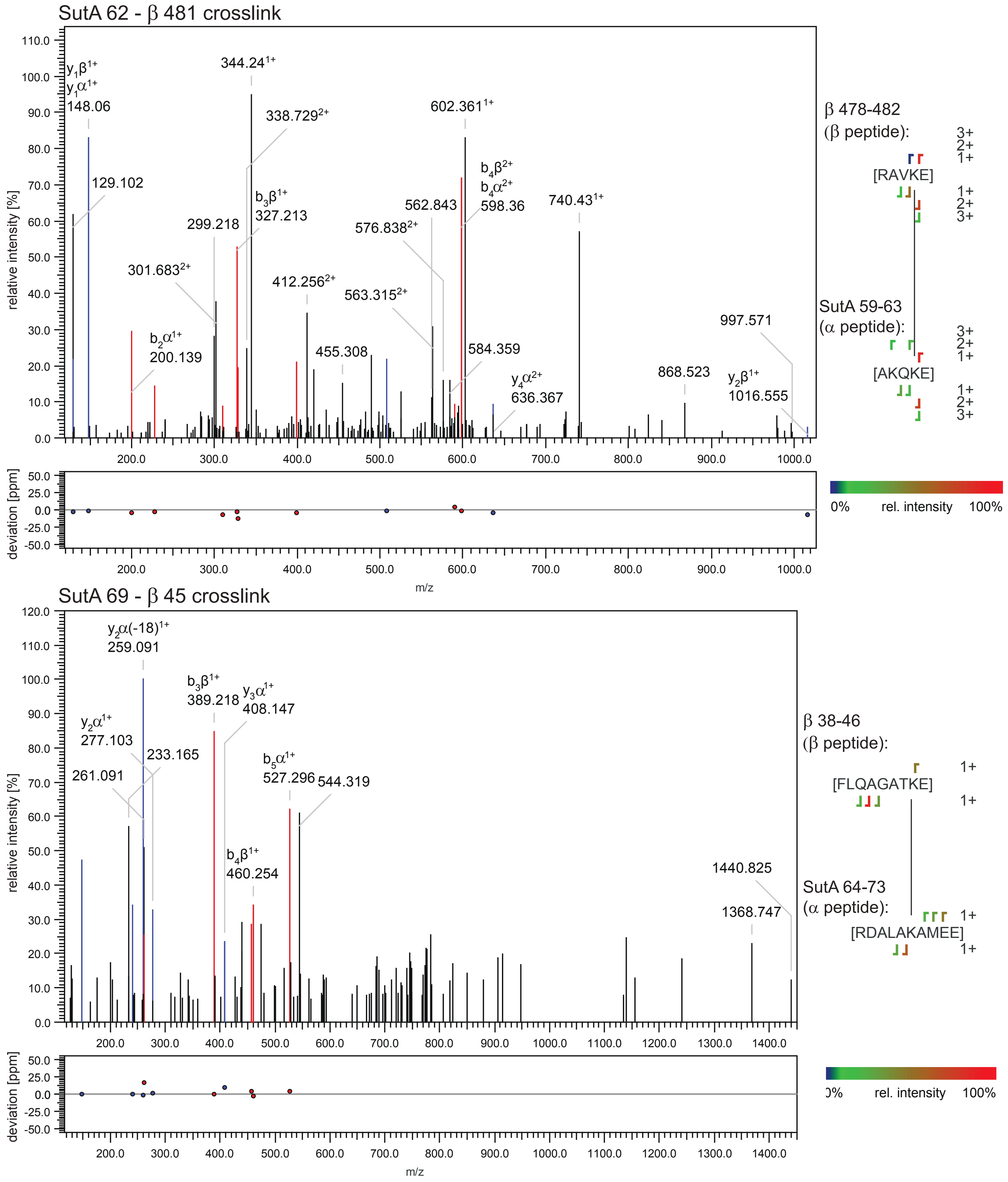


**Figure S7** (continued)

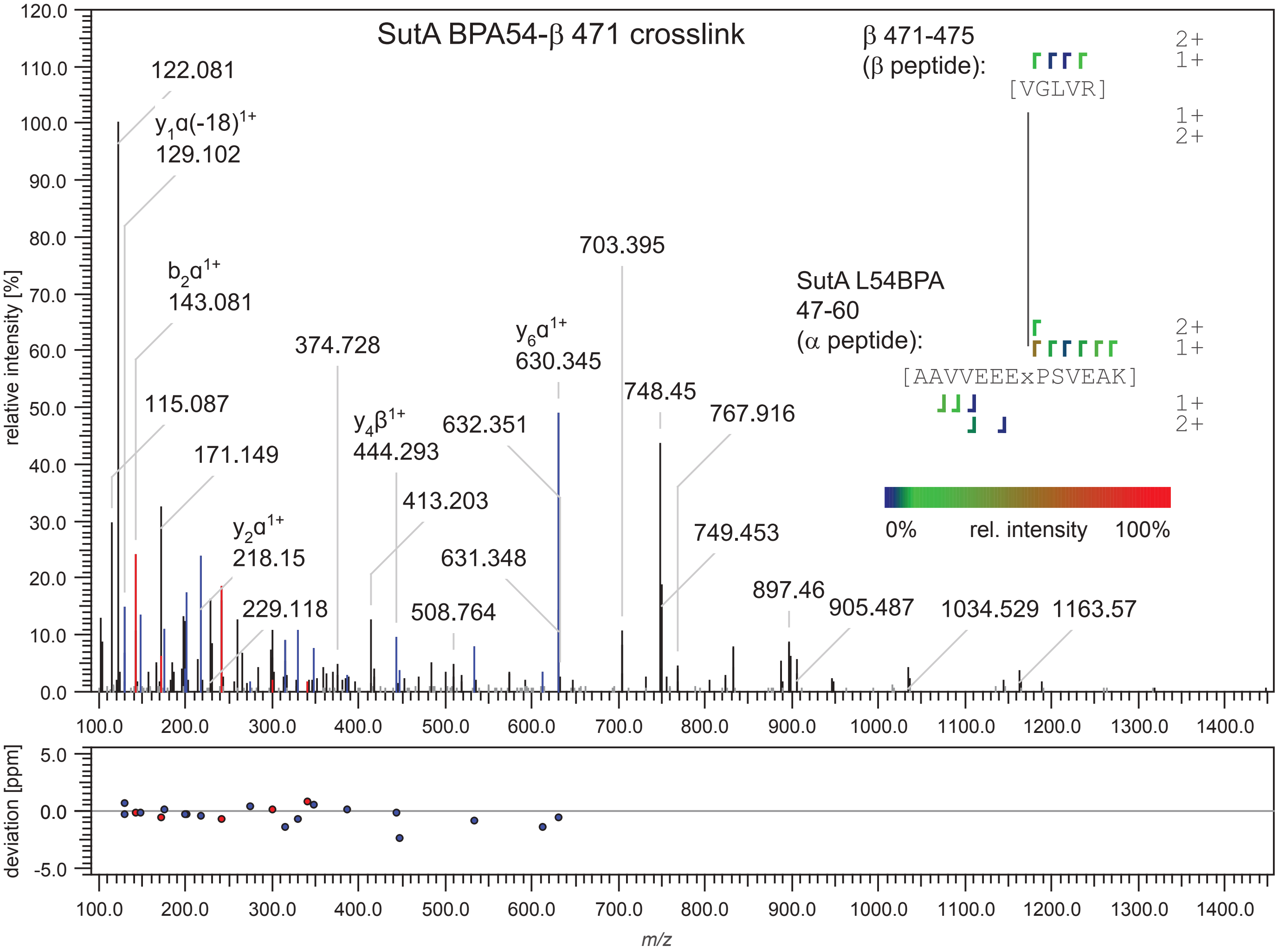


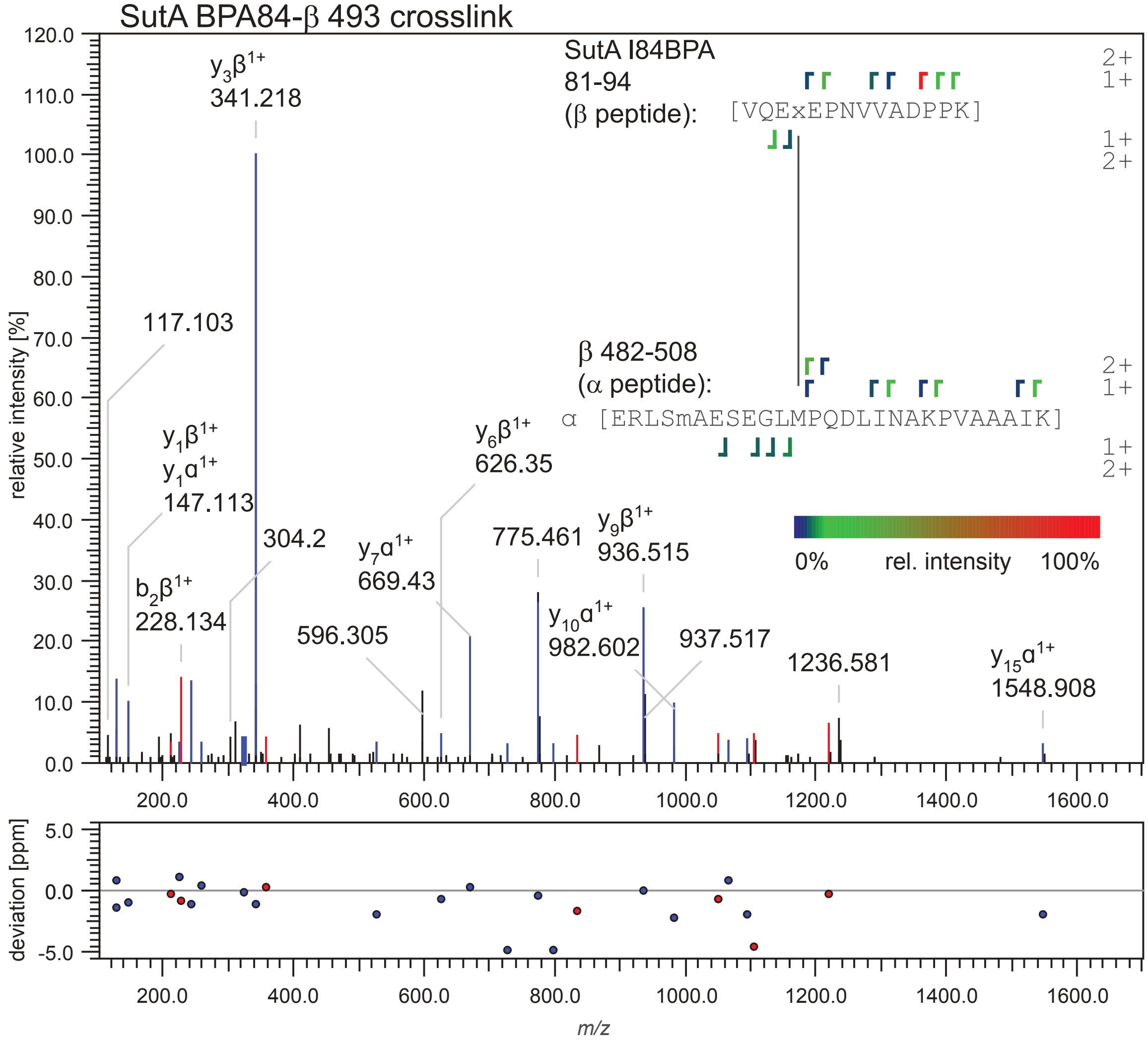


**Figure S8.** LC-MS/MS spectra from cross-linked peptides detected in the BPA experiments. Output from StavroX analysis software shows multiple detected fragment ions from both component peptides, indicating high-quality identifications of cross-linked peptides (continues on next page).

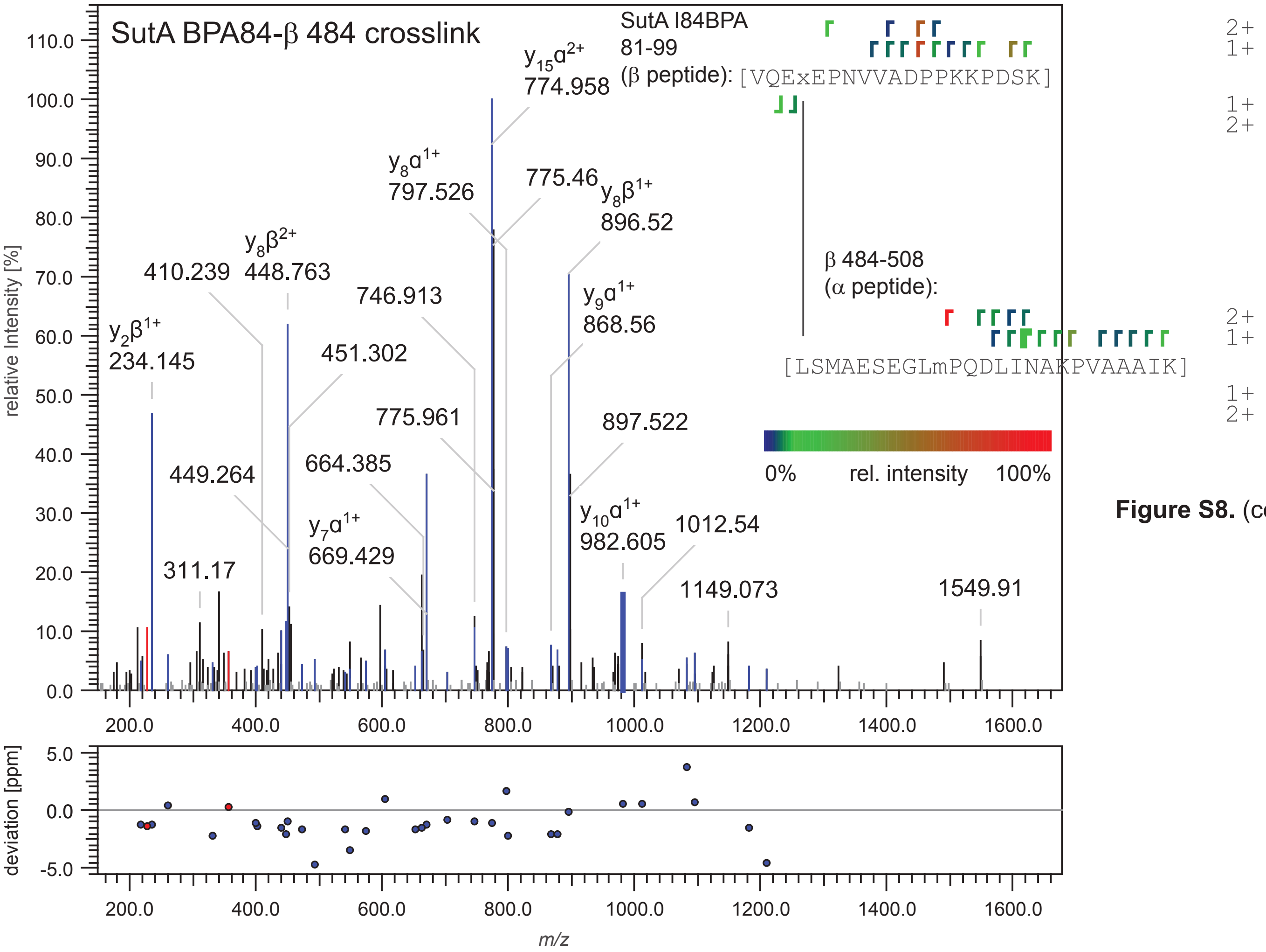


**Figure S8.** (continued)

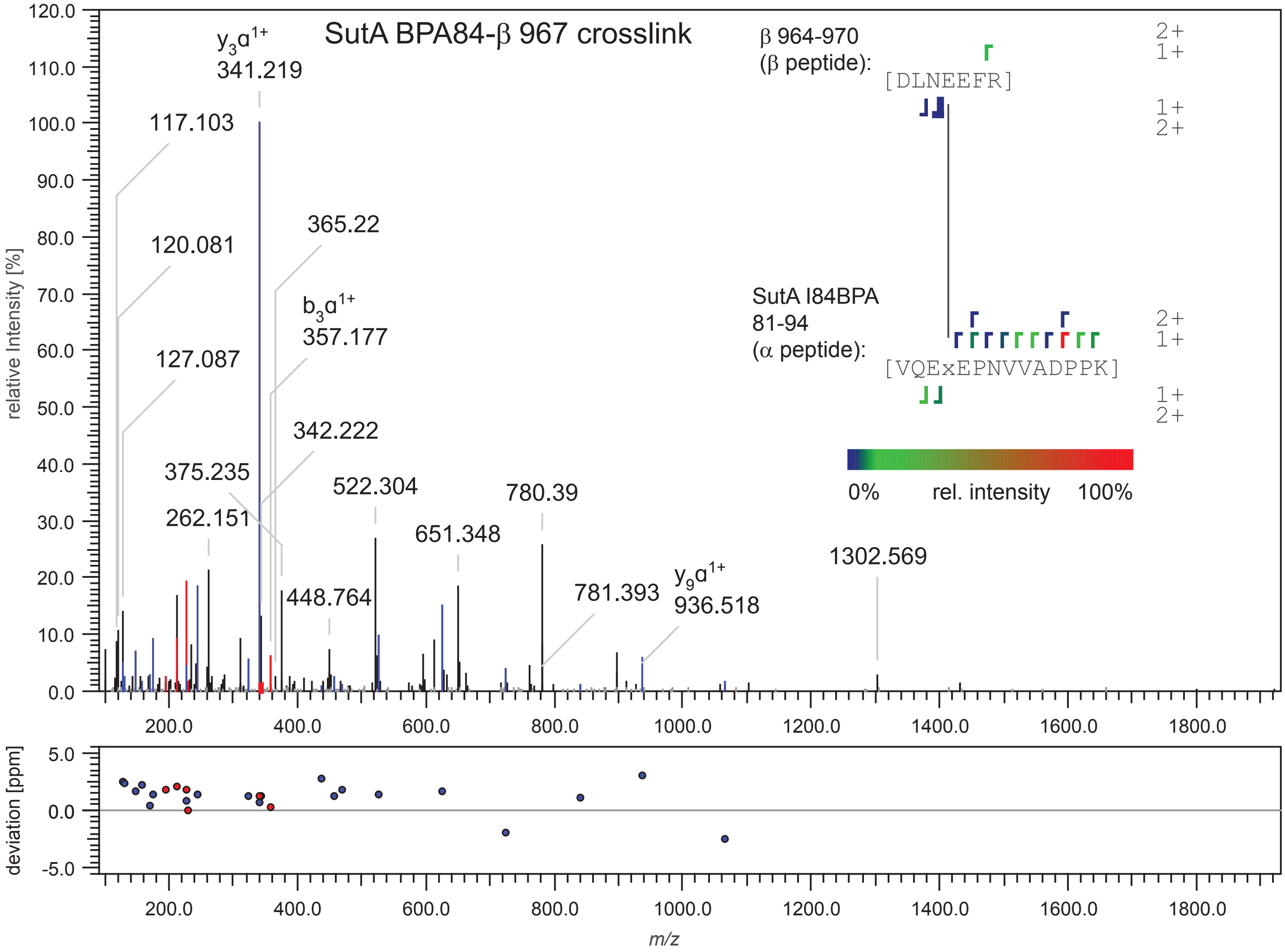

FeBABE cleavage:

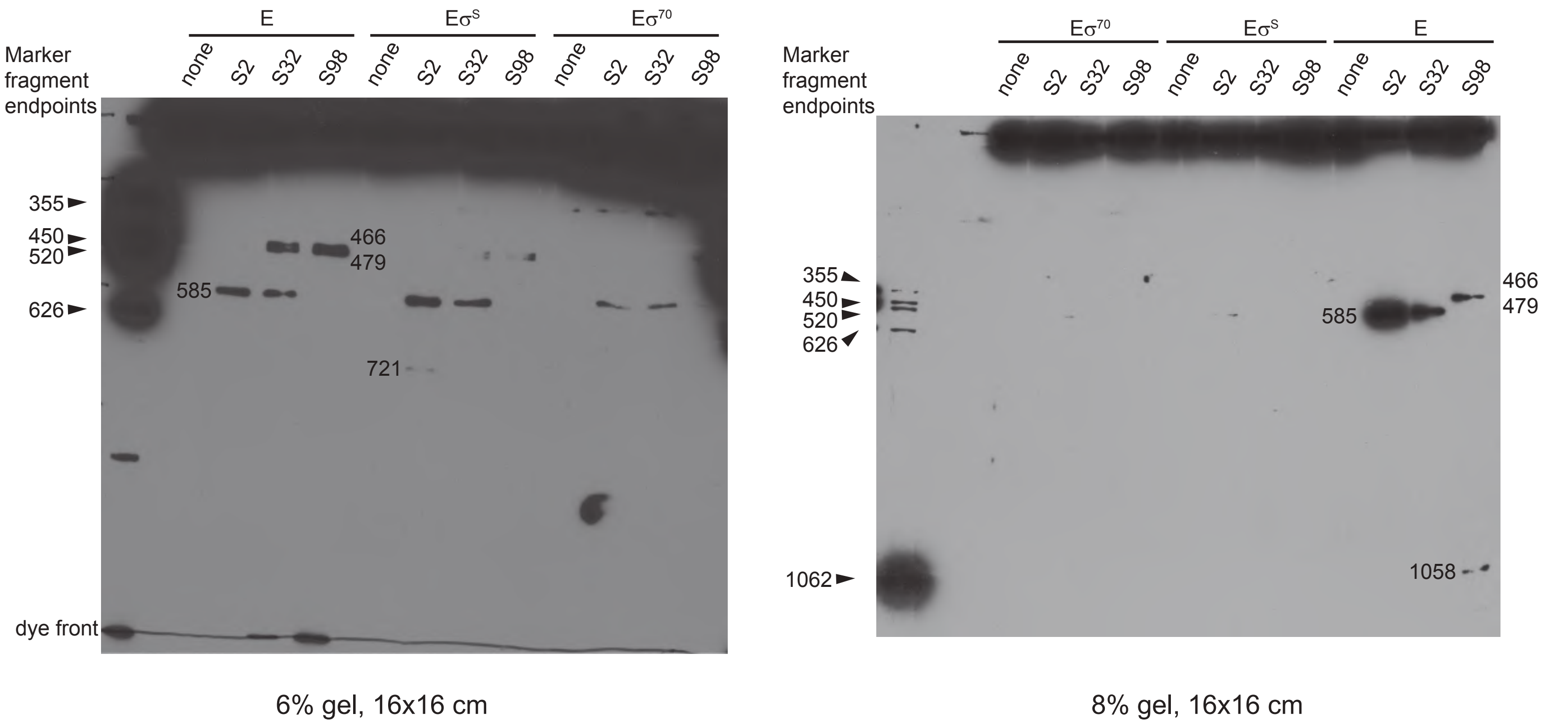


**Figure S9.** 16x16 cm Western blots from gels run with two different percentages of acrylamide, for calculation of FeBABE cleavage positions. Western blotting of FeBABE cleavage products, using a monoclonal antibody raised against a peptide C-terminal to amino acid 1300 of the E. coli b sequence (Abcam EPR18704), was performed in a large format to allow for accurate calculation of the molecular weights of the cleavage products. Known C-terminal fragments, generated by overexpressing a cloned fragment of *P. aeruginosa* β in *E. coli*, were run as markers (numbers along the side of the blot indicate the N-terminal endpoint of the fragment, not its molecular weight). The molecular weights of these marker fragments were calculated using the ExPASy Compute pI/Mw tool, and the log of this value was plotted against the ratio: (distance traveled by band/distance traveled by dye front). The linear relationship established was used to calculate the molecular weights of the FeBABE cleavage products based on their band/dye front ratios, and those molecular weights were used to determine the amino acid position at which the cleavage occurred, which is indicated on the blot next to the band.

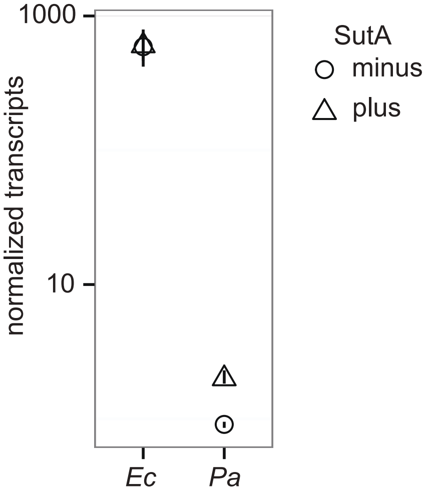


**Figure S10.** Induction of SutA overexpression in E. coli does not cause upregulation of rrn expression. SutA under control of an arabinose-inducible promoter on the pMQ72 plasmid backbone, or the empty vector, was introduced into either E. coli MG1655 or P. aeruginosa UBCPP-PA14 Δ sutA, and cells were grown into late stationary phase in LB in the presence of 20 mM arabinose before harvesting them, extracting RNA, and measuring nascent rrn transcript levels by qRT-PCR. Symbols represent the average value from 3 biological replicates, and vertical lines represent the range of values observed.

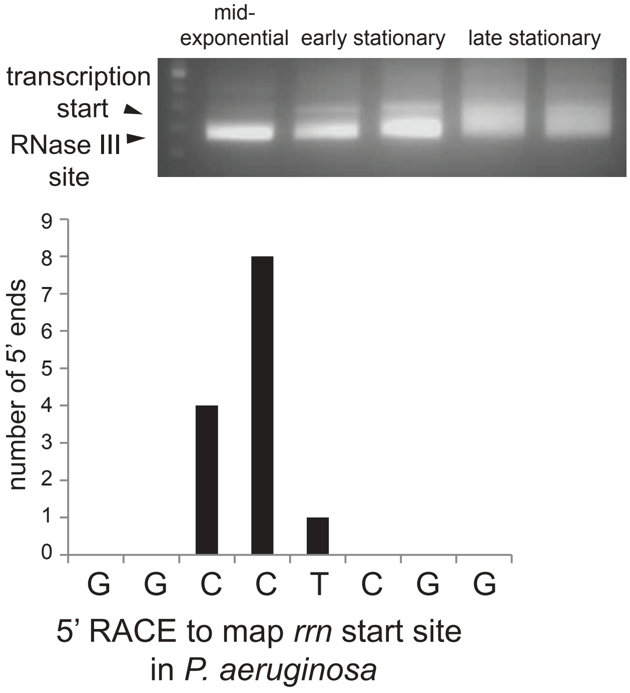


**Figure S11.** 5'RACE to determine the transcription start site for rrn in P. aeruginosa. Total RNA was extracted from P. aeruginosa UCBPP-PA14 in exponential, early stationary, or late stationary phase, and the leader sequence of the rrn transcript was reverse transcribed, T-tailed, PCR-amplified, and cloned into pUC18. Several clones from the stationary phase time points were from transcripts whose 5' ends corresponded to the RNase III cleavage site in the rrn leader, based on comparison to the E. coli sequence, but clones whose 5' ends corresponded to putative transcription start sites were distributed as shown.

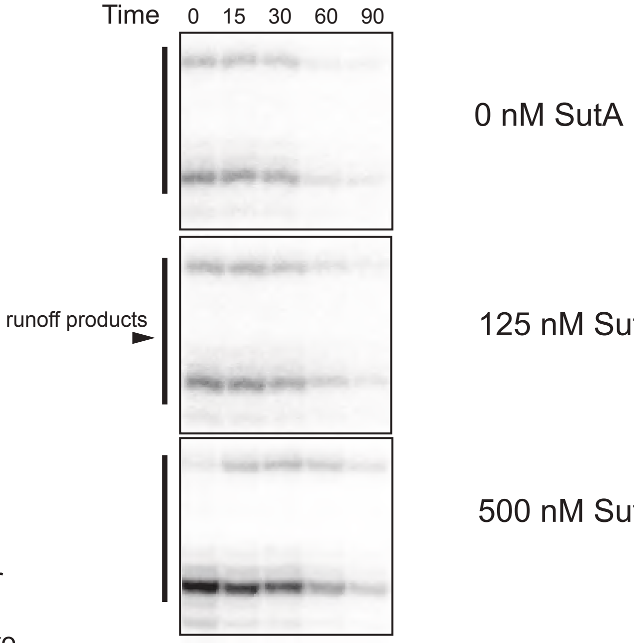


**Figure S12.** Relevant regions of gels showing example reactions in open complex stability assays. See also Figure S15 for full length gel example, and main text and Extended Materials and Methods for details.

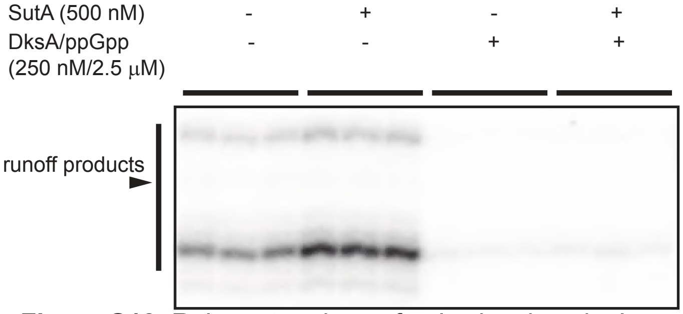


**Figure S13.** Relevant regions of gels showing single turnover initiation assays with or without SutA and DksA/(p)ppGpp. See also Figure S15 for full length gel example and main text and Extended Materials and Methods for details..

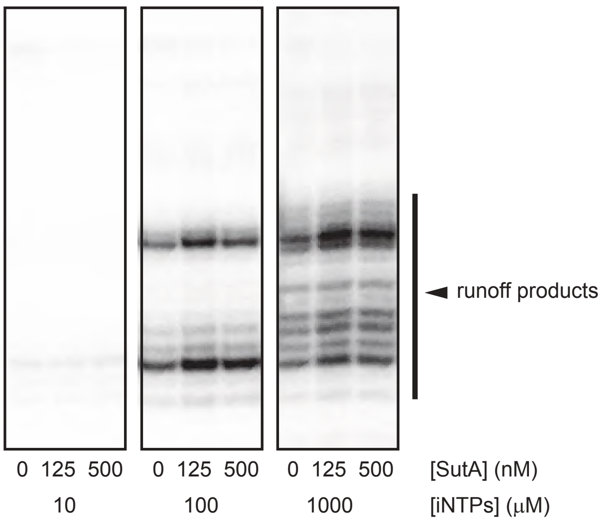


**Figure S14.** Relevant regions of gels showing example reactions in single turnover initiation assays at different [iNTPs]. See also Figure S15 for full length gel example and main text and Extended Materials and Methods for details..

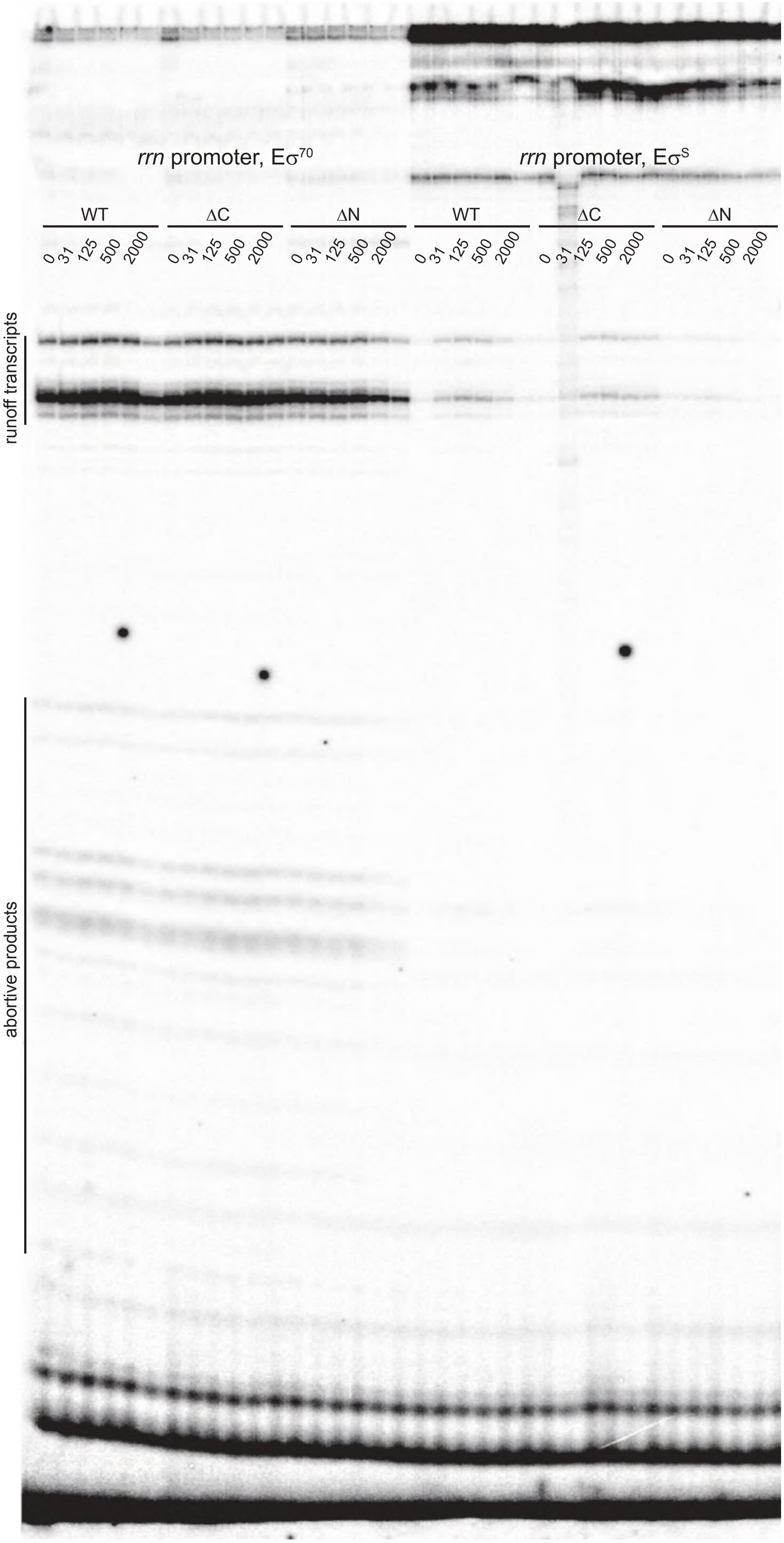


**Figure S15.** Full length gels for one replicate of each holoenzyme/SutA combination shown in Figure 3D. The rrn/$E\sigma^{S}$/31 nM SutA ΔC sample was degraded, and an additional replicate of this sample was run on a subsequent gel.

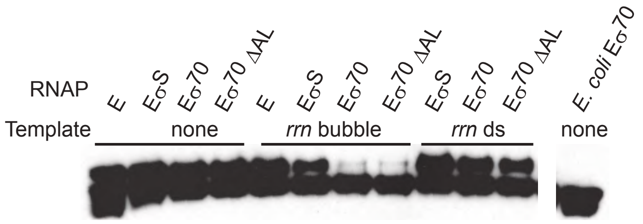


**Figure S16.** Longer exposure of Western blot showing cross-linking of 2 μM L54BPA SutA to β. A low level of cross-linking is detectable in the presence of $E\sigma^{70}$ and the rrn bubble template, but no cross-linking is detected in the presence of E. coli $E\sigma^{70}$, even in the absence of DNA.

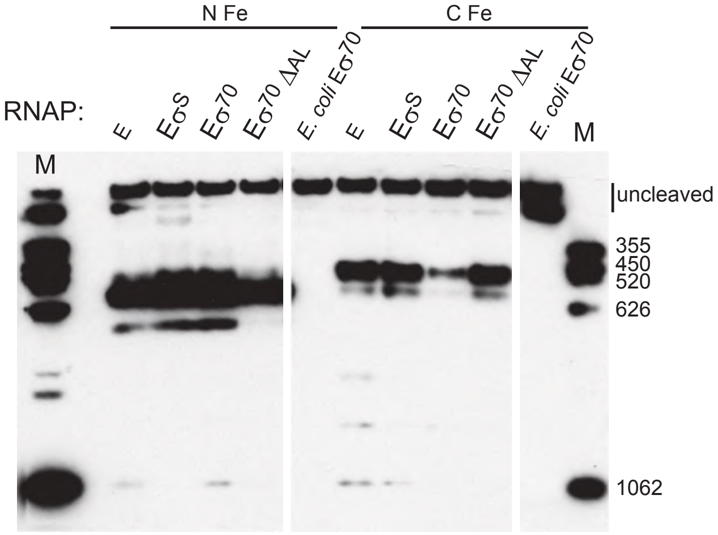


**Figure S17.** Western blot showing FeBABE cleavage experimental controls. β fragment standards used to determine cleavage positions were run on the mini-gel format for direct comparison to cleavage products observed in open complex contexts, and E. coli $E\sigma^{70}$ FeBABE cleavage experiments were also run and showed no detectable cleavage.

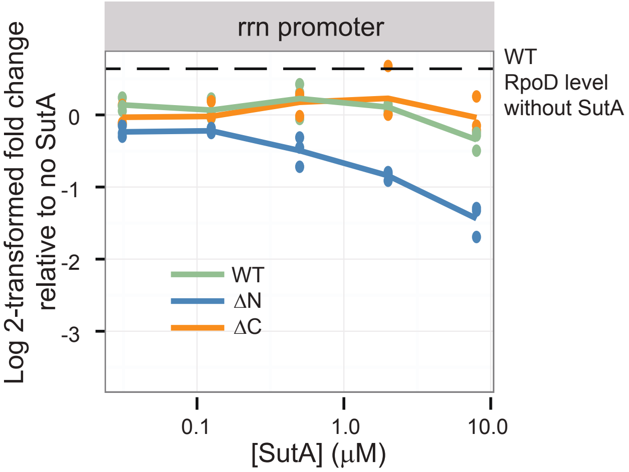


**Figure S18.** In vitro transcription experiments using $E\sigma^{70}\Delta$ AL, with the transcription level of the $E\sigma^{70}$ holoenzyme in the absence of SutA shown for comparison. Single-turnover initiation assays were performed as described in Figure 3D. $E\sigma^{70}\Delta$AL appears to have a mild transcription initiation defect, and causes SutA to have more muted effects on initiation.

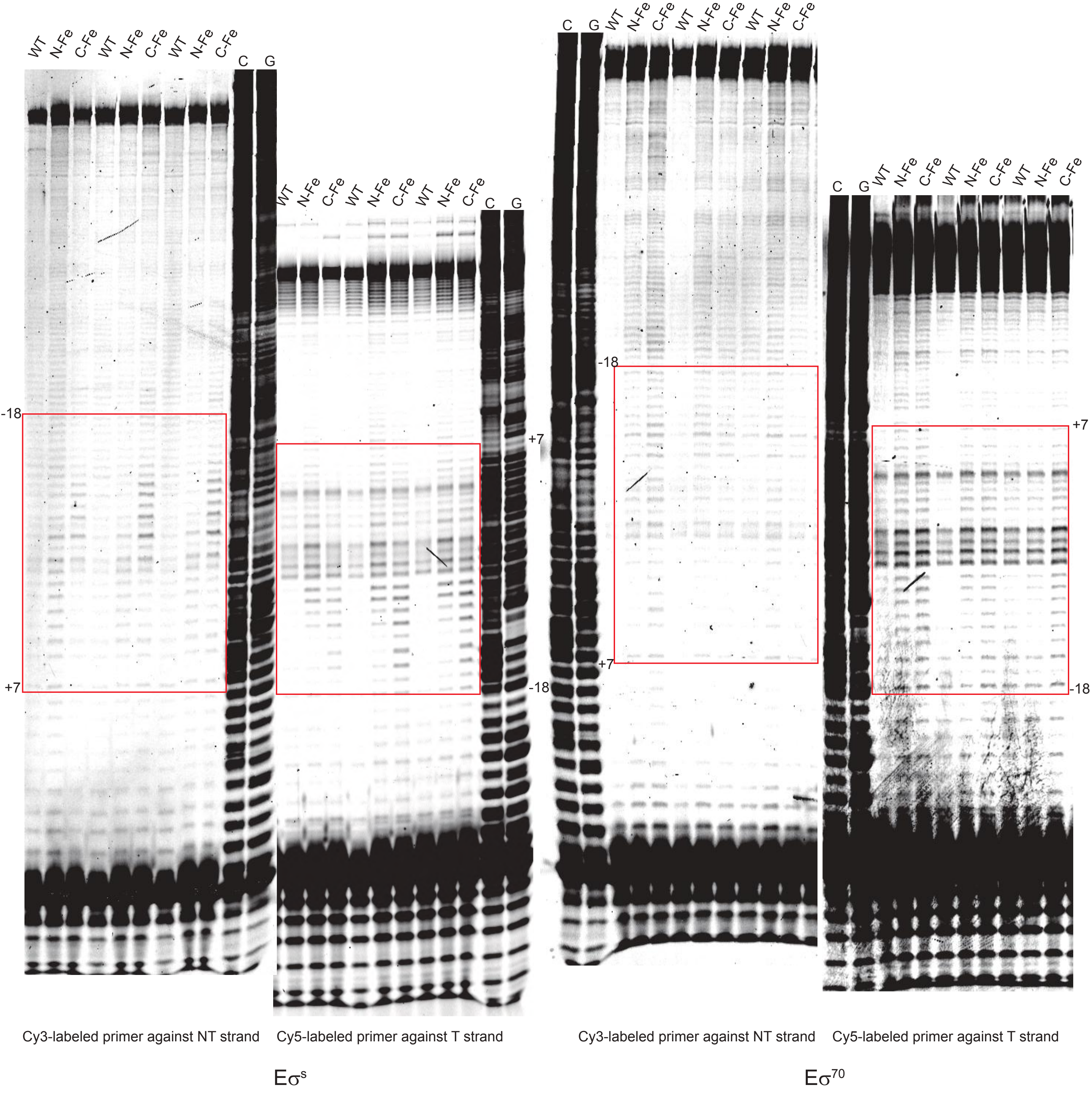


**Figure S19.** Full length gels for triplicate measurements of FeBABE DNA cleavage for rrn promoter with $E\sigma^S$ and $E\sigma^{70}$. Regions represented by color scales in Figure 5C are outlined with red boxes. Gel images are contrast-adjusted to make weak cleavage more visible. Band intensity was quantified with contrast settings that avoided saturation of any bands, and band identities were determined at lower contrast where sequencing ladders were readable.

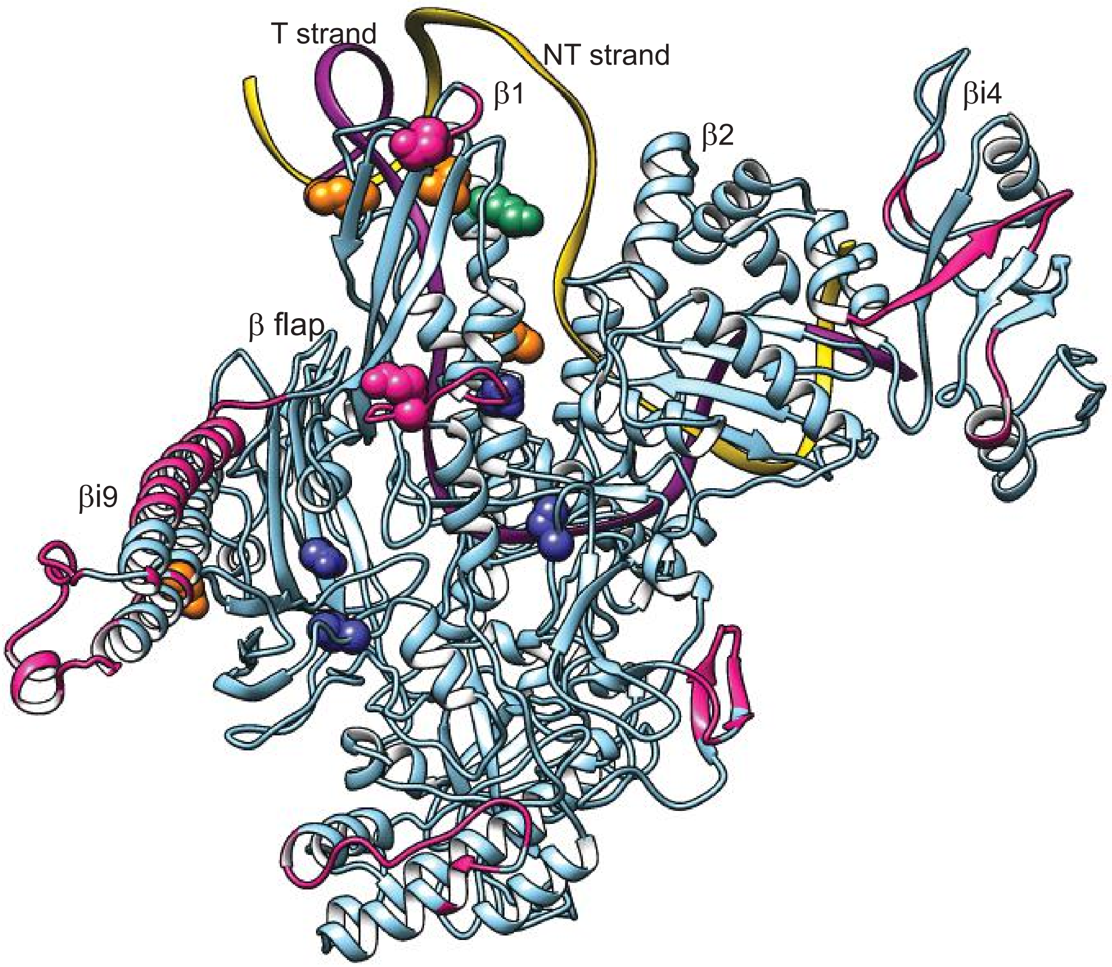


**Figure S20.** A comparison of regions of reduced similarity between *E. coli* and *P. aeruginosa* β, and SutA cross-link or cleavage sites. Because SutA does not influence *rrn* transcription in *E. coli* in vivo and does not show cross-linking or cleavage interactions with *E. coli* $E\sigma^{70}$ in vitro, it stands to reason that the functionally relevant binding site for SutA on β should involve regions that are different between *E. coli* and *P. aeruginosa*. Regions in which a sliding window of 8 aa was less than 50% similar between the two species are colored magenta. Residues involved in SutA cross-links or cleavages are represented as spheres. The color coding from figure 2 is retained ($BS^3$ cross-links green, BPA cross-links orange, FeBABE cleavages blue) except where cross-links coincide with regions of reduced similarity, which are colored magenta. Such regions of lower similarity are a small percentage of the overall sequence, and mostly occur in large species-specific insertions or surface loops. Two of the $BS^3$ crosslinks occur in small surface loops of the β1 domain that are different between *E. coli* and *P. aeruginosa*.

Table 1: SutA Backbone chemical shift values (ppm)

| Residue | number | C | CA | CB | HA | N | HN |
|---|---|---|---|---|---|---|---|
| Met | 1 | 176.33 | 55.92 | 32.94 | ND | ND | ND |
| Ser | 2 | 174.77 | 58.50 | 63.86 | 4.48 | 117.49 | 8.60 |
| Glu | 3 | 176.79 | 57.36 | 29.98 | 4.22 | 122.48 | 8.63 |
| Glu | 4 | ND | 56.79 | 30.12 | 4.25 | 121.69 | 8.31 |
| Glu | 5 | 176.49 | ND | ND | ND | ND | ND |
| Leu | 6 | ND | 55.06 | 42.34 | ND | 124.68 | 8.44 |
| Glu | 7 | 176.49 | 56.41 | 30.00 | ND | 121.66 | 8.43 |
| Gln | 8 | ND | ND | ND | ND | ND | ND |
| Asp | 9 | ND | ND | ND | ND | ND | ND |
| Glu | 10 | ND | ND | ND | ND | ND | ND |
| Leu | 11 | 176.65 | 56.62 | 30.05 | ND | ND | ND |
| Asp | 12 | 177.03 | 55.19 | 42.18 | 4.56 | 123.29 | 8.40 |
| Gly | 13 | 174.06 | 45.35 | ND | 3.86 | 109.45 | 8.50 |
| Ala | 14 | 177.61 | 52.45 | 19.44 | 4.80 | 123.87 | 8.20 |
| Asp | 15 | 176.36 | 54.28 | 41.03 | 4.56 | 120.20 | 8.48 |
| Glu | 16 | 175.75 | 55.69 | 29.67 | 4.24 | 121.24 | 8.43 |
| Asp | 17 | ND | 54.63 | 41.24 | 4.59 | 122.29 | 8.53 |
| Asp | 18 | ND | ND | ND | ND | 122.29 | 8.53 |
| Gly | 19 | ND | 45.38 | ND | ND | 109.38 | 8.49 |
| Glu | 20 | ND | 56.39 | 30.29 | 4.24 | 120.70 | 8.33 |
| Glu | 21 | ND | ND | ND | 4.24 | 122.88 | 8.61 |
| Leu | 22 | 177.18 | 54.97 | 42.36 | 4.18 | 123.21 | 8.27 |
| Ala | 23 | ND | 52.25 | 19.19 | 4.25 | 125.88 | 8.40 |
| Ala | 24 | ND | ND | ND | ND | ND | ND |
| Ala | 25 | ND | 52.45 | 19.28 | ND | ND | ND |
| Asp | 26 | ND | 54.05 | 41.16 | 4.59 | 120.28 | 8.48 |
| Asp | 27 | 176.85 | 54.37 | 41.09 | 4.56 | 121.69 | 8.39 |
| Gly | 28 | ND | 45.39 | ND | ND | 109.95 | 8.42 |
| Glu | 29 | 176.53 | 56.26 | 30.16 | ND | ND | ND |
| Ala | 30 | 177.54 | 52.43 | 19.25 | 4.25 | 125.43 | 8.45 |
| Asp | 31 | 176.47 | 54.10 | 41.24 | 4.56 | 120.31 | 8.42 |
| Ser | 32 | 174.86 | 58.27 | 63.82 | 4.43 | 117.23 | 8.42 |
| Ser | 33 | 174.51 | 58.59 | 63.87 | 4.48 | 118.51 | 8.59 |
| Asp | 34 | 176.76 | 54.53 | 40.99 | 4.25 | 122.61 | 8.47 |
| Gly | 35 | 174.40 | 45.40 | ND | 3.86 | 109.45 | 8.38 |
| Asp | 36 | ND | 54.41 | 41.17 | ND | 120.74 | 8.32 |
| Glu | 37 | 176.00 | 56.19 | 30.11 | 4.32 | ND | ND |
| Ala | 38 | ND | 50.51 | 17.95 | 4.55 | 127.00 | 8.42 |
| Pro | 39 | 176.39 | 63.49 | 31.88 | ND | ND | ND |
| Ala | 40 | ND | 50.46 | 18.05 | ND | 125.96 | 8.51 |
| Pro | 41 | 177.79 | ND | ND | ND | ND | ND |
| Gly | 42 | ND | ND | ND | 3.62 | 109.42 | 8.58 |
| Lys | 43 | 176.72 | ND | ND | 4.24 | ND | ND |
| Lys | 44 | 176.30 | 56.16 | 32.95 | 4.53 | 123.32 | 8.45 |

| | | | | | | | |
|---|---|---|---|---|---|---|---|
| Ala | 45 | ND | 52.28 | 19.22 | ND | 126.21 | 8.42 |
| Lys | 46 | 176.35 | 56.07 | 33.01 | 4.07 | 124.22 | 8.56 |
| Ala | 47 | 177.29 | 52.36 | 19.23 | ND | 126.23 | 8.46 |
| Ala | 48 | 177.67 | 52.27 | 19.11 | ND | 124.41 | 8.46 |
| Val | 49 | 177.67 | 62.24 | 32.81 | ND | 120.94 | 8.28 |
| Val | 50 | 176.04 | 62.06 | 32.73 | 4.08 | 125.85 | 8.45 |
| Glu | 51 | 176.17 | 56.46 | 30.50 | 4.08 | 126.07 | 8.64 |
| Glu | 52 | ND | 56.39 | ND | 4.32 | 122.72 | 8.55 |
| Glu | 53 | 176.20 | 56.22 | 30.48 | 4.53 | 123.05 | 8.56 |
| Leu | 54 | ND | 53.62 | 41.47 | ND | 125.63 | 8.54 |
| Pro | 55 | 177.14 | 63.23 | 32.06 | ND | ND | ND |
| Ser | 56 | 175.28 | 58.39 | 63.97 | ND | 116.74 | 8.44 |
| Val | 57 | ND | 64.43 | 32.23 | ND | 122.36 | 8.38 |
| Glu | 58 | 177.98 | 58.44 | 29.51 | ND | 122.67 | 8.52 |
| Ala | 59 | 180.86 | 54.56 | 18.45 | 4.25 | 124.27 | 8.35 |
| Lys | 60 | ND | 57.55 | 33.44 | ND | 121.16 | 8.14 |
| Gln | 61 | ND | ND | ND | ND | ND | ND |
| Lys | 62 | 178.52 | 57.55 | 36.17 | 4.07 | 120.19 | 8.26 |
| Glu | 63 | 177.73 | ND | ND | ND | 120.71 | 8.03 |
| Arg | 64 | 179.25 | 59.22 | 30.12 | ND | 120.51 | 8.12 |
| Asp | 65 | 178.26 | 56.67 | 40.04 | 4.32 | 121.46 | 8.68 |
| Ala | 66 | 180.75 | 54.98 | 17.93 | 4.18 | 123.80 | 8.15 |
| Leu | 67 | 179.26 | 57.44 | 41.56 | ND | 119.14 | 8.21 |
| Ala | 68 | ND | 54.96 | 17.89 | 4.18 | 122.71 | 8.06 |
| Lys | 69 | ND | 58.67 | 32.21 | 4.14 | 119.95 | 8.11 |
| Ala | 70 | ND | 54.66 | 17.95 | 4.25 | 122.00 | 8.01 |
| Met | 71 | ND | ND | ND | 4.22 | 119.21 | 8.30 |
| Glu | 72 | 178.49 | 58.91 | 28.29 | 4.07 | 120.16 | 8.28 |
| Glu | 73 | 178.49 | 58.80 | 29.30 | 3.99 | 120.20 | 8.23 |
| Phe | 74 | 178.21 | 60.60 | 39.18 | ND | 121.04 | 8.20 |
| Leu | 75 | 179.73 | 56.90 | 41.63 | 4.02 | 120.03 | 8.43 |
| Ser | 76 | 175.43 | 60.27 | 63.26 | 4.32 | 115.72 | 8.18 |
| Arg | 77 | 176.94 | 56.69 | 30.26 | 4.32 | 120.72 | 7.72 |
| Gly | 78 | 174.73 | 45.34 | ND | 3.46 | 108.39 | 8.06 |
| Gly | 79 | 173.34 | 45.18 | ND | ND | 108.55 | 8.06 |
| Lys | 80 | 176.52 | 55.61 | 33.58 | 4.40 | 120.98 | 8.08 |
| Val | 81 | 175.95 | 62.58 | 33.02 | ND | 122.89 | 8.43 |
| Gln | 82 | 175.70 | 55.63 | 29.56 | 4.40 | 124.81 | 8.60 |
| Glu | 83 | 176.16 | 56.40 | 30.33 | 4.24 | 123.71 | 8.60 |
| Ile | 84 | 176.05 | 60.87 | 38.75 | 4.11 | 122.60 | 8.34 |
| Glu | 85 | ND | 54.24 | 29.77 | 4.59 | 127.40 | 8.61 |
| Pro | 86 | 176.54 | 63.06 | 32.17 | ND | ND | ND |
| Asn | 87 | 174.99 | 53.25 | 38.74 | 4.66 | 119.10 | 8.64 |
| Val | 88 | 176.01 | 62.31 | 32.84 | 4.08 | 121.53 | 8.18 |
| Val | 89 | 175.67 | 62.12 | 32.55 | 4.08 | 125.63 | 8.37 |
| Ala | 90 | 177.08 | 52.20 | 19.44 | 4.25 | 128.97 | 8.48 |
| Asp | 91 | ND | 52.56 | 40.22 | 4.25 | 121.53 | 8.42 |

| | | | | | | | |
|---|---|---|---|---|---|---|---|
| Pro | 92 | ND | ND | ND | ND | ND | ND |
| Pro | 93 | 176.82 | 62.75 | 31.95 | ND | ND | ND |
| Lys | 94 | 176.57 | 55.93 | 33.12 | 4.40 | 122.12 | 8.50 |
| Lys | 95 | ND | 54.29 | ND | 4.59 | 124.86 | 8.56 |
| Pro | 96 | 176.59 | 63.16 | 31.97 | ND | ND | ND |
| Asp | 97 | 176.68 | 54.22 | 41.11 | 4.56 | 121.26 | 8.56 |
| Ser | 98 | 176.68 | 58.66 | 63.74 | 4.36 | 117.22 | 8.42 |
| Lys | 99 | 176.67 | 56.85 | 32.54 | 4.02 | 123.24 | 8.49 |
| Tyr | 100 | 176.56 | 58.22 | 38.51 | 4.53 | 120.61 | 8.22 |
| Gly | 101 | 174.06 | 45.21 | ND | 3.62 | 110.87 | 8.30 |
| Ser | 102 | 174.15 | 58.28 | 63.87 | 4.48 | 115.84 | 8.24 |
| Arg | 103 | ND | 54.04 | 30.23 | ND | 124.12 | 8.39 |
| Pro | 104 | 176.07 | 63.34 | 31.80 | ND | ND | ND |
| Ile | 105 | 181.35 | 63.20 | 39.46 | ND | 125.54 | 7.87 |

**Table 2: Strains used in this study**

Strains for *in vivo* experiments:

| Strain | Strain background genotype | Source |
|---|---|---|
| DKN1625 | *P. aeruginosa UCBPP-PA14* ΔsutA | Babin *et al.* 2016 |
| DKN81 | E. coli MG1655 (F− λ− rph-1) | A gift from Doug Lies, originally from the laboratory of Carol Gross |

*E. coli* strains for cloning and protein expression:

| Strain | Plasmid description | Strain background genotype | Source |
|---|---|---|---|
| DKN1298 | pTNS1 | SM10 | Choi and Schweizer 2006 |
| DKN1299 | pRK2013 | HB101 (F− λ− Δ(gpt-proA)62 leuB6 glnV44(AS) araC14 galK2(Oc) lacY1 Δ(mcrC-mrr) rpsL20(StrR) xylA5 mtl-1 recA13 hsdS20) | Choi and Schweizer 2006 |
| DKN1637 | pMQ30_sutA | DH5α ($F^-$ endA1 glnV44 thi-1 recA1 relA1 gyrA96 deoR nupG Φ80d*lacZ* ΔM15 Δ(*lacZYA-argF*)U169, hsdR17($r_K^-$ $m_K^+$), λ–) | Babin et al. 2016 |
| DKN548 | pMQ72 | F− Δ(argF-lac)169 Φ80dlacZ58(ΔM15) glnV44(AS) λ− rfbC1 gyrA96(NalR) recA1 endA1 spoT1 thi-1 hsdR17 deoR | Shanks *et al*. 2006 |
| DKN1640 | pMQ72_HASutA | Mach1 (ΔrecA1398 endA1 tonA Φ80ΔlacZM15 ΔlacX74 hsdR($r_K^-$ $m_K^+$) ) | Babin *et al.* 2016 |
| DKN1639 | pUC18T-mini-Tn7T-$Gm^R$ $P_{BAD}$:*sutA* | Mach1 (ΔrecA1398 endA1 tonA Φ80ΔlacZM15 ΔlacX74 hsdR($r_K^-$ $m_K^+$) ) | Babin *et al.* 2016 |
| DKN1697 | pQE-80L 6xHis-TEV-SutA | BL21 DE3 ($F^-$ ompT gal dcm lon $hsdS_B$($r_B^-$ $m_B^-$) λ(DE3 [lacI lacUV5-T7 gene 1 ind1 sam7 nin5])) | This study |
| DKN1878 | pQE-80L 6xHis-TEV-SutA 46-101 | BL21 DE3 ($F^-$ ompT gal dcm lon $hsdS_B$($r_B^-$ $m_B^-$) λ(DE3 [lacI lacUV5-T7 gene 1 ind1 sam7 nin5])) | This study |
| DKN1879 | pQE-80L 6xHis-TEV-SutAΔN | BL21 DE3 ($F^-$ ompT gal dcm lon $hsdS_B$($r_B^-$ $m_B^-$) λ(DE3 [lacI lacUV5-T7 gene 1 ind1 sam7 nin5])) | This study |
| DKN1880 | pQE-80L 6xHis-TEV-SutAΔC | BL21 DE3 ($F^-$ ompT gal dcm lon $hsdS_B$($r_B^-$ $m_B^-$) λ(DE3 [lacI lacUV5-T7 gene 1 ind1 sam7 nin5])) | This study |
| DKN1881 | pQE-80L 6xHis-TEV-SutA 6amber | Mach1 (ΔrecA1398 endA1 tonA Φ80ΔlacZM15 ΔlacX74 hsdR($r_K^-$ $m_K^+$) ) | This study |
| DKN1882 | pQE-80L 6xHis-TEV-SutA 11amber | Mach1 (ΔrecA1398 endA1 tonA Φ80ΔlacZM15 ΔlacX74 hsdR($r_K^-$ $m_K^+$) ) | This study |
| DKN1883 | pQE-80L 6xHis-TEV-SutA 22amber | Mach1 (ΔrecA1398 endA1 tonA Φ80ΔlacZM15 ΔlacX74 hsdR($r_K^-$ $m_K^+$) ) | This study |
| DKN1884 | pQE-80L 6xHis-TEV-SutA 54amber | Mach1 (ΔrecA1398 endA1 tonA Φ80ΔlacZM15 ΔlacX74 hsdR($r_K^-$ $m_K^+$) ) | This study |
| DKN1885 | pQE-80L 6xHis-TEV-SutA 61amber | Mach1 (ΔrecA1398 endA1 tonA Φ80ΔlacZM15 ΔlacX74 hsdR($r_K^-$ $m_K^+$) ) | This study |
| DKN1886 | pQE-80L 6xHis-TEV-SutA 74amber | Mach1 (ΔrecA1398 endA1 tonA Φ80ΔlacZM15 ΔlacX74 hsdR($r_K^-$ $m_K^+$) ) | This study |
| DKN1887 | pQE-80L 6xHis-TEV-SutA 84amber | Mach1 (ΔrecA1398 endA1 tonA Φ80ΔlacZM15 ΔlacX74 hsdR($r_K^-$ $m_K^+$) ) | This study |
| DKN1888 | pQE-80L 6xHis-TEV-SutA 89amber | Mach1 (ΔrecA1398 endA1 tonA Φ80ΔlacZM15 ΔlacX74 hsdR($r_K^-$ $m_K^+$) ) | This study |
| DKN1889 | pQE-80L 6xHis-TEV-SutA 100amber | Mach1 (ΔrecA1398 endA1 tonA Φ80ΔlacZM15 ΔlacX74 hsdR($r_K^-$ $m_K^+$) ) | This study |
| DKN1890 | pQE-80L 6xHis-TEV-SutA S2C | BL21 DE3 ($F^-$ ompT gal dcm lon $hsdS_B$($r_B^-$ $m_B^-$) λ(DE3 [lacI lacUV5-T7 gene 1 ind1 sam7 nin5])) | This study |
| DKN1891 | pQE-80L 6xHis-TEV-SutA S32C | BL21 DE3 ($F^-$ ompT gal dcm lon $hsdS_B$($r_B^-$ $m_B^-$) λ(DE3 [lacI lacUV5-T7 gene 1 ind1 sam7 nin5])) | This study |
| DKN1892 | pQE-80L 6xHis-TEV-SutA S98C | BL21 DE3 ($F^-$ ompT gal dcm lon $hsdS_B$($r_B^-$ $m_B^-$) λ(DE3 [lacI lacUV5-T7 gene 1 ind1 sam7 nin5])) | This study |
| DKN1893 | pQE-80L 6xHis-TEV-DksA | BL21 DE3 ($F^-$ ompT gal dcm lon $hsdS_B$($r_B^-$ $m_B^-$) λ(DE3 [lacI lacUV5-T7 gene 1 ind1 sam7 nin5])) | This study |
| DKN1894 | pET15b 6xHis-TEV-RpoS | BL21 DE3 ($F^-$ ompT gal dcm lon $hsdS_B$($r_B^-$ $m_B^-$) λ(DE3 [lacI lacUV5-T7 gene 1 ind1 sam7 nin5])) | This study |
| DKN1895 | pQE-80L 6xHis-TEV-RpoB B1 | E. coli B $F^-$ ompT hsdS($r_B^-$$m_B^-$) $dcm^+$ $Tet^r$ gal λ(DE3) endA Hte [cpn10 cpn60 $Gent^r$] (Arctic Express (DE3) from Agilent) | This study |
| DKN1896 | pQE-80L RpoB355 notag | DH10β ($F^-$ endA1 recA1 galE15 galK16 nupG rpsL ΔlacX74 Φ80lacZΔM15 araD139 Δ(ara,leu)7697 mcrA Δ(mrr-hsdRMS-mcrBC), $λ^-$) | This study |
| DKN1897 | pQE-80L RpoB450 notag | DH10β ($F^-$ endA1 recA1 galE15 galK16 nupG rpsL ΔlacX74 Φ80lacZΔM15 araD139 Δ(ara,leu)7697 mcrA Δ(mrr-hsdRMS-mcrBC), $λ^-$) | This study |
| DKN1898 | pQE-80L RpoB520 notag | DH10β ($F^-$ endA1 recA1 galE15 galK16 nupG rpsL ΔlacX74 Φ80lacZΔM15 araD139 Δ(ara,leu)7697 mcrA Δ(mrr-hsdRMS-mcrBC), $λ^-$) | This study |
| DKN1899 | pQE-80L RpoB626 notag | DH10β ($F^-$ endA1 recA1 galE15 galK16 nupG rpsL ΔlacX74 Φ80lacZΔM15 araD139 Δ(ara,leu)7697 mcrA Δ(mrr-hsdRMS-mcrBC), $λ^-$) | This study |
| DKN1900 | pQE-80L RpoB1062 notag | DH10β ($F^-$ endA1 recA1 galE15 galK16 nupG rpsL ΔlacX74 Φ80lacZΔM15 araD139 Δ(ara,leu)7697 mcrA Δ(mrr-hsdRMS-mcrBC), $λ^-$) | This study |
| DKN1901 | pET15b 6xHis-TEV-RpoD | BL21 DE3 ($F^-$ ompT gal dcm lon $hsdS_B$($r_B^-$ $m_B^-$) λ(DE3 [lacI lacUV5-T7 gene 1 ind1 sam7 nin5])) | This study |
| DKN1902 | pET15b 6xHis-TEV-RpoD Δ171-214 | BL21 DE3 ($F^-$ ompT gal dcm lon $hsdS_B$($r_B^-$ $m_B^-$) λ(DE3 [lacI lacUV5-T7 gene 1 ind1 sam7 nin5])) | This study |
| DKN1903 | pUC18 rrn template | DH10β ($F^-$ endA1 recA1 galE15 galK16 nupG rpsL ΔlacX74 Φ80lacZΔM15 araD139 Δ(ara,leu)7697 mcrA Δ(mrr-hsdRMS-mcrBC), $λ^-$) | This study |

**Table 3: Primers used in this study**

| Name | Sequence | Purpose |
|---|---|---|
| SutATEV F | gagaacctgtacttccagagcATGAGCGAAGAAGAACTGGA | Plasmid mutagenesis |
| SutATEV R | GTGATGGTGATGGTGATGCG | Plasmid mutagenesis |
| SutA46-101 gF | GAACCTGTACTTCCAGAGCATGAAAGCCGCCGTGGTGGAAG | Gibson cloning into plasmid |
| SutA46-101 gR | AGCTCAGCTAATTAAGCTTTCAGCCGTACTTGCTGTCCGG | Gibson cloning into plasmid |
| SutA46-101 pF | CCGGACAGCAAGTACGGCTGAAAGCTTAATTAGCTGAGCT | Gibson cloning into plasmid |
| SutA46-101 pR | CTTCCACCACGGCGGCTTTCATGCTCTGGAAGTACAGGTTC | Gibson cloning into plasmid |
| SutAdN F | ggcaagaaggcgaaagccg | Plasmid mutagenesis |
| SutAdN R | CATgctctggaagtacaggt | Plasmid mutagenesis |
| SutAdC F | TGAAAGCTTAATTAGCTGAGCTTGG | Plasmid mutagenesis |
| SutAdC R | gttgggttcgatctcctgca | Plasmid mutagenesis |
| SutA S2C F | GCGAAGAAGAACTGGAACAG | Plasmid mutagenesis |
| SutA S2C R | ACATGCTCTGGAAGTACAGG | Plasmid mutagenesis |
| SutA S32C F | GCAGTGACGGCGACGAGG | Plasmid mutagenesis |
| SutA S32C R | AGTCCGCTTCGCCGTCGT | Plasmid mutagenesis |
| SutA S98C F | GCAAGTACGGCAGCCGCC | Plasmid mutagenesis |
| SutA S98C R | AGTCCGGCTTCTTCGGCG | Plasmid mutagenesis |
| SutA 6amber F | gaaTAggaacaggacgagctggacg | Plasmid mutagenesis |
| SutA 6amber R | ttcttcgctcatGCTCTGGAAGTAC | Plasmid mutagenesis |
| SutA 11amber F | GtaGGACGGCGCTGACGAG | Plasmid mutagenesis |
| SutA 11amber R | TCGTCCTGTTCCAGTTCTTCTTCgctc | Plasmid mutagenesis |
| SutA 22amber F | GCGAAGAGTAGGCCGCGGCCGACGACGGC | Plasmid mutagenesis |
| SutA 22amber R | CGTCGTCCTCGTCAGCGCCGTCC | Plasmid mutagenesis |
| SutA 54amber F | GAAGAGGAATAGCCCTCGGTCGAAGCCAAG | Plasmid mutagenesis |
| SutA 54amber R | CACCACGGCGGCTTTCGCCTTCTTG | Plasmid mutagenesis |
| SutA 61amber F | gTagaaagagcgtgacgccctcg | Plasmid mutagenesis |
| SutA 61amber R | ttggcttcgaccgagggcagt | Plasmid mutagenesis |
| SutA 74amber F | GGAGGAATAGCTTTCCCGCGGTGGAAAGG | Plasmid mutagenesis |
| SutA 74amber R | ATCGCCTTGGCGAGGGCGTCAC | Plasmid mutagenesis |
| SutA 84amber F | TAGGAACCCAACGTGGTGGCCGA | Plasmid mutagenesis |
| SutA 84amber R | CTCCTGCACCTTTCCACCGCG | Plasmid mutagenesis |
| SutA 89amber F | gTAggccgatccgccgaag | Plasmid mutagenesis |
| SutA 89amber R | acgttgggttcgatctcctgcacc | Plasmid mutagenesis |
| SutA 100amber F | TAGGGCAGCCGCCCCATCTGAAAG | Plasmid mutagenesis |
| SutA 100amber R | CTTGCTGTCCGGCTTCTTCG | Plasmid mutagenesis |
| RpoS gF | CCATCATCATCATCATCACGAGAACCTGTACTTCCAGGGCATGGCACTCAAAAAAGAAGG | Gibson cloning into plasmid |
| RpoS gR | GAGGCCCCAAGGGGTTATGCTAGTCACTGGAACAGCGCGTCAC | Gibson cloning into plasmid |
| RpoS pF | GTGACGCGCTGTTCCAGTGACTAGCATAACCCCTTGGGGCCTC | Gibson cloning into plasmid |
| RpoS pR | CCTTCTTTTTTGAGTGCCATGCCCTGGAAGTACAGGTTCTCGTGATGATGATGATGATGG | Gibson cloning into plasmid |

| | | |
|---|---|---|
| DksA gF | gagaacctgtacttccagagcATGTCCACCAAAGCAAAACA | Gibson cloning into plasmid |
| DksA gR | TCCAAGCTCAGCTAATTAAGCTTTCAGGAGCCGAGTTGCTTCT | Gibson cloning into plasmid |
| DksA pF | AGAAGCAACTCGGCTCCTGAAAGCTTAATTAGCTGAGCTTGGA | Gibson cloning into plasmid |
| DksA pR | TGTTTTGCTTTGGTGGACATgctctggaagtacaggttctc | Gibson cloning into plasmid |
| RpoD gF | CCATCATCATCATCATCACGAGAACCTGTACTTCCAGGGCATGTCCGGAAAAGCGCAACA | Gibson cloning into plasmid |
| RpoD gR | GAGGCCCCAAGGGGTTATGCTAGTCACTCGTCGAGGAAGGAGC | Gibson cloning into plasmid |
| RpoD pF | GCTCCTTCCTCGACGAGTGACTAGCATAACCCCTTGGGGCCTC | Gibson cloning into plasmid |
| RpoD pR | TGTTGCGCTTTTCCGGACATGCCCTGGAAGTACAGGTTCTCGTGATGATGATGATGATGG | Gibson cloning into plasmid |
| RpoD 171-214 F | GGTCCGGATCCGGAAGAA | Plasmid mutagenesis |
| RpoD 171-214 R | GGGATCGATATAGCCGCTGA | Plasmid mutagenesis |
| RpoB B1_us F | GAACCTGTACTTCCAGAGCATGTTGCTGGCCATCCAGCTGGATT | Gibson cloning into plasmid |
| RpoB B1_us R | TCCAGCTGGATGGCCAGCAACATGCTCTGGAAGTACAGGTTCT | Gibson cloning into plasmid |
| RpoB B1_mid F | TCCCAGCTGCACCGTTCCGGTGGTATCGACCACCTGGGCAAC | Gibson cloning into plasmid |
| RpoB B1_mid R | GTTGCCCAGGTGGTCGATACCACCGGAACGGTGCAGCTGGGA | Gibson cloning into plasmid |
| RpoB B1_ds F | TTCGAGCCAGCTGTCGCAGTGAAAGCTTAATTAGCTGAGCT | Gibson cloning into plasmid |
| RpoB B1_ds R | AGCTCAGCTAATTAAGCTTTCACTGCGACAGCTGGCTCGAA | Gibson cloning into plasmid |
| RpoBfrag pF | GACATCGAACTGGAAACCGAATGAAAGCTTAATTAGCTGAGCTTGGA | Gibson cloning into plasmid |
| RpoBfrag pR | CGATCCTCTCATAGTTAATTTCTCCT | Gibson cloning into plasmid |
| RpoB355 F | GGAGAAATTAACTATGAGAGGATCGCTGAAGATCGACAACACCAGC | Gibson cloning into plasmid |
| RpoB450 F | GGAGAAATTAACTATGAGAGGATCGATCGACCACCTGGGCAACCG | Gibson cloning into plasmid |
| RpoB520 F | GGAGAAATTAACTATGAGAGGATCGTTCATGGGCCAGAACAACCCG | Gibson cloning into plasmid |
| RpoB626 F | GGAGAAATTAACTATGAGAGGATCGACCCTCAACGAGAAGGGTCAAC | Gibson cloning into plasmid |
| RpoBfrag R | CAGCTAATTAAGCTTTCATTCGGTTTCCAGTTCGATGTCG | Gibson cloning into plasmid |
| temp_plasmid F | CCGCGCAAGGCACAGTCGAAAGACTGGGCCTTTCGTTTTGCTTAATTAGCTGAGCTTGG | Gibson cloning into plasmid |
| temp_plasmid R | GCTTCGTGTCGAGCCCTTCGCCACGCCCCTCTTTAATACG | Gibson cloning into plasmid |
| rrn_temp F | CGAAGGGCTCGACACGAAGCTTGAAGGTCGGGCGCAAGC | Gibson cloning into plasmid |
| rrn_temp R | TTCGACTGTGCCTTGGCGCGGATTCGACTTGTTAAAGAGCA | Gibson cloning into plasmid |
| generic_temp F | CGAAGGGCTCGACACGAAGC | Template production |
| rrn_temp_short R | CTTGTTAAAGAGCAGTTGGTC | Template production |
| bubble_T | AGGCTTTCGTCTCAACCGAGGCGCGCGTAAGAACAGCAACCTCTCTTTCCGTCAAGCGTTATTTTCGAAAATTTTCTTTT | Template production |
| bubble_NT | AAAAGAAAATTTTCGAAAATAACGCTTGACGGAAAGAGAGGTTGCTGTAGAATGCGCGCGGACGGTTGAGACGAAAGCCT | Template production |
| Cy5 primer ext. | Cy5_GAAAATAACGCTTGACGGAA | Primer extension |
| Cy3 primer ext. | Cy3_AGAGCAGTTGGTCAAGGC | Primer extension |
| rrn RTforRACE | CGAATTCACGAGTGTTAC | 5' RACE |
| rrn RACE_PCR1 F | CGAAGGGCTCGACACGAAGCAAAAAAAAAAAAAAAAAA | 5' RACE |
| rrn RACE_PCR1 R | TTCGACTGTGCCTTGCGCGGTTGCGCTGCTGATAATCTTG | 5' RACE |
| rrn RACE_PCR2 F | CGTATTAAAGAGGGGCGTGGCGAAGGGCTCGACACGAAGC | 5' RACE |
| rrn RACE_PCR2 R | CCAAGCTCAGCTAATTAAGCAAAACGAAAGGCCCAGTCTTTCGACTGTGCCTTGCGCGG | 5' RACE |
| RACE plasmid F | CCGCGCAAGGCACAGTCGAAAGACTGGGCCTTTCGTTTTGCTTAATTAGCTGAGCTTGG | 5' RACE |
| RACE plasmid R | GCTTCGTGTCGAGCCCTTCGCCACGCCCCTCTTTAATACG | 5' RACE |

### SUPPLEMENTAL INFORMATION REFERENCES


1. Shanks RM, Caiazza NC, Hinsa SM, Toutain CM, & O'Toole GA (2006) *Saccharomyces cerevisiae*-based molecular tool kit for manipulation of genes from gram-negative bacteria. *Applied and Environmental Microbiology* 72(7):5027-5036.
2. Gibson DG (2011) Enzymatic assembly of overlapping DNA fragments. *Methods in Enzymology* 498:349-361.
3. Kuznedelov K, *et al.* (2011) The antibacterial threaded-lasso peptide capistruin inhibits bacterial RNA polymerase. *J Mol Biol* 412(5):842-848.
4. Chin JW, Martin AB, King DS, Wang L, & Schultz PG (2002) Addition of a photocrosslinking amino acid to the genetic code of *Escherichia coli*. *Proceedings of the National Academy of Sciences of the United States of America* 99(17):11020-11024.
5. Meares CF, Datwyler SA, Schmidt BD, Owens J, & Ishihama A (2003) Principles and Methods of Affinity Cleavage in Studying Transcription. *Methods in Enzymology* 371:25.
6. Vendrell J & Aviles FC (1986) Complete Amino Acid Analysis of Proteins by Dabsyl Derivatization and Reversed-Phase Liquid Chromatography. *J Chromatogr A* 358:13.
7. Vranken WF, *et al.* (2005) The CCPN data model for NMR spectroscopy: development of a software pipeline. *Proteins* 59(4):687-696.
8. Bahrami A, Assadi AH, Markley JL, & Eghbalnia HR (2009) Probabilistic interaction network of evidence algorithm and its application to complete labeling of peak lists from protein NMR spectroscopy. *PLoS Comput Biol* 5(3):e1000307.
9. Kim DE, Chivian D, & Baker D (2004) Protein structure prediction and analysis using the Robetta server. *Nucleic Acids Research* 32(Web Server issue):W526-531.
10. Chaudhury S, Lyskov S, & Gray JJ (2010) PyRosetta: a script-based interface for implementing molecular modeling algorithms using Rosetta. *Bioinformatics* 26(5):689-691.
11. Bradley P, Misura KM, & Baker D (2005) Toward high-resolution de novo structure prediction for small proteins. *Science* 309(5742):1868-1871.
12. Tamura K, Stecher G, Peterson D, Filipski A, & Kumar S (2013) MEGA6: Molecular Evolutionary Genetics Analysis version 6.0. *Molecular Biology and Evolution* 30(12):2725-2729.
13. Waterhouse AM, Procter JB, Martin DM, Clamp M, & Barton GJ (2009) Jalview Version 2--a multiple sequence alignment editor and analysis workbench. *Bioinformatics* 25(9):1189-1191.
14. Pettersen EF, *et al.* (2004) UCSF Chimera--a visualization system for exploratory research and analysis. *Journal of Computational Chemistry* 25(13):1605-1612.
15. Artsimovitch I & Henkin TM (2009) In vitro approaches to analysis of transcription termination. *Methods* 47(1):37-43.
16. Schindelin J, *et al.* (2012) Fiji: an open-source platform for biological-image analysis. *Nature Methods* 9(7):676-682.
17. Steitz JA & Young RA (1979) Tandem Promoters Direct *E. coli* Ribosomal RNA Synthesis. *Cell* 17:10.
18. Kessner D, Chambers M, Burke R, Agus D, & Mallick P (2008) ProteoWizard: open source software for rapid proteomics tools development. *Bioinformatics* 24(21):2534-2536.
19. Trnka MJ, Baker PR, Robinson PJ, Burlingame AL, & Chalkley RJ (2014) Matching cross-linked peptide spectra: only as good as the worse identification. *Mol Cell Proteomics* 13(2):420-434.
20. Opalka N, *et al.* (2010) Complete structural model of *Escherichia coli* RNA polymerase from a hybrid approach. *PLoS Biology* 8(9).
21. Götze M, *et al.* (2012) StavroX—A Software for Analyzing Crosslinked Products in Protein Interaction Studies. *J. Am. Soc. Mass Spectrom.* 23:12.
22. Adusumilli R & Mallick P (2017) Data Conversion with ProteoWizard msConvert. *Methods in Molecular Biology* 1550:339-368.
23. Wickham H (2016) *ggplot2: Elegant Graphics for Data Analysis* (Springer-Verlag New York, New York).
